# ATP synthase evolution on a cross-braced dated tree of life

**DOI:** 10.1101/2023.04.11.536006

**Authors:** Tara A. Mahendrarajah, Edmund R. R. Moody, Dominik Schrempf, Lénárd L. Szánthó, Nina Dombrowski, Adrián A. Davín, Davide Pisani, Philip C. J. Donoghue, Gergely J. Szöllősi, Tom A. Williams, Anja Spang

## Abstract

The timing of early cellular evolution from the divergence of Archaea and Bacteria to the origin of eukaryotes remains poorly constrained. The ATP synthase complex is thought to have originated prior to the Last Universal Common Ancestor (LUCA) and analyses of ATP synthase genes, together with ribosomes, have played a key role in inferring and rooting the tree of life. Here we reconstruct the evolutionary history of ATP synthases using an expanded sampling of Archaea, Bacteria, and eukaryotes. We developed a phylogenetic cross-bracing approach making use of endosymbioses and ancient gene duplications of the major ATP synthase subunits to infer a highly resolved, dated species tree and establish an absolute timeline for ATP synthase evolution. Our analyses show that the divergence of the ATP synthase into F- and A/V-type lineages, was a very early event in cellular evolution dating back to more than 4Ga potentially predating the diversification of Archaea and Bacteria. Our cross-braced, dated tree of life also provides insight into more recent evolutionary transitions including eukaryogenesis, showing that the eukaryotic nuclear and mitochondrial lineages diverged from their closest archaeal (2.67-2.19Ga) and bacterial (2.58-2.12Ga) relatives at roughly the same time, with the nuclear stem being moderately longer.

## Introduction

The phylogeny and timeline of early cellular evolution, including the age of the last universal common ancestor, the radiations of the archaeal and bacterial domains, and the origin of eukaryotes and their progenitor prokaryote lineages, is poorly constrained^1^. Recent genomics approaches have greatly improved our sampling of natural diversity and uncovered previously unknown microbial lineages which have proven key for our understanding of early cellular evolution^2^. For instance, the Asgard archaea^3–5^ (now referred to as Asgardarchaeota^6^ and Supplementary Table 1), appear to include the closest known sister lineage of the Eukaryota^3, 4, 7, 8^, and have provided support for the evolution of the eukaryotic cell through a symbiosis between at least one asgardarchaeal and one alphaproteobacterial partner^9–15^. The discovery of the symbiotic and genome reduced members of the bacterial Candidate Phyla Radiation (CPR) and the DPANN archaea (named after the first member lineages of this group, the Diapherotrites, Parvarchaeota, Aenigmarchaeota, Nanoarchaeaota and Nanohaloarchaeota)^16–18^, which originally were thought to form early diverging branches on each side of the root of the TOL^2^, might be important for our understanding of the deep split separating Archaea and Bacteria^19^. Though, more recent phylogenomic analyses suggest that the CPR instead form a sister group to the Chloroflexota within the Terrabacteria^19–23^. Irrespective, timing molecular evolution is challenging because the rate of molecular evolution has varied substantially through time^19, 24–27^ and, with few fossil calibrations (e.g. maximum age constraints), clock models struggle to capture this rate variation. This has led to uncertain and in some cases implausible estimates for the age of key nodes in the tree of life, such as the last universal common ancestor (LUCA)^19, 27^. Additional sources of temporal information beyond fossil and geochemical calibrations are therefore crucial.

The ATP synthase is a protein complex central to energy conservation through the synthesis and hydrolysis of ATP^28, 29^. It is a useful marker to address key evolutionary transitions due to the presence of this enzyme across all domains of life^24, 28, 30–36^. The ATP synthase family is classified into the F-, A-, and V-type ATP synthases^29, 32, 33^ based on their taxonomic affiliation, function, and cellular localization^29, 33, 37^. F-type ATP synthases are ubiquitous across bacteria and eukaryotes and localize to cellular, mitochondrial, and plastid membranes^38^. In line with this, eukaryotic F-type ATP synthases are hypothesized to be derived from the bacterial ancestors of these organelles^32, 34, 39^. The A-type ATP synthase ^33, 37, 40^, found primarily in Archaea, belongs to a larger family of A/V-type ATP synthases ^37^, which also include eukaryotic complexes found in vacuoles^31–33, 37, 39–41^. The F- and A/V-type ATP synthases share a common foundational architecture consisting of a soluble cytoplasmic component (R1) connected to an insoluble membrane component (R0). The hexameric headpiece of the R1 complex contains three copies each of a catalytic (*c*) and non-catalytic (*nc*) subunit and is the site of ATP synthesis and hydrolysis. The catalytic and non-catalytic subunits comprising the soluble hetero-hexameric R1 component, are paralogous to each other and arose prior to LUCA through an ancient duplication of a RecA family protein (P-loop NTPase) followed by the loss of the catalytic function in one subunit^28, 31–34, 37, 39, 42^. Due to this ancient gene duplication, each paralog can act as an outgroup to the other, providing a way to root the tree of life^43^. Since the duplication occurred before the divergence between Archaea and Bacteria, speciation events during the subsequent history of life appear at least twice in the ATP synthase gene tree. This circumstance has been used to improve date estimates for eukaryotic evolution by “cross-bracing” (constraining to the same unknown age) equivalent speciation nodes in the gene tree^24^, which propagates the limited fossil evidence across the tree. In principle, the same approach might be applied to the universal TOL in which eukaryotes appear multiple times due to the mitochondrial and plastid endosymbioses. Cross-bracing a ribosomal species tree could help to avoid difficulties arising from HGT events during ATP synthase evolution^31, 32, 39, 44^, and the limited resolving power of single gene trees.

To improve our understanding of the evolutionary history of ATP synthases and cellular evolution, we used an updated taxon sampling set, gene tree-species tree reconciliation methods^45–47^, ancestral sequence reconstruction (ASR)^48^, and novel molecular dating approaches implementing cross-bracing^24, 49, 50^. We assembled a set of ribosomal marker proteins that includes three distinct clades of eukaryotic homologs derived from archaeal, alphaproteobacterial, and cyanobacterial ancestors. Gene duplications (ATP synthase) and endosymbioses (ribosomal marker genes and ATP synthase), meant cross-bracing could be applied to both datasets. Our analyses confirm the split of the catalytic and non-catalytic ATP synthase subunits prior to LUCA and reveal the prevalence and early evolution of A/V-type ATP synthases in Bacteria. Our dating analyses allowed us to establish absolute time estimates for LUCA, the Last Bacterial Common Ancestor (LBCA), and the Last Archaeal Common Ancestor (LACA), and to link early cellular evolution with the origin of the head component of ATP synthases, which has diversified earlier than previously assumed. Finally, our analyses improve time estimates for the origin of eukaryotes from its prokaryotic ancestors and thereby inform on eukaryogenesis.

## Results

### Distribution of ATP synthases across Bacteria, Archaea and Eukaryotes

We analysed the distribution of ATP synthase genes across our reference dataset of 800 Archaea, Bacteria, and eukaryotes (Figure 1-2, Supplementary Figure 2, Supplementary Tables 1-4). In agreement with previous work^28, 31–35, 37, 39^, our results indicate a partitioning of the F- and A/V-Type ATP synthases by domain, with Archaea and Bacteria containing primarily A/V-Type and F-Type subunits, respectively, and eukaryotes harboring complexes of both types (Figure 1, Supplementary Figure 2). However, in Bacteria the pattern is more complex than often assumed^32, 34, 51^. Consistent with emerging evidence that several Bacteria contain A/V-type ATP synthases^34^, we found that 46% (23/50) of bacterial phylum-level lineages encode genes for A/V-Type ATP synthases in conjunction with (n=19), or to the exclusion of (n=4), a bacterial F-Type ATP synthase (Figure 1, Supplementary Figure 2, Supplementary Table 4). Conversely, only three members of a single archaeal lineage, the Methanosarcinales, contain F-Type ATP synthases in addition to their A/V-Type complex (Figure 1, Supplementary Figure 2, Supplementary Table 4), as observed previously^52–54^.

**Figure 1:**
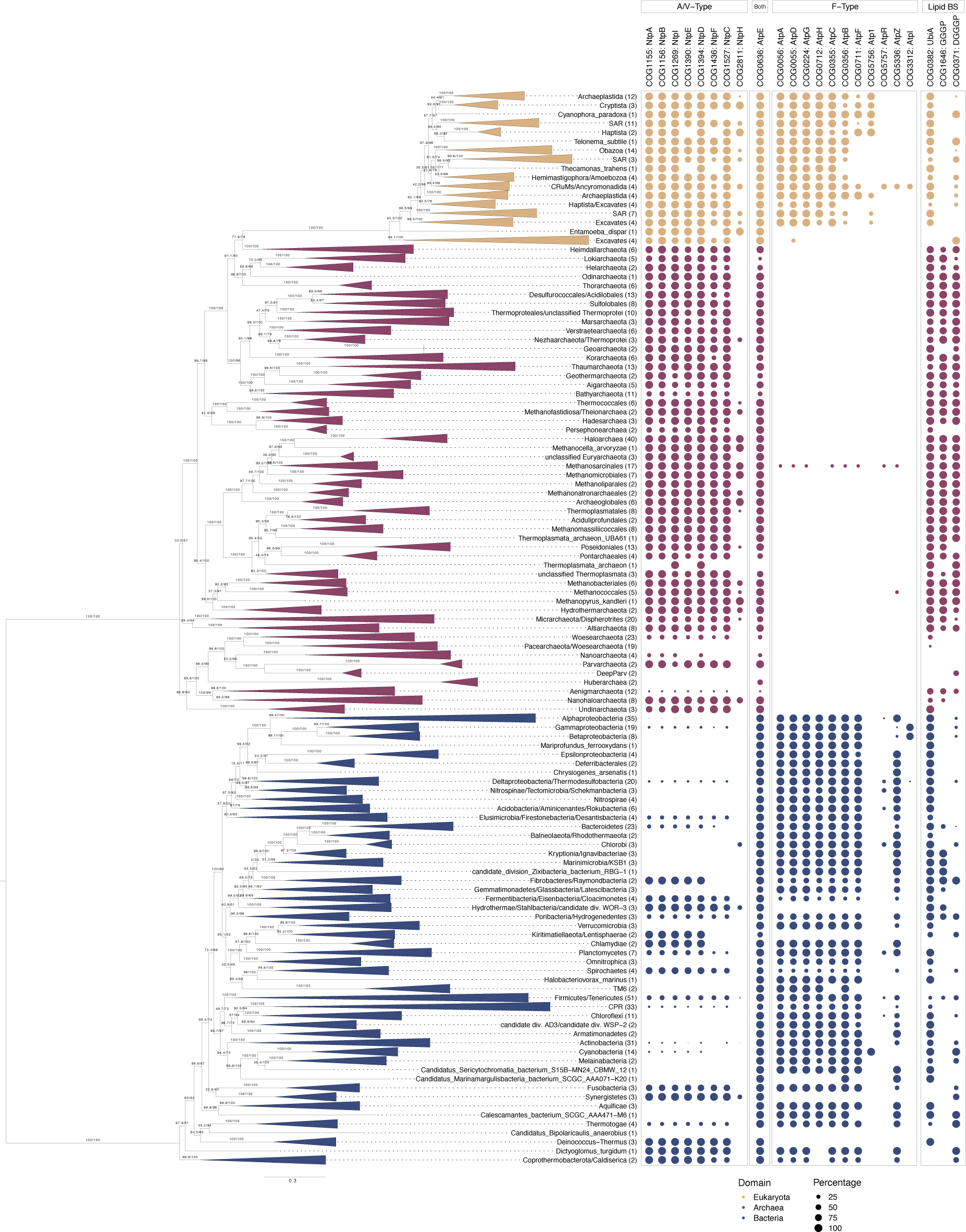
Distribution of COG families representing the F- and A/V-type ATP synthase subunits and select lipid biosynthesis genes across the TOL. COG families corresponding to the ATP synthase subunits and lipid biosynthesis genes (see Methods for selection of COG families, Supplementary Table 3) are represented as a percentage presence by phylogenetic cluster, consistent with collapsed taxonomic clades in the maximum-likelihood concatenated species tree. The concatenated alignment contains 780 taxa and was trimmed with BMGE v1.12 (settings: -m BLOSUM30 -h 0.55) ^101^ to remove poorly-aligning positions (final alignment length = 3367 amino acids). The maximum likelihood tree was inferred using IQ-TREE2 v2.1.2 with the LG+C20+R+F model with ultrafast bootstrap approximation (left) and SH-like approximate likelihood (right), each with 1000 replicates ^102–104^. The scale bar corresponds to the expected number of substitutions per site. Color code: archaea = red, bacteria = blue, eukaryotes = yellow.

**Figure 2:**
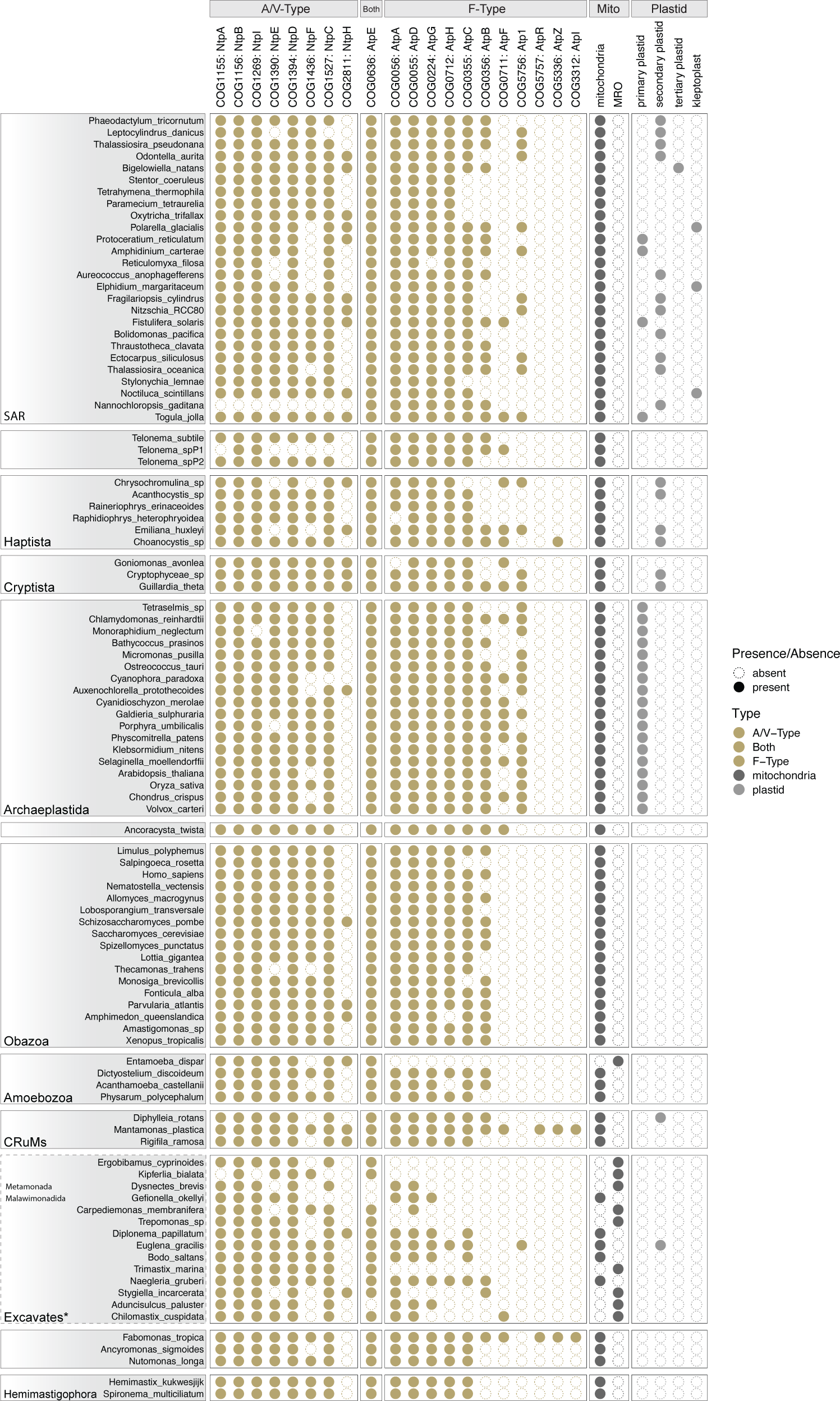
Occurrence of COG families representing the F- and A/V-type ATP synthase subunits and the presence/absence of key metabolic organelles across the 100 sampled eukaryotes. COG families representing the ATP synthase subunits (see Methods for selection of COG families, Supplementary Table 3) are presented as binary presence-absence counts per taxon. The relationships among eukaryotic supergroups is consistent with Burki, 2020^105^. Dashed lines represent groups with greater uncertainty. Mito = mitochondria and mitochondrion-related organelles (MROs). Plastid = primary-, secondary-, and tertiary-plastid, and kleptoplast. See Supplementary Table 5 for additional information on organelle distribution.

Despite the core role of ATP synthase in energy conservation, some prokaryotes, including members of the DPANN archaea^55–57^, lack functional homologs (Figure 1, Supplementary Figure 2, Supplementary Table 4). These losses are present across related lineages, suggesting a genuine ancestral loss, rather than metagenome assembled genome (MAG) incompleteness. Other DPANN lineages, such as Nanohaloarchaeota, may have inherited their ATP synthase from a DPANN ancestor (Supplementary Figure 6) or acquired ATP synthase genes from symbiotic partners (Supplementary Figures 7-10, Discussion)^44, 58^. Several Bathyarchaeota lack ATP synthase complexes (Figure 1, Supplementary Figure 2, Supplementary Table 4), consistent with genome-based predictions indicating that Bathyarchaeota BA1 and BA2 lack an ATP synthase complex, possibly producing ATP through substrate-level phosphorylation using a putative ATP-forming acetyl-CoA synthetase^59, 60^. We found that 60.6% (20/33) of CPR (phyla) encode F-type ATP synthases, suggesting inheritance from a common ancestor with Chloroflexi (Figure 1, Supplementary Figure 2, Supplementary Table 4). Conversely, 30.3% (10/33) of the sampled CPR have lost genes that would enable the formation of canonical ATP synthase complex (Supplementary Figure 2, Supplementary Table 4). Interestingly, three members of the CPR in our dataset lack an F-type ATP synthase but have a near-complete or complete A/V-type complex (Figure 1, Supplementary Figure 2), indicating a recent acquisition of the A/V-type complex via HGT potentially by members of the Synergistetes (Supplementary Figures 2, 6-10).

All eukaryotic lineages contained core functional subunits of both F- and A/V-type ATP synthases, with the exception of 10/100 analyzed representatives, including *Entamoeba dispar* and certain Excavates (Figure 2, Supplementary Table 5). This is consistent with the energy metabolism of these anaerobes whose mitochondrion-related organelles have lost components of the aerobic electron transport chain^61, 62^ (Figure 2). We observed that 83% (14/17) of Archaeplastida encode genes for Atp1, the F-type alpha subunit of the chloroplast (COG5756, Figure 2, Supplementary Table 5), while this gene is lacking in species without a plastid (Figure 2), consistent with the existence of a second F-type ATP synthase of endosymbiotic origin in Chloroplasts. However, several eukaryotes (15%) lack an Atp1 homolog despite harboring a plastid (Figure 2), indicating subsequent loss of Atp1 in some photosynthetic eukaryotes.

### Evolutionary history of soluble ATP synthase subunits

Phylogenetic analyses including all catalytic (*c*) and non-catalytic (*nc*) subunits of the soluble head component of the F- and A/V-type ATP synthase (Supplementary Figure 1), the F1 beta and A1/V1A and the F1 alpha and A1/V1B, respectively (hereafter referred to as *c*F1, *c*A1V1, *nc*F1, *nc*A1V1), revealed four clades corresponding to each of the four protein families (Figure 3A, Supplementary Figure 10). Based on the gene family tree and the observation that all organisms encoding an ATP synthase possess catalytic (*c*F1 and *c*A1V1) (Figure 3A, Supplementary Figures 4, 6, 8-10) and non-catalytic (*nc*F1 and *nc*A1V1) (Figure 3A, Supplementary Figures 5, 7-10) subunits, our analysis agrees with the consensus view^30, 31, 34, 39^, that the deepest split lies between those families (Figure 3A, Supplementary Figure 10)^30, 31, 34, 39^. Our results suggest an early divergence of the functional capacities of each subunit followed by subsequent bifurcations into F- and A/V-type complexes (Figure 3A, Supplementary Figure 10). The observed deep splits within each of the catalytic and non-catalytic subunits of the F- and A/V-type complexes, respectively, have been hypothesized to coincide with the speciation of Archaea and Bacteria^30, 31, 63^ (Figure 3A).

**Figure 3:**
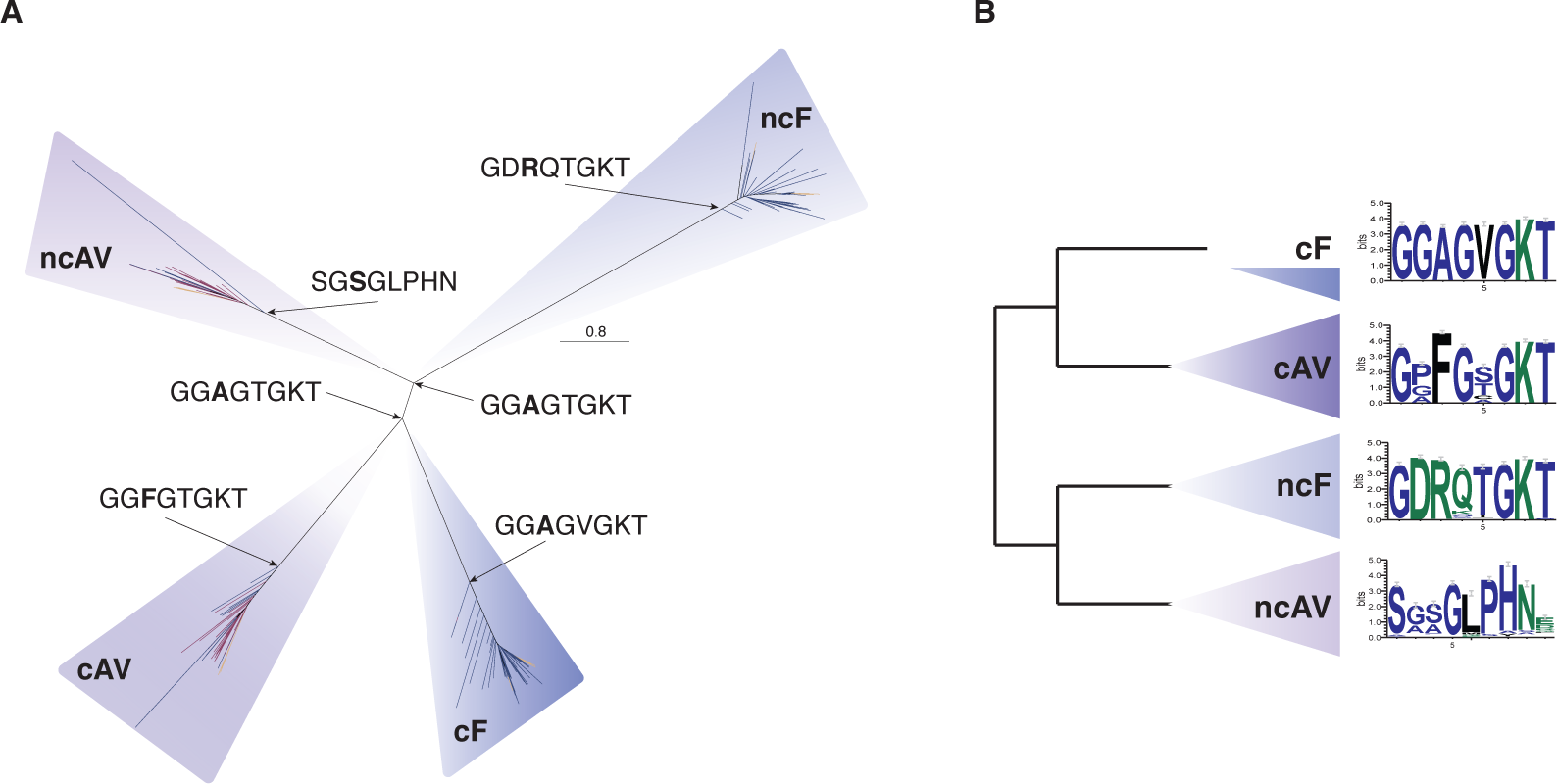
Maximum likelihood tree of all ATP synthase headpiece subunits identified in sampled Archaea (red), Bacteria (blue), and Eukaryotes (yellow). (A) Homologs corresponding to each subunit form monophyletic clusters for each protein family. Catalytic subunits (*c*F1 and *c*A1V1) and non-catalytic subunits (*nc*F1 and *nc*A1V1) cluster together on either side of the root. The alignment contains 1520 sequences and was trimmed with BMGE v1.12 (settings: -m BLOSUM30 -h 0.55) ^101^ (alignment length = 350 amino acids). The maximum likelihood tree was inferred using IQ-TREE2 v2.1.2 with the LG+C50+R+F model, selected using the best-fitting model (chosen by BIC) ^102, 104, 106^. The scale bar corresponds to the expected number of substitutions per site. The Walker-A motif from ancestrally reconstructed sequences ^104^ are shown at their respective nodes. (B) Conserved protein motifs for each subunit derived from the same alignment.

Confidently determining which of these four head-forming subunits was present in LACA, LBCA, and LUCA based on gene tree inspection is challenging. For instance, the identification of A/V-type ATP synthases in many Bacteria (Figure 1, Supplementary Figure 2) and the recent inference of the presence of components of both F- and A/V-Type ATP synthases in the genome of LBCA^21^, challenge a late horizontal acquisition of the A/V-type ATP synthase by Bacteria. To evaluate these hypotheses within a statistical framework, we used the Amalgamated likelihood estimation (ALE) probabilistic approach^46^ to reconcile gene trees for each of ATP synthase subunits with the species tree as a whole, using distinct data treatments (Methods, Supplementary Table 6). These analyses agreed with manual inspection of the gene trees, suggesting that the *c*A1V1 and *nc*A1V1 subunits were present in LACA (presence probability, PPs = 0.99-1)^64^ and the *c*F1 and *nc*F1 subunits were present in LBCA (PPs = 0.99-1)^21^. We recovered support for the presence of the *c*A1V1 (PPs = 0.64-1) subunit in LBCA, as has been suggested recently^21^ while the presence of the *nc*A1V1 subunit in LBCA was supported only in trees inferred using LG+C20+R+F (C20: PPs = 0.99-1, C60: PPs = 0.21-0.28, Supplementary Table 6).

The *nc*F1 (PPs = 0.79-1), *nc*A1V1 (PPs = 0.99-1), *c*F1 (PPs = 1), and *c*A1V1 (PPs = 0.99-1) gene families were estimated to having been present in LUCA, suggesting a putative pre-LUCA duplication of both the catalytic and non-catalytic subunits into the F- and A/V-type lineages (Supplementary Table 6, Figure 4). However, deep branches in the gene trees are susceptible to systematic error, and distinguishing ancestral presence from early horizontal acquisition is difficult^21^. Nonetheless, the widespread presence of genes encoding A/V-type subunits in modern Bacteria (Figure 1, Supplementary Figure 2, Supplementary Table 4) suggests that these genes were acquired early in bacterial evolution.

**Figure 4:**
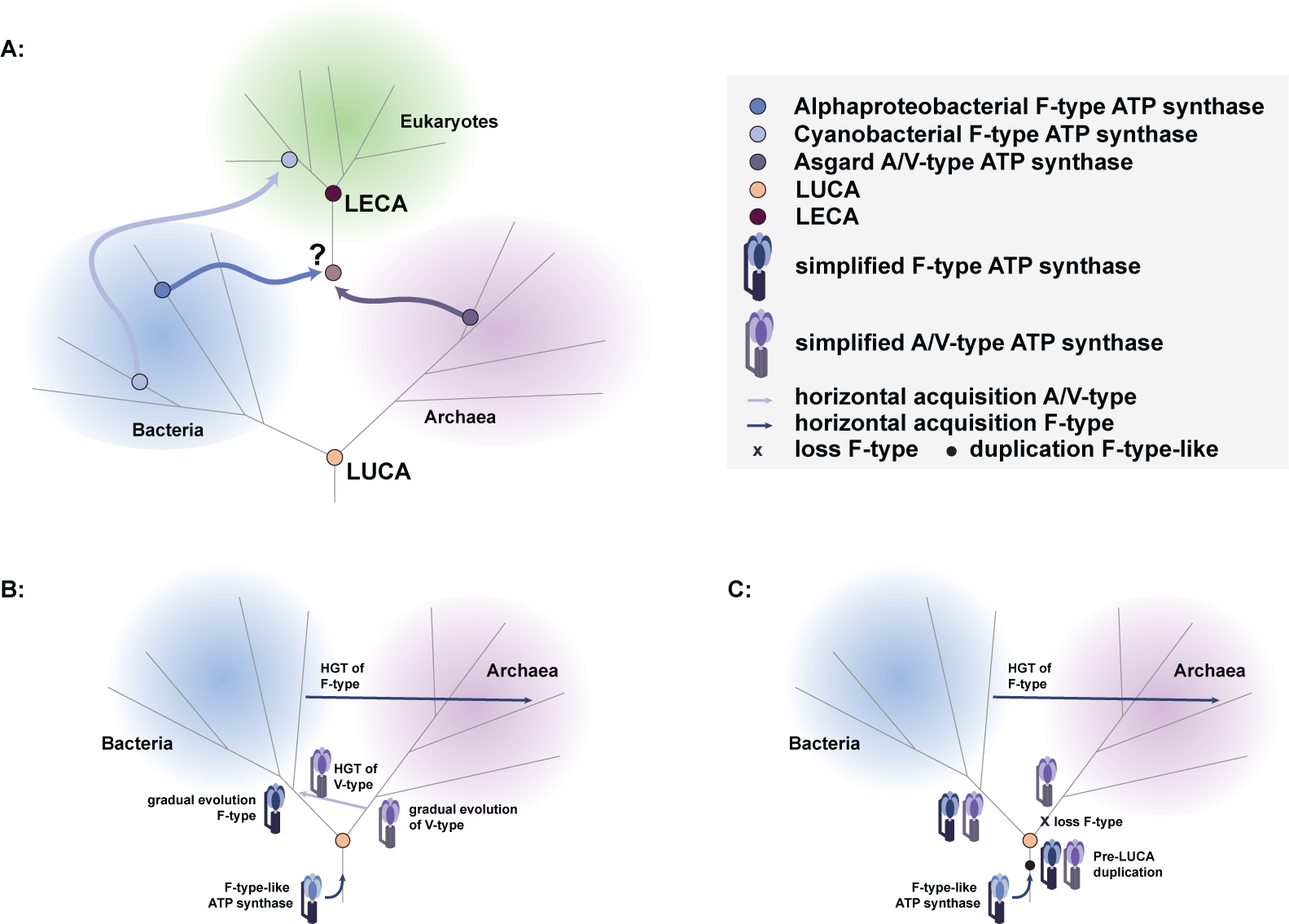
ATP synthase evolutionary scenarios. (A) Overview of possible ancestral ATP synthase acquisition in LECA from the putative prokaryotic hosts; the A/V-type derived from the archaeal host, and F-type ATP synthases derived from bacterial endosymbionts. (B) Evolutionary proposal supporting an F-type-like ancestral ATP synthase present pre-LUCA with subsequent divergence consistent with the split between Bacteria and Archaea and early transfers of A/V-type ATP synthases into the bacterial stem, and late HGT of F-type ATP synthases to Archaea. (C) Evolutionary proposal supporting an F-type-like ancestral ATP synthase and pre-LUCA duplication and divergence of at least the head components of the F- and A/V-type subunits with subsequent loss of the F-type components along the archaeal stem.

Presence of all four subunits in Bacteria is consistent with ideas for a root of the universal tree within Bacteria^65–67^. However, our analyses estimated significantly lower likelihoods (C60, AU = 0.00009; C20, AU = 0.0002) (Supplementary Table 6) for ATP synthase subunits on a re-rooted species tree including eukaryotes in which the root was placed between the Gracilicutes and the rest of life, consistent with previous placements of the bacterial root^20, 21^. When eukaryotes were excluded, the within-Bacteria root also had a lower likelihood, though not significant (C60, AU = 0.093; C20, AU = 0.547) (Supplementary Table 6). These results agree with the consensus root between Archaea and Bacteria.

To investigate the key motifs characterizing the catalytic and non-catalytic subunits of the F- and A/V-type ATP synthases throughout the TOL, we examined conserved protein motifs in extant taxa and conducted ASR to predict amino acid composition of the ancestral sequences^48, 68^. We focused on the sequence identity of the Walker-A motif (Supplementary Discussion), which has an amino acid composition of *GXXXXGKT*^42^. This region comprises the primary “P-loop” domain responsible for binding phosphate during ATP synthesis/hydrolysis and is highly conserved across phosphate-binding proteins and fundamental to the activity of the ATP synthase^69, 70^. Our analyses of the Walker-A motifs across the *nc*F1, *c*F1, and *c*A1V1 subunits revealed a conserved Walker-A motif with variation in positions 2-5 (Figure 3B). However, the *nc*A1V1 subunit lacks a recognisable Walker-A motif and instead contains a *SGSGLPHN* motif in the same position as the expected Walker-A motif with the phosphate binding properties of this motif being unknown (Figure 3B)^71^.

We performed ASR on the alignment of the unrooted combined phylogeny (Figure 3A, Supplementary Table 7) to determine the ancestral sequence at the root of each of the four subunits (*nc*F1: Node123; *c*F1: Node126; *c*A1V1: Node516; and *nc*A1V1: Node883) as well as the root of the catalytic versus non-catalytic subunits (Figure 3A) (*nc*F1 and *nc*A1V1: Node124; *c*F1 and *c*A1V1: Node125). Consistent with our observations of the conserved extant motifs, we found Walker-A motifs in the ancestrally reconstructed sequences for the *nc*F1, *c*F1, and *c*A1V1 families and the alternative motif (*SGSGLPHN*) for the *nc*A1V1 family (Figure 3A). The alanine (A) and phenylalanine (F) dichotomy in the third position of the *nc*F1, *c*F1 and *c*AV ancestors is consistent with previous findings distinguishing F- and A/V-type ATP synthase catalytic binding loops, respectively (Figure 3A)^40^. A motif pattern of *GGAGTGKT* was inferred for both the ancestor of the catalytic and non-catalytic subunits (Figure 3A), compatible with a previously proposed scenario in which the progenitor ATP synthase was suggested to have contained 6 catalytic sites similar to the *c*F1, indicating a pre-LUCA F-type-like ancestor^33^. Our results imply that the *nc*A1V1 subunit lost its Walker-A motif after divergence from the other subunits, though the functional consequence of the degenerated binding loop in the *nc*A1V1 subunit is unknown. Interestingly, while ancestral sequences inferred for the *nc*F1, *nc*A1V1, *c*F1 and *c*A1V1 are most similar to those in extant representatives of each of those families, the sequences inferred for each ancestor of the *nc* and *c* families were both most similar to extant members of the *c*F1 subunits from F-type ATP synthases (Figure 3, Supplementary Table 7). Taken together, this may indicate that the ancestral head component of the ATP synthase was more similar to the R1 complex of F-type ATP synthase and is consistent with the hypothesis that they evolved by duplication from a catalytic ancestor belonging to the “P-loop” NTPases^24, 28, 30, 33, 36^.

### The origins of ATP synthases in eukaryotes

In agreement with symbiogenetic models for the origin of the eukaryotic cell^9–15^, our ATP synthase phylogenies suggest that eukaryotes have inherited A/V- and F-type ATP synthases from their archaeal and bacterial ancestors (Figure 3A, Supplementary Figures 4-10)^11, 31–33, 39^. We observed that the phylogenetic relationships inferred from the non-catalytic subunits (both *nc*F1 and *nc*A1V1) were more consistent with established views on the species-level relationships than those inferred from the catalytic subunits (Supplementary Figures 4 and 6, Supplementary Table 8). Specifically, the relationship between Asgard archaea and eukaryotes was evident in phylogenies of the *nc*A1V1 subunit (Supplementary Figures 6-10, Supplementary Table 8), with the strongest bootstrap support being 95.8/95 (Supplementary Figure 7, Supplementary Table 8). In phylogenies for the catalytic subunits the position of eukaryotes was mostly unresolved or placed elsewhere within prokaryotes (Supplementary Discussion, Supplementary Figures 6-10, and Supplementary Table 8). The origin of eukaryotic F-type sequences from Alphaproteobacteria and Cyanobacteria was consistently recovered across a range of analyses including both Baysian and ML inferences (Supplementary Discussion, Supplementary Figures 4-5, Supplementary Figure 11, Supplementary Table 8). Within the F-type subunits, the *nc*F1 phylogenies placed the sequences of eukaryotic plastids sister to *Gloeomargarita lithophora*, the closest living relative of the plastid^72^, while the *c*F1 phylogeny grouped plastids together with most Cyanobacteria (Supplementary Figure 11). Differences in selective constraints between the catalytic and non-catalytic subunits might render the non-catalytic subunit more amenable to phylogenetic analysis.

### Dating the species tree and establishing an absolute timeline for ATP synthase evolution

To establish a timeframe for the evolution of the ATP synthase, we built on the pioneering strategy of Shih and Matzke (2013) to brace equivalent speciation nodes in both the ATP synthase phylogeny and a universal species tree^24^. We took advantage of the greatly expanded sampling of organisms sequenced since the previous study (1520 total *nc* and *c* ATP synthase subunit sequences included in this study versus 149 total sequences in Shih and Matzke, 2013) and applied more fossil calibrations (ATP synthase gene tree n=10, species tree n=12 versus n=7 in Shih and Matzke, 2013), based on a reappraisal of the early fossil record (Supplementary Discussion, Supplementary Tables 9). Importantly, we developed a new molecular dating software (McmcDate) which implements both cross-bracing (two nodes are constrained to be the same age) as well as relative constraints (one node is constrained to always be younger than another node^50, 73^ (Supplementary Discussion, Supplementary Table 9, Supplementary Data Files).

Molecular dating analyses revealed that bracing the nuclear, mitochondrial, and plastid eukaryotic clades had a significant overall impact (Z-test statistic was -233.0 with a p-value of 0.0) on inferred rates of evolution, which are 16.5% higher overall than in the non-braced analysis (2.6e-4 average number of substitutions per million years and site, for ribosomal protein tree; Figure 5A, Figure 5C, Supplementary Figures 12-17, Supplementary Table 10). As a result, age ranges (measured as 95% HPD: the boundaries of the central 95% highest posterior densities of the distributions on ages) are modestly younger in the braced analysis, though similar overall (Supplementary Table 10). We inferred that LUCA lived 4.52-4.32Ga (Figure 5A, Supplementary Figure 16). Of the two prokaryotic domains, LBCA was inferred to be older (4.49-4.05Ga) than LACA (3.95-3.37Ga) indicating higher extinction or lower sampling rates for total group Archaea (Figure 5A, Supplementary Figure 16). The divergence between eukaryotes and their closest archaeal and bacterial relatives was suggested to have occurred during the Palaeoproterozoic. Specifically, our analyses suggest that eukaryotes diverged from their closest known asgardarchaeal relatives 2.67-2.19Ga (Hodarchaeota + Eukaryotes), and from Alphaproteobacteria 2.58-2.12Ga. Plastids diverged from free-living Cyanobacteria 2.14-1.73Ga (Figure 5A, Figure 5C, Supplementary Figure 16, Supplementary Table 10). We inferred LECA to have lived between 1.93-1.84Ga (Supplementary Figure 16, Supplementary Table 10). These revised ages for key nodes in the species tree provide a timeline to study ATP synthase diversification in the context of cellular evolution: the split between the catalytic and non-catalytic ATP synthase subunits (4.52-4.46Ga) likely predates, or at the latest was contemporary with, LUCA (4.52-4.32Ga), while the divergence into F1- and A1V1-types within the catalytic (4.52-4.38Ga) and noncatalytic (4.52-4.42Ga) clades overlaps in time with LUCA (4.52-4.32Ga) and LBCA (4.49-4.05Ga) but predates LACA (3.95-3.37Ga) by more than 0.5Ga (Figure 5B, Supplementary Figures 18-19, Supplementary Table 10). An early origin of the A/V-type ATP synthases is a prerequisite for their presence in LBCA, and - if the split between F- and A/V-types corresponds to the speciation of Archaea and Bacteria - an older age for the A1/V1 clade compared to crown Archaea (LACA) might hint at a sampling or extinction “bottleneck” on the stem lineage leading to extant Archaea (Figure 4B). Alternatively, as the inferred ages of the divergences into F1- and A1V1-types overlap with LUCA’s age, the dating analysis is consistent with a scenario in which the head components of the F- and A/V-type ATP synthases were already present prior to the divergence of Archaea and Bacteria (Figure 4C) in agreement with our reconciliations (see above and discussion).

**Figure 5:**
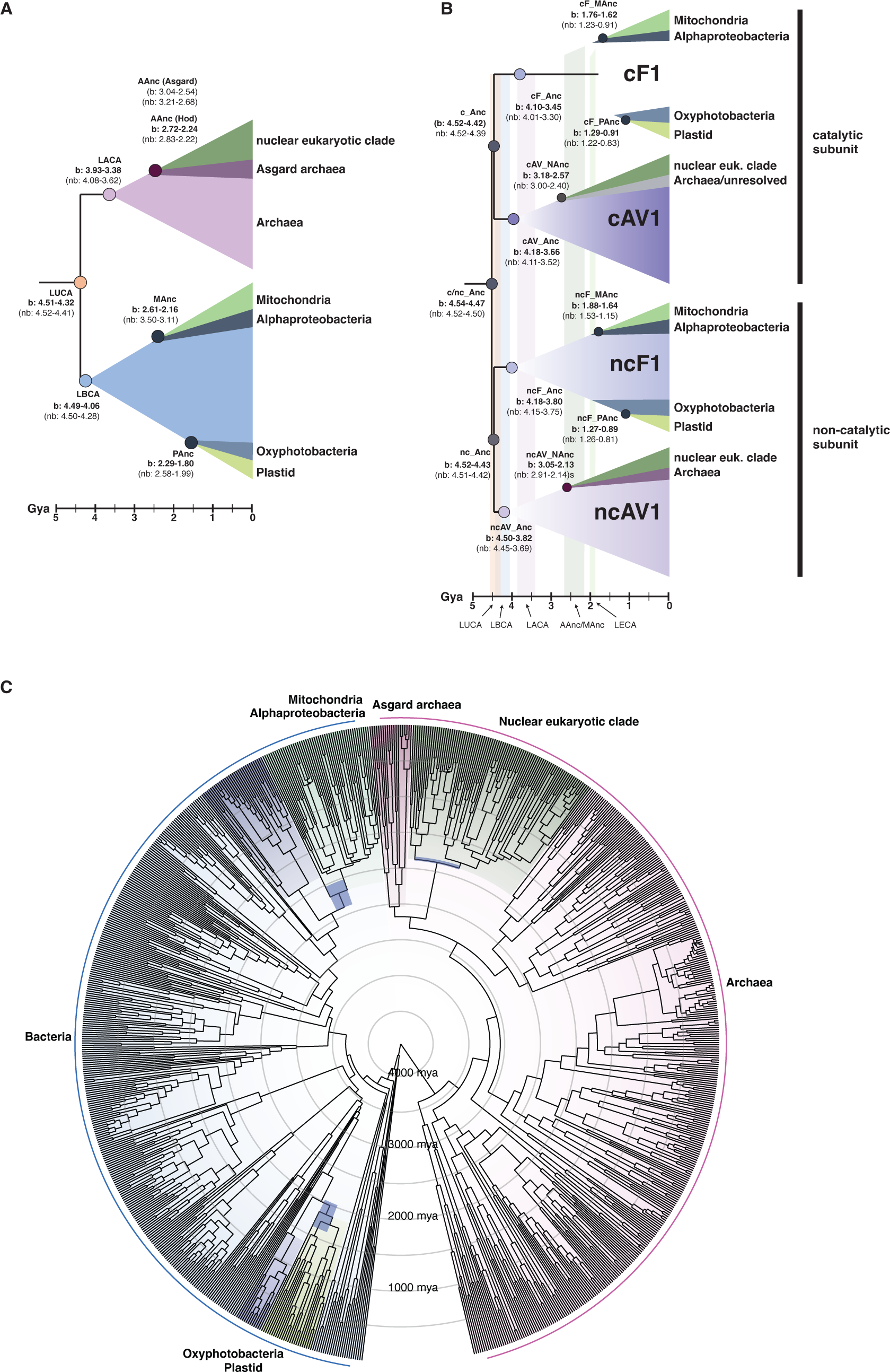
Timing of cellular evolution across the TOL based on a cross-braced dated ribosomal species tree and ATP synthase gene tree. (A) Suggested timing of key evolutionary events based on a schematic ribosomal species tree. (B) Suggested timing of key evolutionary events based on a schematic ATP synthase gene tree. (C) Dated cross-braced ribosomal species tree (Edited2, see Methods) including nuclear, mitochondrial, and plastid eukaryotic homologs. See Methods for inference of the maximum likelihood concatenated ribosomal species phylogeny and constraints (Edited2). The alignment contained 863 sequences and was trimmed with TRIMAL ^107^ (alignment length = 2133 amino acids), and the maximum likelihood phylogeny was inferred using IQ-TREE2 v2.1.2 the LG+C60+R+F model with ultrafast bootstrap approximation (left) and SH-like approximate likelihood (right), each with 1000 replicates ^102–104^.

## Discussion

Our analyses confirm that the A/V-type ATP synthase was present in LACA^64^ and the F-type ATP synthase in LBCA^19–22^. They also revealed that A/V-type ATP synthases are broadly distributed in Bacteria and might have already been present in LBCA (Figure 1, Figure 3A, Supplementary Figure 2, Supplementary Figures 6-8, Supplementary Figure 10). Previous analyses suggested that the acquisition of A/V-type ATP synthases in Bacteria occurred via HGT from hyperthermophilic Archaea^32, 34, 51^, contradicting the observation that many mesophilic Bacteria also contain A/V-type ATP synthases (Figure 1, Figure 3A, Supplementary Figure 2). In contrast, only three archaeal genomes (all within the genus *Methanosarcina*) appear to encode F-type ATP synthases (Figure 1, Figure 3A, Supplementary Figure 2, Supplementary Figures 4-5) belonging to a family of ATP synthases known as N-ATPases, a distinct horizontally-acquired F-type ATP synthase which exists in addition to a native F- or A/V-type ATP synthase in Bacteria and Archaea^54^. Experimental studies revealed that the N-ATPase of *M. acetivorans* is not required for growth^53, 54^; therefore the function of these F-type ATP synthases is debated.

ATP synthase evolution in Bacteria seems to be driven by frequent transfers from Archaea to Bacteria; the alternative possibility is that A/V-type ATP synthases were already present in LBCA or LUCA, in which case they must have been lost in many bacterial lineages. The latter possibility is in line with a scenario in which transfers from Bacteria to Archaea have been more common during evolution^74^, but requires a loss of the F-type ATP synthase along the branch leading to Archaea (Figure 4C). In agreement with this, the duplication giving rise to the catalytic and non-catalytic subunits (4.52-4.42Ga 95% HPD, Supplementary Figures 18-19, Supplementary Table 10) and the divergence into F- and A/V-lineages within the catalytic and non-catalytic clades (4.52-4.38Ga 95% HPD, Supplementary Figures 18-19, Supplementary Table 10) were inferred to have occurred very early in the history of cellular life, prior to or overlapping with our age for LUCA (4.52-4.32Ga, Figure 5A, Figure 5C, Supplementary Figure 16, Supplementary Table 10). This may seem at odds with previous inferences^10, 28, 31, 75^ and suggestions of deep divergences between Archaea and Bacteria coinciding with distinct informational processing machinery^19^, ATP synthases, and membrane lipids^35, 76^. However, the ‘lipid divide’^77^ appears less pronounced than assumed previously^78, 79^ and LUCA may have had the mevalonate pathway^80, 81^ and been able to synthesize bacterial and archaeal-type lipids^21, 82^. The occurrence of the F- and/or A/V-types ATP synthases in modern Bacteria, suggest their membrane lipids are compatible with either ATP synthase type. It is unclear whether the lack of Archaea with a complete substitution of an A/V-with F-type ATP synthase can be explained by constraints imposed by archaeal membrane lipid composition. Alternatively, it is difficult to exclude the possibility that while the diversification of the catalytic and non-catalytic subunits into the respective F1 and A1V1 families may have predated LUCA, the hexameric headpieces may have functioned independently from extant membrane components and thus differed from canonical membrane-bound ATP synthases^28^. In this scenario, the evolution of the membrane components could have occurred later, potentially in conjunction with the speciation of Bacteria and Archaea and the ‘lipid divide’^35, 76^.

Despite the wide distribution of ATP synthases across cellular life, our analyses revealed that many DPANN Archaea and CPR Bacteria may have minimal complexes, as is the case in the DPANN *Nanoarchaeum equitans*^83^, or even lack all genes for an ATP synthase complex (Figure 1, Supplementary Figure 2, Supplementary Table 4). This finding suggests that ATP synthases are not as essential as previously assumed^31, 32, 44, 63^ with loss in DPANN and CPR lineages likely being the result of genome streamlining processes consistent with their predicted host-dependent lifestyles^56, 57^. For instance, various members of the DPANN lack ATP synthase homologs (60/103) while others encode homologs clustering with other DPANN or potential hosts (Figure 1, Supplementary Figures 6-10, Supplementary Table 4). We observed putative symbiont-host gene transfers between acidophilic Micrarchaeota and their hosts belonging to the Thermoplasmata^84^ consistent with work supporting extensive HGT among ATP synthase genes of acidophilic archaeal lineages ^58^ (Supplementary Figures 6-10, Supplementary Table 4). Furthermore, our trees show support for HGT of the *c*A1V1 subunit between the symbiotic Nanohaloarchaeota and their halobacterial hosts (Supplementary Figure 6). The possible HGT of ATP synthase genes underpins the debate over the phylogenetic placement of Nanohaloarchaeota^44^, originally placed as sister-group to Halobacteria^44, 85^ but later recovered as a member of the DPANN lineage^16, 17, 86–88^. Recently, Feng and coworkers found that the catalytic and non-catalytic subunits of the A/V-type ATP synthase of Nanohaloarchaeota form sister-groups to halobacterial homologs^44^. Although their concatenated species trees placed Nanohaloarchaeota with other DPANN^44^, the authors argued that this placement was an artifact due to compositional biases in the concatenated dataset, with the ATP synthase gene tree recording the true organismal history. By contrast, Wang et al. (2019) suggested that the incongruence of the species tree and ATP synthase gene trees for halophilic Archaea result from the HGT of an ATP synthase operon from Halobacteria into the the common ancestor of Nanohaloarchaeota; ecological association and symbiotic interactions between these organisms might have facilitated such a transfer. We include a larger representation of DPANN and MAGs (GCA_003660905, GCA_003660865) belonging to a divergent sister lineage of the Nanohaloarchaeota (Figure 1, Supplementary Figure 2, Supplementary Figure 20), providing an opportunity to reconsider and distinguish hypotheses. Our results group Nanohaloarchaeota *nc*A1V1 subunits with Halobacteria, while the *nc*A1V1 genes of the Nanohaloarchaeota sister-lineage branch with other DPANN; in the *c*A1V1 subtree, the Nanohaloarchaeota group with DPANN (including the sister lineage, Supplementary Figures 6-10). Our analyses are most compatible with a scenario in which the nanohaloarchaeotal ancestor already possessed an ATP synthase complex inherited vertically from its DPANN relatives and replaced one subunit (*nc*A1V1) through HGT from a halobacterial host early during the lineage’s evolution, potentially as an adaptation to halophily. Alternatively, the *nc*A1V1 subunit might be compositionally attracted to homologs of the Halobacteria as a result of convergent adaptations to halophily. This suggests that even genes whose synteny is conserved across lineages may be individually affected by HGT and can have distinct evolutionary histories; in such cases, the phylogenetic signal encoded in a larger number of marker genes may provide a more reliable estimate of the species tree.

Thus, ATP synthase evolution appears complex involving both vertical and horizontal components. Consistent with observations that niche expansion of Thaumarchaeota into acidic soils and high pressure oceanic zones was linked to their horizontal acquisition of a variant V-type ATP synthase operon^58^, the instances of HGT presented here illustrate the potential role of symbiont-host gene exchange and environmental factors in ATP synthase evolution. Prospective studies focusing on genome evolution of DPANN archaea, can help further assess the presence of ATP synthases and other metabolic components in the various DPANN ancestors and elucidate instances of transfer and loss of genes throughout DPANN diversification and adaptation to their respective symbiotic hosts.

Our molecular clock analyses suggest cross-bracing nodes is effective in propagating temporal information across the tree of life, using the eukaryotic fossil record to improve the precision and accuracy of divergence time estimates within the prokaryotic domains. Bracing resulted in higher estimated rates of molecular evolution overall (Supplementary Figure 17, Supplementary Table 10), with the result that various deeper nodes of the tree were estimated to be slightly younger when compared with the unbraced analyses. For example, in the absence of bracing, the posterior age distribution for LUCA approaches the root maximum (the moon-forming impact at 4.52Ga), as has been observed in previous analyses using this maximum age^19, 27^. In contrast, LUCA is estimated as being slightly younger (Figure 5A, Figure 5C, Supplementary Table 10) in braced analyses. This suggests that bracing helps to ameliorate the problem of an under-calibrated clock inferring rates that are too low to account for the amount of genetic change that has occurred since the root of the universal tree^19, 89^. See Supplementary Discussion for further details about the resulting age estimates for major prokaryotic clades.

The LECA estimate (1.93-1.84Ga, 95% HPD) from our species tree analysis falls within published molecular clock estimates, placing LECA within a broad interval ∼1-2.4Ga^25–27, 90–92^ (Figure 5A and 5C, Supplementary Figure 16, Supplementary Table 10). More recent analyses have tended to resolve an older LECA, with ages closer to 1Ga being less plausible on the basis of fossils from that period that can uncontroversially be assigned to crown Archaeplastida. These fossils include the green alga *Proterocladus antiquus* (1Ga)^93^ and the red alga *Bangiomorpha pubescens* (>1030 Mya)^94^. The ages of some of these nodes, including LECA and particularly the last plastid common ancestor (LPCA), were inferred to be younger in the ATP synthase analysis (Figure 5, Supplementary Figures 16 and 18, Supplementary Table 10). This may partly be due to the shorter alignment of ATP synthase (433-512AA, Supplementary Figures 4-7) homologs resulting in low resolution^95^ and subsequent inability to apply all of the species tree calibrations and braces to the ATP synthase phylogeny through gene and species tree incongruence (Supplementary Information). Our analyses are of interest for the timing of mitochondrial acquisition relative to other hallmark features of eukaryotes such as the nucleus^96, 97^, and help to explain the differences in the length of the stem between eukaryotic genes of archaeal and bacterial origin reported previously^96, 98^. Note that while the LECA nodes within the mitochondrial and nuclear lineages can be cross-braced to the same unknown age, the lengths of the antecedent stems, that is, the divergence times of the mitochondrial and nuclear lineages from their closest bacterial and archaeal relatives, might be very different. Our analyses support a moderately longer stem for the nuclear lineage (520.3Ma (mean), 291 — 789 Ma, 95% HPD) than the mitochondrion (438.8Ma, 233 — 682Ma, 95% HPD), suggesting the divergence of the nuclear lineage from the closest sampled Asgard occurred before the divergence of the mitochondrial lineage from Alphaproteobacteria. However, the credible age ranges for these divergences overlap, therefore some additional factor - potentially a faster evolutionary rate prior to LECA in eukaryotic genes of archaeal origin - may contribute to the observed differences in stem lengths between these two categories of genes^96, 98^. Interestingly, the inferred timescale is sensitive to the phylogenetic position of eukaryotes within Asgard archaea and Alphaproteobacteria: in an alternative analysis in which eukaryotes were placed sister to all Asgard archaea, and mitochondria within Alphaproteobacteria, the difference in stem group ages was more pronounced (mean 812.1Ma, 95% HPD 540-1105Ma, nuclear stem: 310.1Ma, mitochondrial stem: 150-508Ma) (Supplementary Figure 12). While this result tells us something about the shape of the tree of life it does not distinguish between hypotheses of an “early” or “late” mitochondrial acquisition. This is because these hypotheses make competing predictions about the order in which key features without direct correspondence to nodes in the tree were acquired relative to the mitochondrial endosymbiosis, which does not exist as a specific node in the tree.

## Conclusions

Our analyses including ancestral gene and genome reconstruction, as well as our novel cross-braced molecular clock approach provide insights into the diversification of the ATP synthase gene family in light of a dated tree of life and established age estimates for key nodes in the tree of life (e.g. LUCA, LACA, LBCA, and LECA). Our results suggest that while LACA solely harbored an A1/V1-type ATP synthase, LBCA may already have encoded homologs of the head component of both the F- and a A/V-type ATP synthase. Studying how A/V-type ATP synthases function in Bacteria will help to explain the distribution we observed and the functional consequences of the ancient divergence between F- and A/V-type ATP synthases. In contrast to previous work, our inferences are consistent with the hypothesis that the divergence of the F1- and A/V1-type ATP synthase components may have predated LUCA. Furthermore, ATP synthase evolution supports scenarios on eukaryotic origins from an Asgard archaeal host^3, 4, 13, 14^ and alphaproteobacterial symbiont^99, 100^ and, together with our dated species tree, provide an updated timescale of cellular evolution, placing the origin of the eukaryotic cell into a geological context that can help to test eukaryogenesis models.

## Figure legends

## Online Methods

### Selection of 800 taxa comprising the backbone genome reference dataset

#### Archaeal reference genomes

A representative set of archaeal genomes was selected from a broad phylogeny of all archaeal genomes present in NCBI. A set of 51 marker proteins^87^ was used to infer an initial concatenated phylogeny of 574 archaeal genomes meeting a threshold of >40% completeness and <13% contamination (Supplementary Table 1). Individual markers were aligned with MAFFT L-INS-i (settings: –reorder)^108^, trimmed using BMGE (settings: -m BLOSUM30 -h 0.55)^101^ and concatenated with a custom script (catfasta2phyml.pl; https://github.com/nylander/catfasta2phyml). A phylogenetic tree was generated with IQ-TREE v1.6.7 (settings: -m LG+C60+F+R -bb 1000 -alrt 1000)^102^.

Based on this tree, 350 archaeal genomes were subsampled to evenly represent archaeal phylogenetic diversity (Supplementary Table 1). Type-strains were preferentially selected, while high quality MAGs and SAGs were selected based on completeness and contamination levels.

#### Bacterial reference genomes

The bacterial reference backbone, prioritizing type-strains and reference genomes, but also high-quality MAGs and a subselection of representatives from candidate phyla, was derived using an initial phylogeny of bacterial genomes available in NCBI as described above. Homologs of a conserved set of 29 marker proteins, i.e. a subset of 48 single-copy marker proteins previously defined in Zaremba-Niedzwiedzka et al. (2017)^4^ were identified in those bacterial genomes, aligned using MAFFT (settings: –reorder)^108^, trimmed using (settings: -m BLOSUM30 -h 0.55)^101^ and concatenated to reconstruct a phylogenetic tree using IQ-TREE v1.6.7 (settings: -m LG+G -bb 1000 -alrt 1000)^101, 102^. We subsampled the concatenated phylogeny for 349 bacterial genomes that represent known bacterial genomic diversity, ensuring selection of major bacterial taxonomic clades. The genome of *Schaalia odontolytica* ATCC 17982, which represents the host of members of the Saccharibacteria (formerly phylum TM7)^109, 110^, was downloaded from NCBI in 2020 and manually added to the bacterial backbone dataset (Supplementary Table 1).

#### Eukaryote reference genomes

A set of 100 published genome-wide datasets (genomes and, for lineages lacking complete genomes, largely complete transcriptomes) were sampled to represent the major lineages of eukaryotes (Supplementary Table 1, Supplementary Table 11). We also included sequences from the unpublished *Diplonema papillatum* genome project, with the permission of the sequencing consortium (see Acknowledgements).

#### Functional annotations

To identify sequences of ATP synthase subunits within all genomes in the 800-backbone set, all protein coding sequences were annotated using the KEGG and COG databases. Sequences were compared to KO profiles within the KEGG Automatic Annotation Server (KAAS, downloaded April 2019) (KAAS; downloaded April 2019)^111^, to COG profiles within the NCBI COG database (downloaded May 2020)^112–114^, and to Pfam profiles in the Pfam database (Release 34.0)^115^. KOs and COGs were assigned using hmmsearch v3.1b2 (settings: --tblout sequence_results.txt -o results_all.txt -- domtblout domain_results.txt --notextw -E 1e-5)^116^. Pfams were assigned using hmmsearch v3.1b2 (settings: --tblout sequence_results.txt -o results_all.txt --domtblout domain_results.txt --notextw -E 1e-10)^116^.

### Inference of a concatenated species phylogeny including Archaea, Bacteria and Eukaryotes

#### Marker gene homology search

A concatenated phylogeny of the 800 bacterial, archaeal, and eukaryotic genomes included in this study was inferred using a previously defined set of 27 single-copy marker genes^19^ (Supplementary Figure 20, Supplementary Table 12). To collect the corresponding homologs, the 800 reference genomes were queried against all COG HMM profiles with a custom script built on the hmmsearch [options] <reference genomes> <hmmfile> algorithm^117^: hmmsearchTable Whole_ArcBacEuk_800_vs2_clean.faa NCBI_COGs_Oct2020.hmm -E 1e-5 > 1_Hmmersearch/HMMscan_Output_e5 (HMMER v3.3.2)^118^, and all homologs corresponding to the 27 single-copy marker genes were identified, cleaned, and parsed. The approaches used to identify the appropriate homologs for prokaryotes and eukaryotes are described below:

#### Selection of prokaryotic homologs

For prokaryotes, the best-hit sequences were selected based on e-value and bitscore and the corresponding protein sequences were extracted from the reference genome backbone. Protein sequences assigned to each marker gene were aligned using MAFFT L-INS-i v7.453 (settings: --reorder)^108^ and trimmed using BMGE (settings: -t AA -m BLOSUM30 -h 0.55)^101^. Maximum-likelihood phylogenies with ultrafast bootstrap approximation (UFBoot) for each single-copy marker gene were constructed using IQ-TREE2 v2.1.2 (settings: -m LG+G -wbtl -bb 1000 -bnni)^102–104^. Individual marker gene trees were manually inspected for domain-level monophyly, the presence of paralogous protein families, and signs of contamination including LBA and horizontal gene transfer (HGT) (Supplementary Table 12, Data repository: 10.5281/zenodo.7807739). Marker genes, where domain-level lineages were paraphyletic were excluded and sequences with indications of LBA, HGT, and paralogy were manually removed using a custom script: remove_seq_with_specific_header3.py.

#### Selection of nuclear eukaryotic homologs

To distinguish between the nuclear, plastid, and mitochondrial homolog and select the correct eukaryotic representative sequence, we relaxed the selection criteria and in place of the best hit using e-value and bitscore, we collected all eukaryotic hmmsearch hits and downsampled them with CD-HIT using a threshold of 90% sequence identity^119, 120^. The filtered eukaryotic sequences were combined with the previously inspected prokaryotic sequences and all sequences for each single-copy marker gene were aligned using MAFFT L-INS-i v7.453 (settings: --reorder)^108^, trimmed using BMGE v1.12 (settings: -t AA -m BLOSUM30 -h 0.55)^101^, and single gene phylogenies were inferred using FastTree (settings: -lg)^121^. KEGG and Pfam annotations (see above) were mapped to the tips of the eukaryotic sequences for manual inspection of multiple paralogs per taxon. First, the eukaryotic sequences were inspected by removing any sequence failing monophyly (i.e., HGTs in prokaryotic clades) or not clearly derived from the nuclear source (i.e., the plastid and/or mitochondrial sequences). Duplicate nuclear eukaryotic sequences were filtered in a two-step procedure: 1) if duplicate sequences are monophyletic, select a single representative based on protein annotation consistent with the identity of the single-copy marker gene, and 2) if duplicate sequences are paraphyletic, remove taxon completely from the single-copy marker gene. The presence/absence distribution of the 21 single-copy marker genes was calculated for all eukaryotic taxa and any representatives with fewer than 65% of the marker genes (20 taxa removed, 80 eukaryotes in total) were removed from this analysis (Supplementary Table 13).

#### Inspection of final marker gene sequence sets

The final set of eukaryotic nuclear sequences were combined with the previously inspected sequences for archaea and bacteria (see above) and aligned using MAFFT L-INS-i v7.453^108^, trimmed with BMGE v1.12 (settings: -t AA -m BLOSUM30 -h 0.55)^101^, and single gene trees were inferred using maximum likelihood with UFBoot approximation methods in IQ-TREE2 v2.1.2 (settings: -m LG+G -wbtl -bb 1000 -bnni)^102–104^. Upon inspection of single gene trees including homologs from Archaea, Bacteria and Eukaryotes, six single-copy markers (COG0064, COG0085, COG0086, COG0202, COG0480, and COG5257 (Supplementary Table 12) were flagged for removal (e.g. lack of clear nuclear paralog, or absence of archaeal or bacterial sequences in the tree).

#### Inference of the concatenated phylogeny

Alignments for the 21 single-copy marker genes were generated and trimmed following the approaches outlined above and individual marker alignments were concatenated using the script catfasta2phyml.pl (https://github.com/nylander/catfasta2phyml). The final concatenated sequence alignment contained 3367 positions and was used to infer maximum-likelihood phylogenies using varying models of evolution in IQ-TREE2 v2.1.2 (settings: -m LG+C60+R+F *or* LG+C20+R+F -bb 1000 -alrt 1000)^102–104^. We used the Approximately-Unbiased (AU) test to statistically assess confidence in differing tree topologies^122^. Specifically, we examined the statistical support for topologies of the two concatenated species trees inferred under different models of evolution (LG+C60+R+F and LG+C20+R+F, see above). The AU test was implemented in IQ-TREE2 v.2.1.2 (settings: - s 21eLife_ArcBacEuk_wHuber_vs1.faa -m [LG+C20+R+F/LG+C60+R+F] -z 21eLife_ArcBacEuk_wHuber_vs1_bothtrees.treefile -pre [C20/C60] -n 0 -zb 10000 -au^102, 104, 122^. Results are shown in Supplementary Table 15. Despite statistical exclusion of the LG+C20+R+F topology, we chose to use this tree for phylogenetic interpretation because the placement of key lineages such as the Asgard Archaea and CPR, is most consistent with recent evidence^4, 19, 21, 22^ (Supplementary Figure 20, Supplementary Table 15).

### Constructing a ribosomal marker phylogeny including nuclear, mitochondria, and plastid homologs

#### Selection of eukaryotic nuclear, mitochondrial, and plastid homologs

Eukaryotes encode two or more ribosomes of distinct prokaryotic origins, i.e. archaeal, alphaproteobacterial and, in the case of the presence of a plastid, a cyanobacterial origin (hereafter: the nuclear, mitochondrial, and plastid, respectively). A concatenated phylogeny including, if identified, the nuclear, mitochondrial, and plastid ribosomal protein homologs for each eukaryote, was inferred for molecular dating and bracing analyses. To this end, we constructed single gene trees of the 15 ribosomal marker genes (subset of the 21 single-copy marker genes described above) which included all eukaryotic homologs (i.e., the nuclear, mitochondrial, and plastid). Note that the nuclear eukaryotic sequences were the same set of sequences reported in the final inspection of the concatenated species phylogeny (see above). To identify the plastid homologs, we selected the monophyletic clade of eukaryotic sequences affiliated with the Cyanobacteria. The mitochondrial sequences appeared to demonstrate variable placements with some affiliating with the alphaproteobacteria and others branching basally in the Bacteria. Therefore, we made our sequence selection based on the position of known mitochondrial genes of the type-species *Homo sapiens* and *Saccharomyces cerevisiae*. First, we manually located *H. sapiens* in the phylogenies and searched the protein accession in Uniprot and/or NCBI^123^ to confirm sequence annotation and identity as a mitochondrial sequence. In the absence of a *H. sapiens* homolog, we used *S. cerevisiae* mitochondrial homologs. Of the 15 ribosomal markers, three had no distinguishable mitochondrial homolog for either type-species and were dropped from the dataset, therefore resulting in 12 ribosomal markers (Supplementary Table 16). All eukaryotic sequences, that clustered with the *H. sapiens/S. cerevisiae* homolog and formed either a clade with alphaproteobacteria or a basal clade in the phylogeny, were selected for subsequent analyses. Selected sequences were de-replicated using the following criteria: 1) if paralogous sequences are monophyletic retain one homolog based on annotation or manual selection, and 2) if paralogous sequences are paraphyletic remove all sequences from that organism. Dereplicated sequences marked for removal are in Supplementary Table 16. Gene trees with selected sequences have been deposited in data repository: 10.5281/zenodo.7807739.

Ribosomal protein homologs were then annotated based on their distinct origin (nuclear, mitochondrial, plastid) and the percent distribution of homologs across the 12 ribosomal markers by eukaryotic taxon was calculated. Only taxa that had at least 50% of the markers of nuclear, mitochondrial, or plastid origin were retained, resulting in 88 nuclear taxa, 50 mitochondrial taxa, and 25 plastid taxa (Supplementary Table 13). The eukaryotic GenomeIDs for each sequence were annotated with the suffix of the origin (i.e., EukGenome_nuclear, EukGenome_mito, etc.) for downstream concatenation. In total, the sequence sets for the 12 ribosomal markers contained archaeal and bacterial homologs, and the eukaryotic nuclear, mitochondrial, and plastid sequences, respectively. Alignments were generated using MAFFT L-INS-i v7.453 (settings: --reorder)^108^ and trimmed with TRIMAL v1.2rev59 (settings: -gappyout)^107^. The alignments of the 12 ribosomal markers were concatenated using the script catfasta2phyml.pl (https://github.com/nylander/catfasta2phyml) and the final concatenated alignment contained 2133 sites.

#### Inference of concatenated phylogenies

A maximum-likelihood phylogeny was inferred using IQ-TREE2 v.2.1.2 (settings: -m LG+C60+R+F -bb 1000 -alrt 1000)^102–104^ (Supplementary Figure 21).

### Assessing distribution of ATP synthase genes across 800 taxa backbone

We performed a comparative genomic analysis of the distribution of ATP synthase genes across the 800 taxa included in this study. COG families corresponding to each subunit of the ATP synthase (Supplementary Table 3) were extracted from the 800 reference genomes. Results were compiled, counted in R v4.1.1, and used to generate a count table of each gene within each reference genome (Supplementary Table 4). The count table was converted to a binary presence/absence matrix that was summarized using the ddply function of the plyr package (v1.8.6) by the respective phylogenetic clustering methods: 1) species-level according to tip order of individual species in the inferred concatenated phylogeny (BinID and Tip_Order, Supplementary Table 17), and 2) class- and phylum-level for Archaea and phylum-level for Bacteria corresponding to clade clustering in the concatenated phylogeny (CladeCluster and Clade_Order, Supplementary Table 17). The percentage distribution of subunits within each phylogenetic cluster was visualized in a bubble plot implemented using the ggplot function with geom_tile and facet_grid from the ggplot2 package v3.3.5. The binary presence/absence of subunits by species was visualized with the ggplot function using geom_point and facet_grid from ggplot2 v3.3.5. All heatmaps and bubble plots were manually merged with the corresponding concatenated species tree in Adobe Illustrator CC v22.0.1.

### Phylogenetics of ATP synthase subunits

#### Sequence retrieval and selection

Interpro domains that characterize the protein families corresponding to the subunits present in the catalytic (R1) domain of the F-Type and A/V-Type ATP synthases were selected at the family-level^124^ and include: ipr005294 (F-Type alpha, hereafter *nc*F1), ipr005722 (F-Type beta, hereafter *c*F1), ipr022878 (A/V-Type A, hereafter *c*A1/V1), and ipr022879 (A/V-Type B, hereafter *nc*A1/V1). All protein sequences assigned to the corresponding interpro domains were extracted from the UniProt Knowledge Base^123^, and were searched against the 800 reference genomes using DIAMOND v0.9.22.123 (settings: blastp -p 4 -f 6 qseqid stitle pident length mismatch gapopen qstart qend sstart send e-value bitscore)^125^. Top hits were selected based on best e-value and sequence identity, and all unique protein accessions (from the 800 reference taxa) were used to extract the amino acid sequences from the 800-genome reference dataset. Sequences with undefined characters (i.e., X, x) and/or outside of the average sequence length of homologs, i.e. 300-675 bp, were filtered from the sequence sets. To avoid highly similar duplicates, sequences with 99-100% identity were removed using CD-HIT^119, 120^. Additionally, for consistency with the concatenated species phylogeny (see above), eukaryotic taxa that fell below the 65% marker gene presence cutoff (20 eukaryotic taxa, Supplementary Table 13) were removed from the single-subunit sequence sets.

### ATP synthase subunit phylogenies: *c*F1, *nc*F1, *c*A1V1, *nc*A1V1

#### A series of seven different sequence sets were generated for analysis

1. Single subunits sets F1-alpha (*nc*F1), F1-beta (*c*F1), A1/V1A (*c*A1V1), and A1/V1B (*nc*A1V1) (four in total)
2. Combined orthologous subunits for outgroup rooting: F1A+A1/V1B (*nc*F1+*nc*A1V) and F1B+A1/V1A (*c*F1+*c*A1V1)
3. All four subunits combined

Potential duplicates were removed from the combined sets using CD-HIT with a 100% identity (settings: cd-hit -1)^119, 120^, sequences were aligned using MAFFT L-INS-i v7.453 (settings: --reorder)^108^ and trimmed using BMGE v1.12 (settings: -m BLOSUM30 -h 0.55)^101^. The best-fit model was determined using the Model Finder Plus tool implemented in IQ-TREE2 v.2.1.2 (settings: -m MFP -mset LG -madd LG+C10,LG+C20,LG+C30,LG+C40,LG+C50,LG+C60,LG+C10+R+F,LG+C20+R+F,LG+C30+R+F,LG+C40+R+F,LG +C50+R+F,LG+C60+R+F --score-diff all -bb 1000 -alrt 1000 -bnni -wbtl)^102–104, 106^ and the best-fitting model for each gene tree was selected based on the Bayesian Information Criterion (BIC, Supplementary Table 8) and used to infer the maximum-likelihood phylogeny. Genome identifiers containing the GenomeID and protein accession were converted to a modified NCBI taxonomic string using an in-house script (Replace_tree_names.pl, https://github.com/ndombrowski/Phylogeny_tutorial/tree/main/Input_files/5_required_Scripts). Trees were viewed in FigTree v1.4.4, and inspected for topological congruence and phylogenetic artifacts to iteratively improve the sequence selection, i.e. to exclude distant paralogs and sequences subject to long branch attraction (LBA)^126^.

### Tracing the phylogenetic relationships of Eukaryotic F-Type ATP synthases

To better resolve the evolutionary origins of eukaryotic F-type ATP synthases we constructed phylogenies with a subset of sequences which included eukaryotic sister lineages (alphaproteobacteria and cyanobacteria for the mitochondrial and plastid-type F-ATP synthase, respectively) as well as an outgroup lineage (hereafter: plastid and mitochondria subsets). For the plastid origin dataset, we selected all eukaryotic ATP synthase sequences from the *nc*F1 and *c*F1 subunit gene trees (see above) that clustered with the Cyanobacteria, and added cyanobacterial and melainabacterial homologs. Similarly, the mitochondria origin subset was generated by collecting all eukaryotic ATP synthases sequences from the *nc*F1 and *c*F1 subunit gene trees that clustered with alphaproteobacterial homologs and adding additional alphaproteobacterial sequences and gammaproteobacterial homologs. Note that for both the plastid and mitochondrial sets, we used an expanded selection of cyanobacteria and alphaproteobacteria, respectively, from a larger in-house reference genome set. Sequence selections were filtered to retain high quality sequences without ambiguous amino acids (i.e., X and x, etc.) and within the range of 450-550 bp. Closely related paralogous sequences were removed using CD-HIT (settings: -c 0.99)^119, 120^ and alignments were generated using MAFFT L-INS-i v7.453^108^ and trimmed using BMGE (settings: -m BLOSUM30 -h 0.55)^101^. We inferred phylogenies using the best-fit model determined in the Model Finder Plus tool in IQ-TREE2 v2.1.2 (settings: -m TESTONLY -mset LG -madd LG+C10,LG+C20,LG+C30,LG+C40,LG+C50,LG+C60,LG+C10+R+F,LG+C20+R+F,LG+C30+R+F,LG+C40+R+F,LG +C50+R+F,LG+C60+R+F --score-diff all)^102–104, 106^ and the maximum likelihood trees were constructed in IQ-TREE v1.6.10 using the best-fit model based on the BIC^102^ (Supplementary Table 8).

Additionally, we used Bayesian analysis to further verify the placement of eukaryotic F1-type ATP synthase sequences amongst the proposed sister lineages. Due to computational limitation, we downsampled the taxa subsets containing eukaryotes, alphaproteobacteria, and gammaproteobacteria to a maximum of 250 taxa (*nc*F1: 211, *c*F1:185 sequences). Sequences were cleaned, filtered, de-replicated, aligned, and trimmed using the same conditions described above. Bayesian phylogenies were constructed using PhyloBayes-MPI (version 1.5) using the CAT-GTR model with four discrete gamma categories for rates across sites; for each alignment, four independent Markov Chain Monte Carlo (MCMC) chains were run. Each chain was run over 100,000 iterations (or until convergence) Convergence was evaluated using the bpcomp and tracecomp tools within PhyloBayes-MPI, with 1000 generations discarded as burn-in and sub-sampling every 10 trees. The final consensus trees were generated through bpcomp using the same settings.

### Ancestral sequence reconstruction

Sequence alignments and the accompanying maximum likelihood trees for the ATP synthase subunits, the orthologous pairs, and the set of four combined subunits were used to reconstruct the ancestral protein sequences. For ancestral sequence reconstruction we used a tool implemented in IQ-TREE2 v2.1.2 (settings: -m [model] -asr -te [maximum likelihood tree] -keep_empty_seq)^104^. Ancestral sequences were determined based on the proposed amino acid states at specified node positions in the rooted combined ATP synthase protein tree (Supplementary Figure 10, Supplementary Table 7).

### Conserved nucleotide-binding motifs

Untrimmed and trimmed alignments of F- and A/V-ATP synthase subunits from the 800 reference genomes (see above) were manually inspected in Jalview v2.10.5^127^ for the presence of the WalkerA (P-loop) motif^42^. The signature nucleotide-binding motif is characterized by the amino acid sequence: *GXXXXGK(T/S)* where X denotes any amino acid. The WalkerA motif sequence segment was extracted from the full alignment and used to generate conserved motif logos in WebLogo3 v3.7.4^128^ (http://weblogo.threeplusone.com/).

### Dating the tree of life and ATP synthase phylogenies

The absolute time calibrations used in the dating analysis are detailed in the Supplementary Discussion. As the fossil evidence with which to constrain early microbial evolution is limited, we also used cross-bracing^24^ to propagate the available calibrations across the tree, implemented in McmcDate (see below). In particular, we braced the LECA node that appears in the nuclear and mitochondrial clades (setting their ages to be the same), along with all calibrated nodes within eukaryotes that were present in two or more of the eukaryotic clades (that is, we braced all nodes within eukaryotes where a geological calibration was applied). Finally, we implemented a relative constraint^50^ that the crown plastids must be younger than archaeal- and mitochondrial LECA (Supplementary Discussion).

We used McmcDate (https://github.com/dschrempf/mcmc-date) for molecular dating. McmcDate approximates the phylogenetic likelihood using a multivariate normal distribution obtained from an estimate of the posterior distribution of trees with branch lengths measured in average number of substitutions per site. We estimated the posterior distribution of trees in a previous step. For this previous step, we used PhyloBayes, the LG+G4 model, and a fixed phylogeny as described above. We sampled 10000 values of the posterior distribution of trees and observed good convergence with estimated sample size (ESS) values of around 8000.

Using McmcDate, we sampled 12000 time trees. We used a birth-death tree prior on the time tree, and an uncorrelated log normal relaxed molecular clock model. We calibrated node ages using uniform distributions with bisected normal distributions at the boundaries. Similarly, we constrained the node order using the tails of normal distributions. We set the steepness of the boundaries individually depending on the quality and certainty of the auxiliary data. In a similar way, we used normal distributions to brace nodes. ESS values indicated good convergence and ranged from 3000 to 6000.

Application of fossil calibrations to the inferred ML ribosomal species tree (see above, Supplementary Figure 20) was limited due to poor resolution of within-Eukaryote relationships. To apply an extended set of fossil calibrations we fixed the within-eukaryote topology to reflect established relationships among supergroups^105^ and to allow the within-eukaryote fossil calibrations to be applied to the tree (see calibrations justification, Supplementary Discussion). In addition to these eukaryotic constraints, one topology (hereafter, Edited1) placed the nuclear eukaryotic homologs as the sister lineage to all Asgard archaea and the mitochondrial homologs as the sister lineage to a single *Neorickettsia* (Supplementary Figure 12). We used a more conservative approach to investigate the timing of LECA via the nuclear and mitochondrial eukaryotic nodes by adding additional constraints (hereafter, Edited2) to position the nuclear homologs as the sister lineage to the Hodearchaea (formerly Heimdallarchaeota LC3), their predicted closest asgardarchaeal relatives^4, 129^, and the mitochondrial homologs sister to all alphaproteobacteria, consistent with previous work^99, 100^ (Supplementary Figure 13). The focal analysis described here is derived from dating the Edited2 topology (Figure 5C, Supplementary Figures 13 and 16). An AU test was applied to assess the statistical support for the different topologies inferred from the ribosomal species trees utilized for cross-calibrated dating. The AU test was implemented in IQ-TREE2 v.2.1.2^102–104^ (settings: iqtree2 -s 12Ribosomal_eLife_ArcBacEuk_gappyout_v1b.faa -m LG+C60+R+F -z Ribo_C60_trees.alltrees.treefile -n 0 -zb 10000 -au). Results are shown in Supplementary Table 18.

In formulating the calibrations (Supplementary Discussion), we sought to follow the best practice principles set out in Parham et al. (2012). However, these were designed with animal and plant fossil-based calibrations and not all of the principles are applicable to calibrations of microbial clades which often lack phenotypic synapomorphies, let alone diagnostic characters that are preserved in fossil remains. Furthermore, the calibrations for many clades rely on geochemical evidence of microbial metabolisms, manifest as isotope fractionation or oxidation states of redox sensitive mineral species.

Consequently, we have adapted the best practice principles to suit the nature of the calibrations. Novel calibrations are justified in full; we indicate the source of calibrations that are justified elsewhere, providing notes where they have been adapted for different clades or where the dating has changed subtly with the revision of the geologic timescale (Supplementary Discussion).

### Gene tree-species tree reconciliation

Ultrafast bootstraps (UFBoot) were inferred for each of the ATP synthase gene trees (see above) in IQ-TREE2 v2.1.2^102–104^, and the inferred ML concatenated species trees (see above). ALEobserve was used to convert bootstrap distributions into ALE objects, which were reconciled using ALEml_undated against each of the four species trees: those with eukaryotes, using the LG+C20+R+F and LG+C60+R+F model, and those lacking eukaryotic sequences, with the same two models (Supplementary Table 6). These four species tree topologies were also rooted in two different ways: a root between Archaea and Bacteria, and a root between Gracilicutes and all other taxa. Two approaches were taken using ALE for gene tree-species tree reconciliation. First, we used the default ALE parameters, i.e. inferring the probability that each subunit originated at the LUCA, LBCA and LACA nodes on the prior assumption that origination at any internal node of the species tree was equally likely. We also tested an alternative approach^21^ in which the origination probability at the root (O_R) is different to the origination probability for all other internal nodes of the tree, with O_R estimated by maximum likelihood. Reconciliation analyses were performed using ALE v1.0 (https://github.com/ssolo/ALE).

To compare support for the traditional Archaea-Bacteria root for the tree of life, and an alternative root within the Bacteria, we used gene tree-species reconciliation. We performed gene tree-species tree reconciliation using the species tree as described above as well as individual gene family subunit trees of ATP synthase: *nc*F1 (F1 alpha), *c*F1 (F1 beta), *c*A1V1 (A1/V1 A), and *nc*A1V1 (A1/V1 B), as well as three combined gene families, *nc*F1+*nc*A1V1, *c*F1+*c*A1V1, and all four families combined (Supplementary Table 6). Two taxon samplings were used as described above, one with 350 Archaea and 350 Bacteria only, and another with 350 Archaea, 350 Bacteria, and 100 eukaryotes. The summed gene family likelihoods of each ATP synthase subunit were compared using an AU test^122^ as implemented in CONSEL^130^ under a range of conditions: species trees inferred under the LG+C20+R+F and LG+C60+R+F models; samples including and excluding eukaryotes, and two different root positions, one with the traditional root between Archaea and Bacteria, and the second with a, within-Bacteria, root on the branch leading to Gracilicutes.

## Acknowledgements

This work was supported by the Gordon and Betty Moore Foundation through grant GBMF9741 to TAW, AS, and GJSz. Furthermore, A.S. has received funding from the Swedish Research Council (VR starting grant 2016-03559), the NWO-I foundation of the Netherlands Organisation for Scientific Research (WISE fellowship) and the European Research Council (ERC) under the European Union’s Horizon 2020 research and innovation programme (grant agreement No. 947317, ASymbEL). Further, this work was supported by a Royal Society University Research Fellowship to TAW. ERRM, DP, PCJD and TAW were supported by the John Templeton Foundation (62220). The opinions expressed in this publication are those of the author(s) and do not necessarily reflect the views of the John Templeton Foundation. GJSz, DS and LLSz received funding from the European Union’s Horizon 2020 research and innovation programme (grant agreement No. 714774, GENECLOCKS).

We thank Gertraud Burger, Julius Lukes, Takeshi Nara, and other members of the Diplonema papillatum sequencing consortium for sharing data. We also want to thank Courtney Stairs and Georg Hochberg for helpful discussions regarding eukaryotic metabolism and ancestral sequence reconstructions, respectively.

## Author contributions

AS and TAM conceptualized the study. TAM, ERRM, TAW, DS, LLSz, ND, GJSz, and AS performed analyses and interpreted data. DP and PCJD contributed data. AS, TAW, GJSz and PCJD acquired funding. TAM wrote the first draft with the help of AS and TAW. TAM, AS, TAW, ERRM, GJSz, and PCJD contributed to the writing of the manuscript and all authors read and approved the final version.

## Data availability

All genomic data of Archaea and Bacteria analyzed are available at NCBI (Supplementary Table 1), while all eukaryotic genomic/transcriptomic material is deposited in our data repository: 10.5281/zenodo.7807739). Additional supplementary files including single gene tree analysis and concatenated phylogenies (i.e., sequence files, alignments, and treefiles) are deposited also in our data repository, 10.5281/zenodo.7807739. Databases used in this study are detailed as follows: KO profiles downloaded from the KEGG Automatic Annotation Server in 2019 [https://www.genome.jp/tools/kofamkoala/], and the NCBI COG Database downloaded May 2020 [https://ftp.ncbi.nih.gov/pub/COG/COG2020/data/].

## Code availability

Workflows for annotations and phylogenies and custom R scripts to analyze and parse annotation data for figure generation have been deposited in our data repository: 10.5281/zenodo.7807739. In addition, we used the following published code: catfasta2phyml.pl [https://github.com/nylander/catfasta2phyml].

## Notes

https://doi.org/10.5281/zenodo.7807739

### Competing Interest Statement

The authors have declared no competing interest.

## References

1. Spang, A., Mahendrarajah, T. A., Offre, P. & Stairs, C. W. Evolving Perspective on the Origin and Diversification of Cellular Life and the Virosphere. Genome Biol. Evol. 14, (2022).

2. Hug, L. A. et al. A new view of the tree of life. Nat Microbiol 1, 16048 (2016).

3. Spang, A. et al. Complex archaea that bridge the gap between prokaryotes and eukaryotes. Nature 521, 173–179 (2015).

4. Zaremba-Niedzwiedzka, K. et al. Asgard archaea illuminate the origin of eukaryotic cellular complexity. Nature 541, 353–358 (2017).

5. Spang, A. et al. Asgard archaea are the closest prokaryotic relatives of eukaryotes. PLoS genetics vol. 14 e1007080 (2018).

6. Rinke, C. et al. A standardized archaeal taxonomy for the Genome Taxonomy Database. Nat Microbiol 6, 946–959 (2021).

7. Williams, T. A., Cox, C. J., Foster, P. G., Szöllősi, G. J. & Embley, T. M. Author Correction: Phylogenomics provides robust support for a two-domains tree of life. Nat Ecol Evol 4, 1568 (2020).

8. Liu, Y. et al. Expanded diversity of Asgard archaea and their relationships with eukaryotes. Nature 593, 553–557 (2021).

9. Guy, L., Saw, J. H. & Ettema, T. J. G. The archaeal legacy of eukaryotes: a phylogenomic perspective. Cold Spring Harb. Perspect. Biol. 6, a016022 (2014).

10. Martin, W. F., Garg, S. & Zimorski, V. Endosymbiotic theories for eukaryote origin. Philosophical Transactions of the Royal Society B: Biological Sciences vol. 370 20140330 Preprint at https://doi.org/10.1098/rstb.2014.0330 (2015).

11. Koonin, E. V. Origin of eukaryotes from within archaea, archaeal eukaryome and bursts of gene gain: eukaryogenesis just made easier? Philos. Trans. R. Soc. Lond. B Biol. Sci. 370, 20140333 (2015).

12. López-García, P., Eme, L. & Moreira, D. Symbiosis in eukaryotic evolution. Journal of Theoretical Biology vol. 434 20–33 Preprint at https://doi.org/10.1016/j.jtbi.2017.02.031 (2017).

13. Spang, A. et al. Proposal of the reverse flow model for the origin of the eukaryotic cell based on comparative analyses of Asgard archaeal metabolism. Nat Microbiol 4, 1138–1148 (2019).

14. Imachi, H. et al. Isolation of an archaeon at the prokaryote-eukaryote interface. Nature 577, 519– 525 (2020).

15. López-García, P. & Moreira, D. The Syntrophy hypothesis for the origin of eukaryotes revisited. Nat Microbiol 5, 655–667 (2020).

16. Rinke, C. et al. Insights into the phylogeny and coding potential of microbial dark matter. Nature 499, 431–437 (2013).

17. Castelle, C. J. et al. Genomic expansion of domain archaea highlights roles for organisms from new phyla in anaerobic carbon cycling. Curr. Biol. 25, 690–701 (2015).

18. Brown, C. T. et al. Unusual biology across a group comprising more than 15% of domain Bacteria. Nature 523, 208–211 (2015).

19. Moody, E. R. R. et al. An estimate of the deepest branches of the tree of life from ancient vertically evolving genes. Elife 11, (2022).

20. Taib, N. et al. Genome-wide analysis of the Firmicutes illuminates the diderm/monoderm transition. Nat Ecol Evol 4, 1661–1672 (2020).

21. Coleman, G. A. et al. A rooted phylogeny resolves early bacterial evolution. Science 372, (2021).

22. Martinez-Gutierrez, C. A. & Aylward, F. O. Phylogenetic Signal, Congruence, and Uncertainty across Bacteria and Archaea. Mol. Biol. Evol. 38, 5514–5527 (2021).

23. Kapli, P., Flouri, T. & Telford, M. J. Systematic errors in phylogenetic trees. Current Biology vol. 31 R59–R64 Preprint at https://doi.org/10.1016/j.cub.2020.11.043 (2021).

24. Shih, P. M. & Matzke, N. J. Primary endosymbiosis events date to the later Proterozoic with cross-calibrated phylogenetic dating of duplicated ATPase proteins. Proceedings of the National Academy of Sciences vol. 110 12355–12360 Preprint at https://doi.org/10.1073/pnas.1305813110 (2013).

25. Parfrey, L. W., Lahr, D. J. G., Knoll, A. H. & Katz, L. A. Estimating the timing of early eukaryotic diversification with multigene molecular clocks. Proc. Natl. Acad. Sci. U. S. A. 108, 13624–13629 (2011).

26. Eme, L., Sharpe, S. C., Brown, M. W. & Roger, A. J. On the age of eukaryotes: evaluating evidence from fossils and molecular clocks. Cold Spring Harb. Perspect. Biol. 007, (2014).

27. Betts, H. C. et al. Integrated genomic and fossil evidence illuminates life’s early evolution and eukaryote origin. Nat Ecol Evol 2, 1556–1562 (2018).

28. Mulkidjanian, A. Y., Makarova, K. S., Galperin, M. Y. & Koonin, E. V. Inventing the dynamo machine: the evolution of the F-type and V-type ATPases. Nat. Rev. Microbiol. 5, 892–899 (2007).

29. Stewart, A. G., Laming, E. M., Sobti, M. & Stock, D. Rotary ATPases—dynamic molecular machines. Current Opinion in Structural Biology vol. 25 40–48 Preprint at https://doi.org/10.1016/j.sbi.2013.11.013 (2014).

30. Iwabe, N., Kuma, K., Hasegawa, M., Osawa, S. & Miyata, T. Evolutionary relationship of archaebacteria, eubacteria, and eukaryotes inferred from phylogenetic trees of duplicated genes. Proc. Natl. Acad. Sci. U. S. A. 86, 9355–9359 (1989).

31. Gogarten, J. P. et al. Evolution of the vacuolar H+-ATPase: implications for the origin of eukaryotes. Proc. Natl. Acad. Sci. U. S. A. 86, 6661–6665 (1989).

32. Hilario, E. & Gogarten, J. P. Horizontal transfer of ATPase genes — the tree of life becomes a net of life. Biosystems vol. 31 111–119 Preprint at https://doi.org/10.1016/0303-2647(93)90038-e (1993).

33. Cross, R. L. & Müller, V. The evolution of A-, F-, and V-type ATP synthases and ATPases: reversals in function and changes in the H /ATP coupling ratio. FEBS Letters vol. 576 1–4 Preprint at https://doi.org/10.1016/j.febslet.2004.08.065 (2004).

34. Mulkidjanian, A. Y., Galperin, M. Y., Makarova, K. S., Wolf, Y. I. & Koonin, E. V. Evolutionary primacy of sodium bioenergetics. Biol. Direct 3, 13 (2008).

35. Mulkidjanian, A. Y., Galperin, M. Y. & Koonin, E. V. Co-evolution of primordial membranes and membrane proteins. Trends Biochem. Sci. 34, 206–215 (2009).

36. Matzke, N. J., Lin, A., Stone, M. & Baker, M. A. B. Flagellar export apparatus and ATP synthetase: Homology evidenced by synteny predating the Last Universal Common Ancestor. Bioessays 43, e2100004 (2021).

37. Müller, V. & Grüber, G. ATP synthases: structure, function and evolution of unique energy converters. Cell. Mol. Life Sci. 60, 474–494 (2003).

38. Kühlbrandt, W. Structure and Mechanisms of F-Type ATP Synthases. Annu. Rev. Biochem. 88, 515– 549 (2019).

39. Gogarten, J. P. & Taiz, L. Evolution of proton pumping ATPases: Rooting the tree of life. Photosynth. Res. 33, 137–146 (1992).

40. Grüber, G., Manimekalai, M. S. S., Mayer, F. & Müller, V. ATP synthases from archaea: the beauty of a molecular motor. Biochim. Biophys. Acta 1837, 940–952 (2014).

41. Forgac, M. Vacuolar ATPases: rotary proton pumps in physiology and pathophysiology. Nat. Rev. Mol. Cell Biol. 8, 917–929 (2007).

42. Walker, J. E., Saraste, M., Runswick, M. J. & Gay, N. J. Distantly related sequences in the alpha- and beta-subunits of ATP synthase, myosin, kinases and other ATP-requiring enzymes and a common nucleotide binding fold. The EMBO Journal vol. 1 945–951 Preprint at https://doi.org/10.1002/j.1460-2075.1982.tb01276.x (1982).

43. Schwartz, R. M. & Dayhoff, M. O. Origins of prokaryotes, eukaryotes, mitochondria, and chloroplasts. Science 199, 395–403 (1978).

44. Feng, Y. et al. The Evolutionary Origins of Extreme Halophilic Archaeal Lineages. Genome Biol. Evol. 13, (2021).

45. Szöllosi, G. J., Boussau, B., Abby, S. S., Tannier, E. & Daubin, V. Phylogenetic modeling of lateral gene transfer reconstructs the pattern and relative timing of speciations. Proc. Natl. Acad. Sci. U. S. A. 109, 17513–17518 (2012).

46. Szöllõsi, G. J., Rosikiewicz, W., Boussau, B., Tannier, E. & Daubin, V. Efficient exploration of the space of reconciled gene trees. Syst. Biol. 62, 901–912 (2013).

47. Morel, B. et al. SpeciesRax: A Tool for Maximum Likelihood Species Tree Inference from Gene Family Trees under Duplication, Transfer, and Loss. Mol. Biol. Evol. 39, (2022).

48. Hochberg, G. K. A. & Thornton, J. W. Reconstructing Ancient Proteins to Understand the Causes of Structure and Function. Annu. Rev. Biophys. 46, 247–269 (2017).

49. Sharma, P. P. & Wheeler, W. C. Cross-bracing uncalibrated nodes in molecular dating improves congruence of fossil and molecular age estimates. Frontiers in Zoology vol. 11 Preprint at https://doi.org/10.1186/s12983-014-0057-x (2014).

50. Szöllősi, G. J. et al. Relative time constraints improve molecular dating. Systematic Biology Preprint at https://doi.org/10.1093/sysbio/syab084 (2021).

51. Lapierre, P., Shial, R. & Peter Gogarten, J. Distribution of F- and A/V-type ATPases in Thermus scotoductus and other closely related species. Systematic and Applied Microbiology vol. 29 15–23 Preprint at https://doi.org/10.1016/j.syapm.2005.06.004 (2006).

52. Sumi, M., Yohda, M., Koga, Y. & Yoshida, M. F0F1-ATPase genes from an archaebacterium, Methanosarcina barkeri. Biochem. Biophys. Res. Commun. 241, 427–433 (1997).

53. Saum, R., Schlegel, K., Meyer, B. & Müller, V. The F1FO ATP synthase genes in Methanosarcina acetivorans are dispensable for growth and ATP synthesis. FEMS Microbiol. Lett. 300, 230–236 (2009).

54. Dibrova, D. V., Galperin, M. Y. & Mulkidjanian, A. Y. Characterization of the N-ATPase, a distinct, laterally transferred Na+-translocating form of the bacterial F-type membrane ATPase. Bioinformatics 26, 1473–1476 (2010).

55. Castelle, C. J. & Banfield, J. F. Major New Microbial Groups Expand Diversity and Alter our Understanding of the Tree of Life. Cell 172, 1181–1197 (2018).

56. Castelle, C. J. et al. Biosynthetic capacity, metabolic variety and unusual biology in the CPR and DPANN radiations. Nat. Rev. Microbiol. 16, 629–645 (2018).

57. Dombrowski, N., Lee, J.-H., Williams, T. A., Offre, P. & Spang, A. Genomic diversity, lifestyles and evolutionary origins of DPANN archaea. FEMS Microbiol. Lett. 366, (2019).

58. Wang, B. et al. Expansion of Thaumarchaeota habitat range is correlated with horizontal transfer of ATPase operons. ISME J. 13, 3067–3079 (2019).

59. Evans, P. N. et al. Methane metabolism in the archaeal phylum Bathyarchaeota revealed by genome-centric metagenomics. Science 350, 434–438 (2015).

60. Evans, P. N. et al. An evolving view of methane metabolism in the Archaea. Nat. Rev. Microbiol. 17, 219–232 (2019).

61. Stairs, C. W., Leger, M. M. & Roger, A. J. Diversity and origins of anaerobic metabolism in mitochondria and related organelles. Philos. Trans. R. Soc. Lond. B Biol. Sci. 370, 20140326 (2015).

62. Gawryluk, R. M. R. & Stairs, C. W. Diversity of electron transport chains in anaerobic protists. Biochim. Biophys. Acta Bioenerg. 1862, 148334 (2021).

63. Hilario, E. & Gogarten, J. P. The Prokaryote-to-Eukaryote Transition Reflected in the Evolution of the V/F/A-ATPase Catalytic and Proteolipid Subunits. Journal of Molecular Evolution vol. 46 703– 715 Preprint at https://doi.org/10.1007/pl00006351 (1998).

64. Williams, T. A. et al. Integrative modeling of gene and genome evolution roots the archaeal tree of life. Proc. Natl. Acad. Sci. U. S. A. 114, E4602–E4611 (2017).

65. Cavalier-Smith, T. Rooting the tree of life by transition analyses. Biol. Direct 1, 19 (2006).

66. Lake, J. A., Skophammer, R. G., Herbold, C. W. & Servin, J. A. Genome beginnings: rooting the tree of life. Philos. Trans. R. Soc. Lond. B Biol. Sci. 364, 2177–2185 (2009).

67. Gouy, R., Baurain, D. & Philippe, H. Rooting the tree of life: the phylogenetic jury is still out. Philos. Trans. R. Soc. Lond. B Biol. Sci. 370, 20140329 (2015).

68. Mascotti, M. L. Resurrecting Enzymes by Ancestral Sequence Reconstruction. Methods Mol. Biol. 2397, 111–136 (2022).

69. Saraste, M., Sibbald, P. R. & Wittinghofer, A. The P-loop--a common motif in ATP- and GTP-binding proteins. Trends Biochem. Sci. 15, 430–434 (1990).

70. Leipe, D. D., Wolf, Y. I., Koonin, E. V. & Aravind, L. Classification and evolution of P-loop GTPases and related ATPases. J. Mol. Biol. 317, 41–72 (2002).

71. Schäfer, I. B. et al. Crystal Structure of the Archaeal A1AO ATP Synthase Subunit B from Methanosarcina mazei Gö1: Implications of Nucleotide-binding Differences in the Major A1AO Subunits A and B. Journal of Molecular Biology vol. 358 725–740 Preprint at https://doi.org/10.1016/j.jmb.2006.02.057 (2006).

72. Ponce-Toledo, R. I. et al. An Early-Branching Freshwater Cyanobacterium at the Origin of Plastids. Curr. Biol. 27, 386–391 (2017).

73. Harris, B. J. et al. Divergent evolutionary trajectories of bryophytes and tracheophytes from a complex common ancestor of land plants. Nat Ecol Evol 6, 1634–1643 (2022).

74. Nelson-Sathi, S. et al. Origins of major archaeal clades correspond to gene acquisitions from bacteria. Nature 517, 77–80 (2015).

75. Lane, N., Allen, J. F. & Martin, W. How did LUCA make a living? Chemiosmosis in the origin of life. Bioessays 32, 271–280 (2010).

76. Sojo, V., Pomiankowski, A. & Lane, N. A bioenergetic basis for membrane divergence in archaea and bacteria. PLoS Biol. 12, e1001926 (2014).

77. Koga, Y. Early evolution of membrane lipids: how did the lipid divide occur? J. Mol. Evol. 72, 274– 282 (2011).

78. Villanueva, L., Schouten, S. & Damsté, J. S. S. Phylogenomic analysis of lipid biosynthetic genes of Archaea shed light on the “lipid divide.” Environ. Microbiol. 19, 54–69 (2017).

79. Villanueva, L. et al. Bridging the membrane lipid divide: bacteria of the FCB group superphylum have the potential to synthesize archaeal ether lipids. ISME J. 15, 168–182 (2021).

80. Lombard, J. & Moreira, D. Origins and early evolution of the mevalonate pathway of isoprenoid biosynthesis in the three domains of life. Mol. Biol. Evol. 28, 87–99 (2011).

81. Hoshino, Y. & Gaucher, E. A. On the Origin of Isoprenoid Biosynthesis. Molecular Biology and Evolution vol. 35 2185–2197 Preprint at https://doi.org/10.1093/molbev/msy120 (2018).

82. Lombard, J., López-García, P. & Moreira, D. The early evolution of lipid membranes and the three domains of life. Nat. Rev. Microbiol. 10, 507–515 (2012).

83. Mohanty, S. et al. Structural basis for a unique ATP synthase core complex from Nanoarcheaum equitans. J. Biol. Chem. 290, 27280–27296 (2015).

84. Krause, S., Bremges, A., Münch, P. C., McHardy, A. C. & Gescher, J. Characterisation of a stable laboratory co-culture of acidophilic nanoorganisms. Sci. Rep. 7, 3289 (2017).

85. Narasingarao, P. et al. De novo metagenomic assembly reveals abundant novel major lineage of Archaea in hypersaline microbial communities. ISME J. 6, 81–93 (2012).

86. Andrade, K. et al. Metagenomic and lipid analyses reveal a diel cycle in a hypersaline microbial ecosystem. The ISME Journal vol. 9 2697–2711 Preprint at https://doi.org/10.1038/ismej.2015.66 (2015).

87. Dombrowski, N. et al. Undinarchaeota illuminate DPANN phylogeny and the impact of gene transfer on archaeal evolution. Nat. Commun. 11, 3939 (2020).

88. Aouad, M. et al. A divide-and-conquer phylogenomic approach based on character supermatrices resolves early steps in the evolution of the Archaea. BMC Ecol Evol 22, 1 (2022).

89. Zhu, Q. et al. Phylogenomics of 10,575 genomes reveals evolutionary proximity between domains Bacteria and Archaea. Nat. Commun. 10, 5477 (2019).

90. Berney, C. & Pawlowski, J. A molecular time-scale for eukaryote evolution recalibrated with the continuous microfossil record. Proc. Biol. Sci. 273, 1867–1872 (2006).

91. Chernikova, D., Motamedi, S., Csürös, M., Koonin, E. V. & Rogozin, I. B. A late origin of the extant eukaryotic diversity: divergence time estimates using rare genomic changes. Biol. Direct 6, 26 (2011).

92. Strassert, J. F. H., Irisarri, I., Williams, T. A. & Burki, F. A molecular timescale for eukaryote evolution with implications for the origin of red algal-derived plastids. Nat. Commun. 12, 1879 (2021).

93. Tang, Q., Pang, K., Yuan, X. & Xiao, S. A one-billion-year-old multicellular chlorophyte. Nat Ecol Evol 4, 543–549 (2020).

94. Gibson, T. M., et al. Precise age of Bangiomorpha pubescens dates the origin of eukaryotic photosynthesis. Geology vol. 46 135–138 Preprint at https://doi.org/10.1130/g39829.1 (2018).

95. Philippe, H. & Forterre, P. The rooting of the universal tree of life is not reliable. J. Mol. Evol. 49, 509–523 (1999).

96. Pittis, A. A. & Gabaldón, T. Late acquisition of mitochondria by a host with chimaeric prokaryotic ancestry. Nature 531, 101–104 (2016).

97. Martin, W. F. et al. Late Mitochondrial Origin Is an Artifact. Genome biology and evolution vol. 9 373–379 (2017).

98. Vosseberg, J. et al. Timing the origin of eukaryotic cellular complexity with ancient duplications. Nat Ecol Evol 5, 92–100 (2021).

99. Martijn, J., Vosseberg, J., Guy, L., Offre, P. & Ettema, T. J. G. Deep mitochondrial origin outside the sampled alphaproteobacteria. Nature 557, 101–105 (2018).

100. Muñoz-Gómez, S. A. et al. Site-and-branch-heterogeneous analyses of an expanded dataset favour mitochondria as sister to known Alphaproteobacteria. Nat Ecol Evol 6, 253–262 (2022).

101. Criscuolo, A. & Gribaldo, S. BMGE (Block Mapping and Gathering with Entropy): a new software for selection of phylogenetic informative regions from multiple sequence alignments. BMC Evol. Biol. 10, 210 (2010).

102. Nguyen, L.-T., Schmidt, H. A., von Haeseler, A. & Minh, B. Q. IQ-TREE: a fast and effective stochastic algorithm for estimating maximum-likelihood phylogenies. Mol. Biol. Evol. 32, 268–274 (2015).

103. Hoang, D. T., Chernomor, O., von Haeseler, A., Minh, B. Q. & Vinh, L. S. UFBoot2: Improving the Ultrafast Bootstrap Approximation. Mol. Biol. Evol. 35, 518–522 (2018).

104. Minh, B. Q., et al. Corrigendum to: IQ-TREE 2: New Models and Efficient Methods for Phylogenetic Inference in the Genomic Era. Mol. Biol. Evol. 37, 2461 (2020).

105. Burki, F., Roger, A. J., Brown, M. W. & Simpson, A. G. B. The New Tree of Eukaryotes. Trends Ecol. Evol. 35, 43–55 (2020).

106. Kalyaanamoorthy, S., Minh, B. Q., Wong, T. K. F., von Haeseler, A. & Jermiin, L. S. ModelFinder: fast model selection for accurate phylogenetic estimates. Nat. Methods 14, 587–589 (2017).

107. Capella-Gutiérrez, S., Silla-Martínez, J. M. & Gabaldón, T. trimAl: a tool for automated alignment trimming in large-scale phylogenetic analyses. Bioinformatics 25, 1972–1973 (2009).

108. Katoh, K., Misawa, K., Kuma, K.-I. & Miyata, T. MAFFT: a novel method for rapid multiple sequence alignment based on fast Fourier transform. Nucleic Acids Res. 30, 3059–3066 (2002).

109. Utter, D. R., He, X., Cavanaugh, C. M., McLean, J. S. & Bor, B. The saccharibacterium TM7x elicits differential responses across its host range. ISME J. 14, 3054–3067 (2020).

110. He, X. et al. Cultivation of a human-associated TM7 phylotype reveals a reduced genome and epibiotic parasitic lifestyle. Proc. Natl. Acad. Sci. U. S. A. 112, 244–249 (2015).

111. Aramaki, T. et al. KofamKOALA: KEGG Ortholog assignment based on profile HMM and adaptive score threshold. Bioinformatics 36, 2251–2252 (2020).

112. Tatusov, R. L., Koonin, E. V. & Lipman, D. J. A Genomic Perspective on Protein Families. Science vol. 278 631–637 Preprint at https://doi.org/10.1126/science.278.5338.631 (1997).

113. Galperin, M. Y., Kristensen, D. M., Makarova, K. S., Wolf, Y. I. & Koonin, E. V. Microbial genome analysis: the COG approach. Brief. Bioinform. 20, 1063–1070 (2019).

114. Galperin, M. Y. et al. COG database update: focus on microbial diversity, model organisms, and widespread pathogens. Nucleic Acids Res. 49, D274–D281 (2021).

115. Bateman, A. et al. The Pfam protein families database. Nucleic Acids Res. 32, D138–41 (2004).

116. Finn, R. D., Clements, J. & Eddy, S. R. HMMER web server: interactive sequence similarity searching. Nucleic Acids Res. 39, W29–37 (2011).

117. Eddy, S. R. A new generation of homology search tools based on probabilistic inference. Genome Inform. 23, 205–211 (2009).

118. Eddy, S. R. Accelerated Profile HMM Searches. PLoS Comput. Biol. 7, e1002195 (2011).

119. Li, W. & Godzik, A. Cd-hit: a fast program for clustering and comparing large sets of protein or nucleotide sequences. Bioinformatics vol. 22 1658–1659 Preprint at https://doi.org/10.1093/bioinformatics/btl158 (2006).

120. Fu, L., Niu, B., Zhu, Z., Wu, S. & Li, W. CD-HIT: accelerated for clustering the next-generation sequencing data. Bioinformatics 28, 3150–3152 (2012).

121. Price, M. N., Dehal, P. S. & Arkin, A. P. FastTree 2 – Approximately Maximum-Likelihood Trees for Large Alignments. PLoS ONE vol. 5 e9490 Preprint at https://doi.org/10.1371/journal.pone.0009490 (2010).

122. Shimodaira, H. An approximately unbiased test of phylogenetic tree selection. Syst. Biol. 51, 492– 508 (2002).

123. UniProt Consortium. UniProt: a worldwide hub of protein knowledge. Nucleic Acids Res. 47, D506– D515 (2019).

124. Mitchell, A. L., et al. InterPro in 2019: improving coverage, classification and access to protein sequence annotations. Nucleic Acids Res. 47, D351–D360 (2019).

125. Buchfink, B., Xie, C. & Huson, D. H. Fast and sensitive protein alignment using DIAMOND. Nat. Methods 12, 59–60 (2015).

126. Bergsten, J. A review of long-branch attraction. Cladistics vol. 21 163–193 Preprint at https://doi.org/10.1111/j.1096-0031.2005.00059.x (2005).

127. Waterhouse, A. M., Procter, J. B., Martin, D. M. A., Clamp, M. & Barton, G. J. Jalview Version 2--a multiple sequence alignment editor and analysis workbench. Bioinformatics 25, 1189–1191 (2009).

128. Crooks, G. E., Hon, G., Chandonia, J.-M. & Brenner, S. E. WebLogo: a sequence logo generator. Genome Res. 14, 1188–1190 (2004).

129. Eme, L. et al. Inference and reconstruction of the heimdallarchaeial ancestry of eukaryotes. Preprint at https://doi.org/10.1101/2023.03.07.531504.

130. Shimodaira, H. & Hasegawa, M. CONSEL: for assessing the confidence of phylogenetic tree selection. Bioinformatics 17, 1246–1247 (2001).

