## Supplementary Material for "ATP synthase evolution on a cross-braced dated tree of life"

Phylogenetic placement of eukaryotic catalytic (A) and non-catalytic (B) F-type ATP synthase homologs 4

#### Supplementary Discussion

##### The ATP synthase and Walker A motifs

The ATP synthase is present across all domains of life[1–4] and shares a common foundational architecture amongst its variants. In particular, the F- and A/V-type ATP synthases are comprised of a soluble hetero-hexameric head, a central stalk, and peripheral components (R1), which are connected to an insoluble membrane-embedded ring complex (R0). The hexameric head of the R1 complex contains three copies each of a catalytic and non-catalytic/regulatory subunit, and is the site of ATP binding, synthesis/hydrolysis. Both the F- and A-type ATP synthases couple chemiosmotic membrane potential to the synthesis of ATP, whereas eukaryotic V-type complexes hydrolyze ATP to generate a proton gradient that allows for acidification of the vacuolar compartment[1,5]. All three subtypes have been shown to be reversible (i.e. functioning as a synthase and/or hydrolase) *in vitro,* however, *in vivo* the eukaryotic V-type ATPase is limited to hydrolytic activity[5]).

Differences in the catalytic potential of the soluble R1 component of F- and A/V-type ATP synthases have been based on the amino acid in the third position in the Walker A motif which differs between the F- and A/V-type subunits, with the former containing an alanine (A) and the latter containing a phenylalanine (F) in the third position[6]. Though alanine and phenylalanine share equivalencies as amino acids, they appear to play different roles in the distinct types of ATP synthases. The phenylalanine (F) of A/V-type complexes stabilizes the P-loop while the alanine (A) of F-type complexes orients itself toward the terminal phosphate during ATP catalysis[6]. This disparity results in varying conformational arrangements of the P-loop, where the bound nucleotide has weaker interactions inside the A1/V1A (*c*A1V1) subunit (compared to the F1-beta/*c*F1), making it more exposed to solvents[7] which also lends to the A/V-type ATP synthase's ability to hydrolyze GTP and UTP via a larger P-loop region[6]. Additionally, although yet to be confirmed, it is possible that the alternative binding motif in the A1/V1B (*nc*A1V1) subunit, might play a role in the alternative positioning of the P-loop and binding of phosphate-containing moieties, whereas the equivalent binding process is tightly constrained in the F-type ATP synthase due to conservation of the Walker A motif in both the catalytic and non-catalytic subunits.

##### Phylogenetic placement of eukaryotic catalytic (A) and non-catalytic (B) A/V-type ATP synthases

The highest support for the relationship between Asgard archaea and eukaryotes was observed in trees containing non-catalytic A/V-type subunits (Supplementary Figures 7, 8, and 10; Supplementary Table 8). We observed eukaryotes placing with members of Asgard and TACK in the *nc*A1V1 phylogeny (bs support = 95.8/95, Supplementary Figure 7, Supplementary Table 8) and the phylogeny containing *nc*A1V1 rooted with the *nc*F1 subunit (bs support = 87.2/92, Supplementary Figure 8, Supplementary Table 8). Eukaryotic homologs of the catalytic A/V subunits branched sister to poorly resolved groups containing members of the Deinococcus-Thermus and Synergistetes in the *c*A1V1 phylogeny (bs support = 50.6/66, Supplementary Figure 6, Supplementary Table 8), in contrast the eukaryotic sequences clustered sister to an unresolved group of Thor- and Odinarchaeota in the phylogeny containing the *c*A1V1 rooted with *c*F1 (bs support = 0/13, Supplementary Figure 9, Supplementary Table 8). Finally, in the phylogeny that comprises all homologs of both F1- and A1/V1-type ATP synthase subunits, eukaryotic *nc*A1V1 homologs cluster sister to the Heimdallarchaeota (bs support = 91.4/56, Supplementary Figure 10, Supplementary Table 8, Data repository: 10.5281/zenodo.7807739), but the *c*A1V1 eukaryotic homologs were sister to a mixed-group of TACK and Euryarchaeota, and various bacterial lineages including the Deinococcus-Thermus and Synergistetes (bs support = 84.8/65, Supplementary Figure 10, Supplementary Table 8, Data repository: 10.5281/zenodo.7807739). These findings indicate that the non-catalytic A1/V1B subunit has retained more information to resolve the placement of eukaryotic homologs within Archaea than the catalytic subunit, which may experience stronger selective constraints.

**Phylogenetic placement of eukaryotic catalytic (A) and non-catalytic (B) F-type ATP synthase homologs**

To test whether the placement of the various eukaryotic F1-type ATP synthase homologs could be resolved using Bayesian approaches, we constructed smaller datasets for placing eukaryotic *c* and *nc* F1 type subunits (see methods) relative to alphaproteobacterial and cyanobacterial homologs. Using outgroup rooting with Gammaproteobacteria and Melainabacteria, respectively, the Bayesian phylogenies revealed a similar topology to what was observed in the larger protein trees inferred using maximum likelihood approaches. The mitochondrial datasets recovered various multifurcations, supporting the observation that the signal harbored within the *c* and *nc* F1 subunits is insufficient to unambiguously identify the alphaproteobacterial donor lineage. Consistent with the larger maximum-likelihood based protein phylogenies (Supplementary Figures 4, 5, 8, 9, and 10), the eukaryote-*Gloeomargarita lithophora* sisterhood relationship was obtained in the *nc*F1 phylogeny (Supplementary Figure 11a), while the phylogeny of the *c*F1 could not resolve the placement of eukaryotes within the Cyanobacteria (Supplementary Figure 11b).

**Deep origins of the F- and A/V-type ATP synthases**

It is worth noting that mapping the presence of a gene to deep nodes on the TOL is challenging, and we cannot exclude the presence of a single ATP synthase in LUCA[8–10] and the acquisition of the A/V-type ATP synthase in LBCA or its early descendents (Supplementary Table 6). Prospective analyses could focus on reconciling a larger set of genes within the TOL to reconstruct the gene content of LUCA using gene tree-species tree reconciliations and integrate data on lipid evolution. This will not only help to further unravel the features of the ancestor of Archaea and Bacteria but also help guide hypotheses on the evolutionary forces underlying the deep split between the primary domains of life and their key distinguishing features including membrane lipids and biogeochemistry.

**Consistency of our dating results with recent literature**

A number of recent studies have estimated the ages of the major prokaryotic lineages, using a range of different calibration strategies and molecular clock approaches[11–16]. Our species tree dating analysis is generally in broadly agreement with these studies, although the 95% highest posterior density (HPD) regions remain broad and it is clear that substantial uncertainty remains. In our analysis, the 95% HPDs for the radiation of oxygenic Cyanobacteria and Proteobacteria are consistent with an origin that overlaps with the Great Oxidation Event (GOE, Figure 5A). Whether crown Oxyphotobacteria (photosynthetic Cyanobacteria) radiated before, after, or during the GOE remains the subject of some debate[12,13,17]. The 95% HPD of our estimate (2.74-2.03 Ga) is sufficiently broad to admit any of these possibilities (Figure 5A).

It is slightly younger than some recent analyses that confidently place the radiation of Oxyphotobacteria prior to the GOE[13,18], but older than others that place the radiation of Oxyphotobacteria at 2 Ga or more recently[19]. Comparative genomics suggests that the common ancestor of Oxyphotobacteria was likely capable of both oxygenic photosynthesis and aerobic respiration [11,20], which is consistent with a post-GOE radiation of the group (that is, that oxygenic photosynthesis evolved on the Oxyphotobacteria stem, and that oxygen concentrations were high enough to favor the retention of oxygen reductases by the time of the crown ancestor). Our estimate for the age of Alphaproteobacteria and Eukaryotes (2.58-2.12 Ga, Figure 5A, Figure 5C, Supplementary Figure 16, Supplementary Table 10) is slightly older than a recent analysis based on mitochondrial proteins (ca. 2.1-1.7 Ga (mean ca. 1.9 Ga))[14]. In either case, a radiation of crown Alphaproteobacteria after the GOE is consistent with the inference that their common ancestor was, based on its ancestrally inferred gene content, a facultative aerobic organism[21].

**Supplementary methods regarding the use of a bracing approach to improve absolute dating of the Tree of Life and ATP synthase phylogenies**

Relative constraints and braced nodes allow to propagate fossil information to poorly-constrained regions of the tree, and also to provide additional dating information for braced or constrained nodes of unknown age. For dating ATP synthase evolution and the tree of life, we used the gene tree presented in Figure 3 (Supplementary Fig 10) and Supplementary Figure 13, respectively. Maximum likelihood phylogenies inferred from these datasets contained some relationships (in particular, internal relationships within the eukaryotes) that, based on external evidence, are likely incorrect and prevented the assignment of some calibrations to the tree (for example, when a well-established group is not recovered as monophyletic). We therefore edited the ML topology (Supplementary Figure 21) to allow these calibrations to be applied (see Methods). As the position of eukaryotes within Archaea [22,23] and Bacteria [24–26] remains debated, we also performed an experiment in which we fixed the nuclear and mitochondrial clades in two different positions (eukaryotes within, or sister to, Asgard archaea; and as sister to Alphaproteobacteria or branching within them - see Supplementary Figures 12-13). The results discussed in the main text are taken from our focal analysis in which eukaryotes branched sister to Hodarchaeota (formerly Heimdall LC3) within Asgards, and as sister to all Alphaproteobacteria (excluding Magnetococcales) (Edited2, see Methods; Figure 5; Supplementary Figures 13 and 16, Supplementary Table 10). Results were generally similar across all analyses, and we discuss the cases where they disagree. The fossil calibrations and cross-braced nodes used in these analyses are summarized in Supplementary Table 9. We also used a single relative age constraint: that the common ancestor of plastids must be younger than nuclear and mitochondrial LECA. Interestingly, an AU test comparing the likelihood of the three topologies: maximum likelihood tree, focal analysis and secondary tree (with alternative placement of eukaryotes) indicated that the two constrained trees were strongly rejected by the alignment and model (Edited1, P = 1.65e-07; Edited2 P = 9.54e-08, AU test; Supplementary Table 18). This result highlights the difficulty of resolving phylogenetic relationships at different depths in a single analysis of 12 ribosomal proteins (2133 amino acid sites).

Dating prokaryotic clades is challenging due to the lack of calibrations, and differences in age estimates among published analyses reflect different choices about which calibrations, sequence data and molecular clock methods to use. While our dating inferences are broadly consistent with previous analyses, some differences might be explained by our use of cross-bracing and relative constraints to inform rate estimates within otherwise poorly-calibrated regions of the tree. For example, cross-bracing the nuclear and mitochondrial LECA nodes may support older ages for lineages ancestral to mitochondria due to the slower "ticking" of the clock with the nuclear clade. Younger ages of crown Oxyphotobacteria[12], likely result from root age differences. In Shih et al. (2017)[12], LBCA was calibrated to 2.4-3.8Ga, which. This is substantially younger than the age estimated for LBCA in our analysis (4.49-4.05Ga) and is likely due to the the antecedent LUCA-root calibrated using the moon-forming impact 4.52Ga), which may explain their younger inferred ages for descendant clades including Oxyphotobacteria.

#### Fossil Calibrations

**LUCA | 3347-4520 Ma**

**Fossil taxon and specimen:** Strelley Pool Formation, Pilbara Craton, Following the justification outlined in Betts et al. (2018).

**Minimum Age justification:** The minimum age of the Strelley Pool Formation is 3.350 Ga ± 0.003 Gyr based on a volcaniclastic tuff, at the base of the overlying Euro Basalt[27] in the Kelly Group. Hence our minimum age constraint is 3.347 Ga.

**Hard maximum age justification:** The Moon forming impact would have effectively sterilized the Earth and so it serves as an effective basis for establishing a hard maximum age constraint on LUCA. Pb-Pb dating carried out on Moon rocks, yielding a date of 4.51 Ga ± 10 Myr[28] a date which has also recently been confirmed by reanalysis of the Apollo mission zircons[29]. Thus, our maximum constraint is 4.52 Ga.

**Total group Oxyphotobacteria | 3225-4520 Ma**

**Clade:** This equates to the traditional concept of cyanobacteria, excluding Melainabacteria[12], but including the stem to this remaining clade.

**Fossil taxon and specimen**: stable Fe and U-Th-Pb isotopes in the Manzimnyama Banded Ironstone Formation, Fig Tree Group, Barberton, South Africa[30].

**Hard Minimum Age**: 3225 Ma

**Hard Maximum Age**: 4520 Ma

**Discussion:** there are claims and counterclaims for life in the Hadean and Archaean and while we accept that there is credible fossil, sedimentologic and isotopic evidence for life by 3347 Ma (see Betts et al. 2018), there has been insufficient consideration of whether these records evidence the establishment of the crown clade of life - i.e., divergence from LUCA. Implicitly or explicitly, most records have been attributed to the crown clade of life based on claims, direct or indirect, of oxygenic cyanobacteria, either as microfossils, sedimentary structures such as MISS or stromatolites, or Banded Ironstones. The existence of stromatolites would appear to provide evidence of phototactic bacteria, but they might otherwise represent microbial organisms competing for other nutrients within the water colum [31]. Similarly, banded ironstones can be formed by reaction with oxygen from abiogenic sources like photolysis[32], hence, banded ironstones only become a significant proxy for life when they occur in volume (and even then they are not necessarily linked to oxygenic photosynthesis). Therefore we follow Betts et al. (2018) who based their calibration minimum for total-group cyanobacteria on Satkoski et al. (2015) who presented evidence of stable Fe and U-Th-Pb isotopes in the Manzimnyama Banded Ironstone Formation (Fig Tree Group, Barberton, South Africa), indicating a level of free oxygen indicative of cyanobacterial activity[17,30].

**Hard minimum age justification**: (from Betts et al. 2018) The isotopic evidence from the Manzimnyama BIF in the Fig Tree Group, Barberton, South Africa[30]. The age of the Fig Tree Group is well constrained with a spherule layer at its base dated at 3258 Ma ± 3 Myr[33], and an overlying volcanic unit at its top dated at 3226 Ma ± 1 Myr[34]. Hence, the minimum date we would assign is 3225 Myr.

**Hard maximum age justification**: The Moon forming impact would have effectively sterilized the Earth and so it serves as an effective basis for establishing a hard maximum age constraint. Pb-Pb dating carried out on Moon rocks, yielding a date of 4.51 Ga ± 10 Myr[28] a date which has also recently been confirmed by reanalysis of the Apollo mission zircons[29]. Thus, our maximum constraint is 4.52 Ga.

**Crown Oxyphotobacteria | 2013.6-3448 Ma**

**Clade:** This equates to the extant clade of cyanobacteria, excluding Melainabacteria[12].

**Fossil taxon and specimen**: *Eoentophysalis belcherensis*. (Holotype) GSC type no. 42770, from the upper part of Kasegalik Formation, Belcher Supergroup, unnamed island at north end of Churchill Sound (Locality A of Hofmann, 1976).

**Phylogenetic justification**: Compared to extant *Entophysalis* (Chroococcales) by[35] based on its similarly sized coccoidal cells, colonial organization resulting from binary fission in three perpendicular planes, and warty (pustular) mamilate-shaped outer layers[35]. These characteristics are convincing of cyanobacterial affinity, however, on the basis of such necessarily limited evidence, that *Eoentophysalis belcherensis* is a close relative of extant *Entophysalis*, not least since so little of traditional cyanobacterial taxonomy (based in cytological, cell division patterns and arrangements) has been corroborated by molecular phylogenetics. Indeed, we cannot be certain that *Eoentophysalis belcherensis* is a member of crown-Oxyphotobacteria, however, evidence of its phenotype and development is sufficient to conclude with high probability that it is a member of crown-Oxyphotobacteria and, on that basis, we use it as a basis for establishing a soft minimum constraint. The alternative would be to rely on *Bangiomorpha pubescens* but this inference of a eukaryote plastid is, by its very nature, remote from the timing of origin of crown-Oxyphotobacteria.

**Soft minimum age justification:** The age of the Kasegalik Formation (Belcher Supergroup) has been established on the basis of U-Pb dating of tufts near the base and top[36]. The latter is dated to 2015.4 Ma ± 1.8 Myr, providing for a 2013.6 Ma minimum constraint on the oldest record of *Eoentophysalis belcherensis* and, therefore, crown-Cyanobacteria.

**Soft maximum age justification:** Claims of Oxyphotobacteria in the Strelley Pool Formation can be rationalised in other ways (see total-group Bacteria, above). Will implement a soft maximum to allow for the possibility that Oxyphotobacteria were established prior to the deposition of the Strelley Pool Formation. The maximum depositional age of the Strelley Pool Formation has been dated to 3414 ± 34 Ma[37], yielding a maximum age interpretation of 3448 Ma.

**Total-group Chromatiaceae | 1631-4520 Ma**

**Fossil taxon and specimen**: Okenane biomarker record from the Barney Creek Formation, McArthur Basin, Australia[38].

**Phylogenetic justification**: Okenane is described as specific to Chromatiaceae (Brocks and Schaeffer, 2008)[13] contend that it may have a more general distribution among Chlorobiaceae but this view is based solely on incompatibility among the calibrations used in that study.
**Soft minimum age justification:** The Barney Creek Formation has recently been dated to 1642.2 Ma ± 3.9 Myr, but this is from a sample close to the base of the Formation[39]. Page and Sweet (1998) provide dates from tufts higher in the sequence, the youngest of which is 1638 Ma ± 7 Myr, providing for a minimum constraint of 1631 Ma[40].
**Hard maximum age justification:** The Moon forming impact would have effectively sterilized the Earth and so it serves as an effective basis for establishing a hard maximum age constraint. Pb-Pb dating carried out on Moon rocks, yielding a date of 4.51 Ga ± 10 Myr[28] a date which has also recently been confirmed by reanalysis of the Apollo mission zircons[29]. Thus, our maximum constraint is 4.52 Ga.

**Total Group Eukaryota | 1619.1-4520 Ma**
**Fossil taxon and specimen:** Changzhougou Formation, North China, following the justification outlined in Betts et al. (2018).
**Minimum Age justification:** The minimum age of the Changzhougou Formation is established based on dated ashes in the overlying Chuanlinggou Formation, dated to 1625.3 ± 6.2 Myr[41], thus 1619.1 Ma.
**Hard maximum age justification:** The Moon forming impact would have effectively sterilized the Earth and so it serves as an effective basis for establishing a hard maximum age constraint on LUCA. Pb-Pb dating carried out on Moon rocks, yielding a date of 4.51 Ga ± 10 Myr[28] a date which has also recently been confirmed by reanalysis of the Apollo mission zircons[29]. Thus, our maximum constraint is 4.52 Ga.

**Crown Group Eukaryota | 1030-1879.6 Ma**
**Fossil taxon and specimen:** *Bangiomorpha pubescens* (HUPC 62912; Slide HUST-1A, England Finder coordinates: O-35; Paleobotanical Collections of Harvard University, USA) from the Hunting Formation in northwestern Somerset Island, Arctic Canada[42].
**Phylogenetic justification:** Morphological similarity of *Bangiomorpha pubescens* with modern red algae *Bangia*[42] evidences affinity to total group Rhodophyta; this is the oldest unequivocal record of a crown-eukaryote.
**Minimum age justification:** Re-Os isotopic dating of sedimentary rocks in the stratigraphic region in which *Bangiomorpha pubescens* was sampled date this fossil at 1.047 +0.013/–0.017 Ga, yielding a minimum age constraint of 1030 Ma[43].
**Hard maximum age justification:** The Gunflint Chert microflora has a long history of study, including claims of eukaryotes. These include process-bearing acritarch-like cysts, such as *Germinosphaera*, suggesting the presence of an actin cytoskeleton. Nevertheless, all such claims of eukaryote affinity have generally been rejected[44]. The Gunflint Chert has been dated to 1878.3 Ma ± 1.3 Myr[45], yielding a maximum age of 1879.6 Ma.

**Total group Rhodophyta | 1030-1879.6 Ma**
**Fossil taxon and specimen:** *Bangiomorpha pubescens* (HUPC 62912; Slide HUST-1A, England Finder coordinates: O-35; Paleobotanical Collections of Harvard University, USA) from the Hunting Formation in northwestern Somerset Island, Arctic Canada[42].

**Phylogenetic justification:** Morphological similarity of *Bangiomorpha pubescens* with modern red algae *Bangia*[42].
**Minimum age justification:** Re-Os isotopic dating of sedimentary rocks in the stratigraphic region in which *Bangiomorpha pubescens* was sampled date this fossil at 1.047 +0.013/–0.017 Ga, yielding a minimum age constraint of 1030 Ma[43].
**Hard maximum age justification:** The Gunflint Chert microflora has a long history of study, including claims of eukaryotes. These include process-bearing acritarch-like cysts, such as *Germinosphaera*, suggesting the presence of an actin cytoskeleton. Nevertheless, all such claims of eukaryote affinity have generally been rejected[44]. The Gunflint Chert has been dated to 1878.3 Ma ± 1.3 Myr[45], yielding a maximum age of 1879.6 Ma.

**Total-group Eumycota *Spizellomyces – Saccharomyces* 879-1879.6 Ma**
**Fossil taxon and specimen:** *Ourasphaira giraldae* (Specimen 74639-W46,3, sample 15RAT-021A1 in the collections of the Department of Geology in the University of Liège, Belgium) from the Shale of the Grassy Bay Formation in the Brock Inlier, Northwestern Territories, Canada[46].
**Phylogenetic justification:** Loron et al. (2019b)[47] establish a eukaryote affinity on the basis of a 'combination of complex morphology, right-angle branching, multicellularity, bilayered wall ultrastructure, compositional recalcitrance and relatively large size', evidencing the presence of a complex cytoskeleton; combined with their FTIR spectroscopy which diagnoses the presence of chitin, they conclude a total-group Dikarya affinity for *Ourasphaira giraldae*.

**Minimum Age justification:** There are no direct dates for the Grassy Bay Formation but the overlying Boot Inlet Formation has been dated to 892 Ma ± 13 Myr[48]. This provides for a minimum age constraint of 879 Ma.
**Soft Maximum age justification:** The Gunflint Chert microflora has a long history of study, including claims of eukaryotes. These include process-bearing acritarch-like cysts, such as *Germinosphaera*, suggesting the presence of an actin cytoskeleton. Nevertheless, all such claims of eukaryote affinity have generally been rejected[44]. The Gunflint Chert has been dated to 1878.3 Ma ± 1.3 Myr[45], yielding a maximum age of 1879.6 Ma.

**Dikarya - Mucoromycota | *Lobosporangium - Saccharomyces* | 452.71 Ma**
**Fossil taxon and specimen:** *Palaeoglomus grayi* UCMP 151983 (University of California Museum of Paleontology collection, Berkeley, California, USA) from Ordovician Guttenberg Formation, Galena Group, Decorah subgroup at Country Highway P site, Dane County, Wisconsin, USA[49,50].
**Phylogenetic justification:** Allied to *Glomus*-type Glomales on the basis of the general gestalt of the spores and their retention of a single hypha which distinguishes it from Acaulosporaceae and Gigasporaceae[50].
**Age justification:** The Guttenberg Formation is characterized by the Guttenberg Isotope Carbon Excursion (GICE) that is dated to 452.71 Ma[51].

**Crown Metazoa | *Amphimedon-Homo* | 574-609 Ma**
**Fossil taxon and specimen**: *Charnia masoni* (OUM ÁT.429/p) from level DRK-10 within the Drook Formation *sensu* Matthews et al. (2020) at Mistaken Point, Newfoundland[52].

**Phylogenetic justification**: *Charnia masoni* has been interpreted as a stem eumetazoan by[53]. **Minimum age justification:** DRK-10 within the Drook Formation at Mistaken Point has been directly dated to 574.17 Ma ± 2.8 Myr through Zircon U/Pb dating 573.4 Ma ± 0.28 Myr using Pb/Pb dating[54]. However, Matthews et al. (2020) demonstrate and Yang et al. (2021)[55] argue that rangeomorphs have been found in the rock record from 574 Ma.

**Soft maximum age justification:** Lantian biota - this biota has extensive macrofossils but nothing that can definitively be classified as metazoans. Yang et al. (2021)[55] establish a maximum age of 609 Ma for the Lantian Biota. This age allows for the possibility of *Eoandromeda* being a crown metazoan.

**Crown Eumetazoa | *Nematostella-Homo* | 561.1-590.8 Ma**

**Fossil taxon and specimen**: *Auroralumina attenboroughii* (GSM 106119; British Geological Survey, Nottingham, UK) from Bed B, Bradgate Formation, Maplewell Group, Charnian Supergroup, Leicestershire, UK[56].

**Phylogenetic Justification:** *Auroralumina attenboroughii* has been shown to be a crown-group member based on tetraradial periderm with corner sulci, allying it with medusozoans[56].

**Minimum age justification:** *Auroralumina attenboroughii* was recovered from Bed B, Bradgate Formation, Charnian Supergroup, which has a minimum age of 563 ± 1.9 Ma, or 561.1 Ma[57]. Newer U-Pb data suggest the minimum age of the Bradgate Formation is 556.6 ± 6.4, however this large age uncertainty entirely encompasses the maximum age for the clade. For this reason, the age of 561.1 Ma, based on Wilby et al. (2011), will be used.

**Soft maximum age justification:** Established based on the Weng'an biota which may contain total group metazoans[58], but there is no convincing evidence of crown metazoans. Yang et al. (2021)[55] establish a 590.8 Ma maximum age for the Weng'an Biota.

**Crown Protostomia | *Lottia-Limulus* | 532-590.8 Ma**

**Fossil taxon and specimen:** *Aldanella yanjiahensis*, from the Dahai member of the Zhujiaqing Formation in the middle Meishucunian of China, TU Berlin collection NO. YXII02-02[59].

**Phylogenetic justification:** *Aldanella janjiahensis* (junior synonym *Aldanella attleborensis*) is a dextrally-coiled stem group gastropod assigned to Pelagiellida[60].

**Minimum age justification:** *Aldanella yanjiahensis* is associated with *Watsonella crosbyi* and *Oelandiella korobkovi* in the Dahai member in the middle Meishucunian of China[59]. Chemostratigraphic correlation places this unit in the Nemakit Daldynian within the interval 534-532 Ma[61].

**Soft maximum justification:** Established based on the Weng'an biota which may contain total group metazoans[58], but there is no convincing evidence of crown metazoans. Yang et al. (2021)[55] establish a 590.8 Ma maximum age for the Weng'an Biota.

**Mesangiospermae | *Arabidopsis-Oryza* | 121.4-247.3 Ma**

**Fossil taxon and specimen:** Tricolpate pollen grain [palynological sample BRN 126], from the middle Atherfield Wealden Bed 35 of the Cowleaze Chine Member, Vectis Formation, Barremian (Early Cretaceous), of the Isle of Wight[62].

**Phylogenetic justification**: Following Clarke et al. 2011[63], our minimum age constraint is based on the earliest occurrences Fischer's rule tricolpate pollen, and knowledge of the distribution of tricoplate pollen across the phylogeny of angiosperms[64].

**Minimum age justification:** Following Clarke at al. (2011)[63], the Cowleaze Chine Member of the Vectis Formation of the Isle of Wight[62] occurs within the M1n polarity chron, the top of which is dated to 121.4 Ma[65].

**Soft maximum age justification:** The soft maximum age constraint is based on sediments devoid of angiosperm-like pollen below their first report in the Middle Triassic. Type C 'Retisulcites' of Hochuli and Feist-Burkhardt were recovered from the Steinkobbe Formation, Norwegian Arctic. The Steinkobbe Formation is considered to be middle Anisian in age [66], thus the maximum age of these pollen grains is at the base of the Anisian, dated at 247.1 ± 0.2 Ma

**Constraint: Last plastid common ancestor younger than last eukaryotic common ancestor.**

**Phylogenetic justification:** The Archaeplastida are nested within crown eukaryotes, and the host cell for the cyanobacterial endosymbiont already possessed both a nucleus and mitochondria[67]. Therefore, the node corresponding to the last common ancestor of plastids (LPCA) must be younger than the nodes corresponding to the last common ancestors of eukaryotes within the archaeal and proteobacterial regions of the universal tree.

Please note that we were unable to apply the *Bangiomorpha* calibration within plastids in the ATP synthase analysis because the sequences from rhodophytes were not recovered as monophyletic (Supplementary Figures 12, 13, 21). As a result, clade ages inferred from the species tree analysis are likely to be more reliable since they make use of all the available evidence, while the main interest of inferred node ages from the ATP synthase tree are for interpreting the evolutionary history of that particular gene family.

### Supplementary Figures

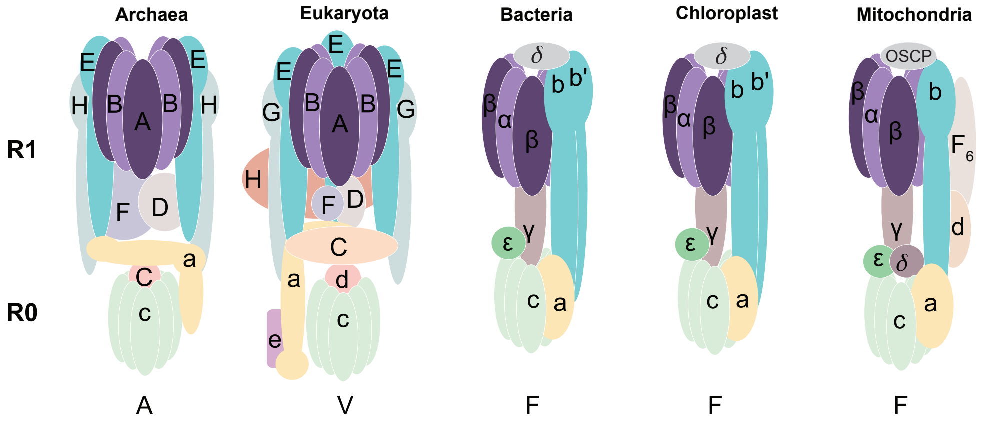

**Supplementary Figure 1**: **Structure of F-, A-, and V-type ATP synthases**. Schematic depicts the conserved R1 and R0 subcomplexes across the ATP synthase types (F-type in Bacteria, Chloroplast, and Mitochondria; the A-type in Archaea; the V-type in Eukaryota) inspired by Nirody et al., 2020[68]. Structural variation is observed in the central and peripheral stalk components. Color-coding corresponds to homologous subunits. The catalytic and non-catalytic subunits of the R1 complex are highlighted as follows: F1-alpha (*nc*F1) and A1/V1 B (*nc*A1V1) (light purple), F1-beta (*c*F1) and A1/V1 A (*c*A1V1) (dark purple).

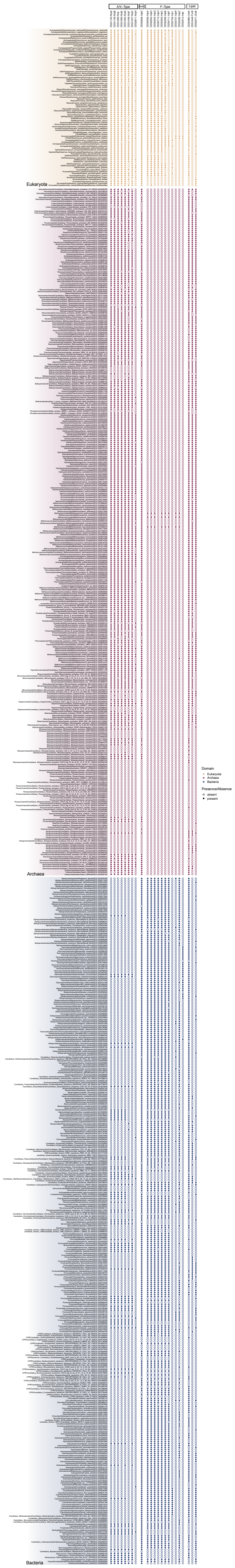
A/V−Type
F−Type
Lipid
Eukaryota
Archaea
Bacteria
Domain
Eukaryota
Archaea
Bacteria
Presence/Absence
absent
present

**Supplementary Figure 2**: **Presence/absence distribution of F- and A/V-type ATP synthase subunits and selected lipid biosynthesis genes across the 780 archaea, bacteria, and eukaryotes present in the concatenated species phylogeny**. COGs corresponding to the A/V- and F-type ATP synthases are as listed in the NCBI COG Pathways May 2020[69–71]. COG0636 represents the proteolipid subunit that is homologous to both A/V- and F-type ATP synthases. Lipid biosynthesis genes correspond to 4-hydroxybenzoate polyprenyltransferase (COG0382), Glycerol-1-phosphate heptaprenyltransferase (COG1646), and Glycerol dehydrogenase or related enzyme (COG0371). A table listing the presence/absence counts for all taxa can be found in Supplementary Table 4 and a list of the COGs shown here can be found in Supplementary Table 3.

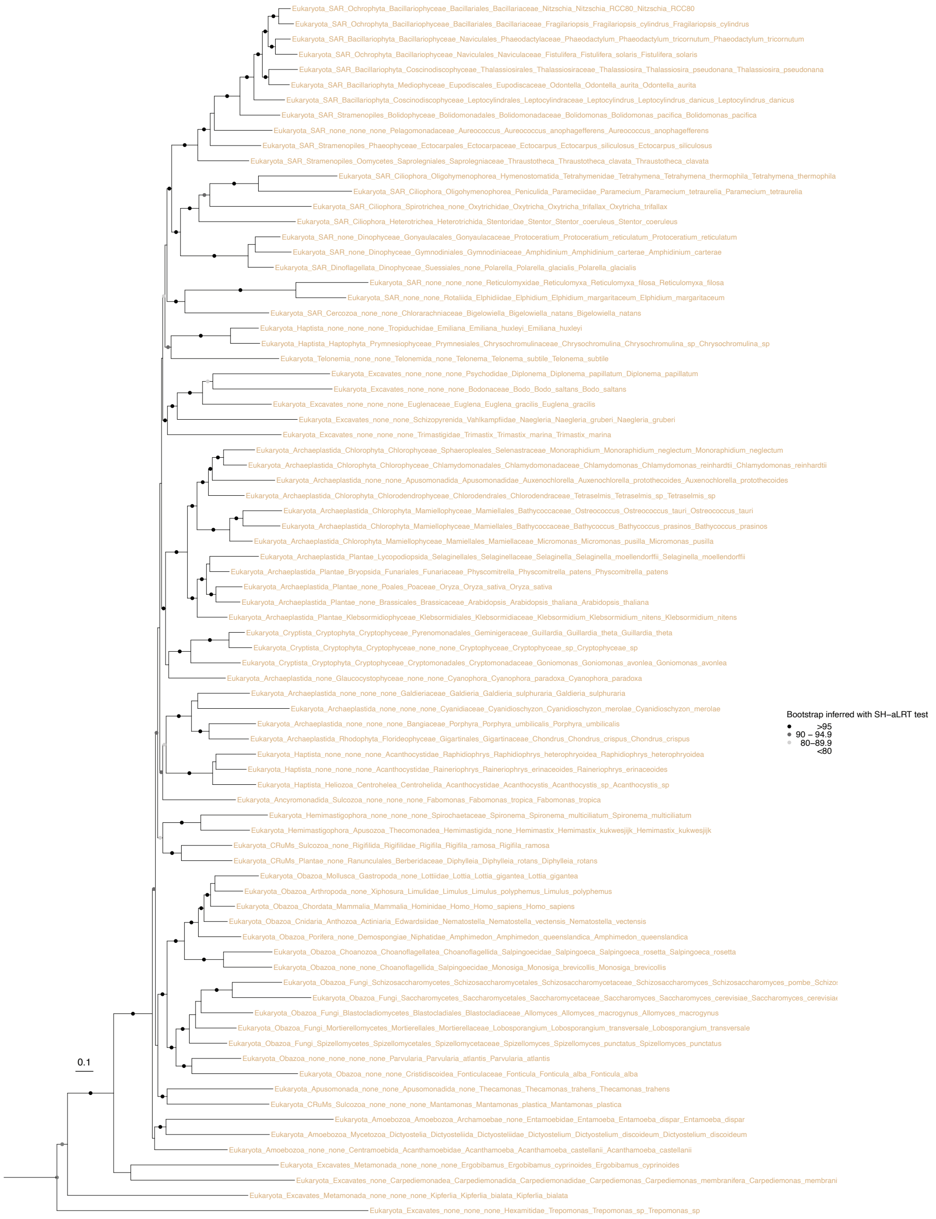

**Supplementary Figure 3**: **Phylogenetic inference of 80 eukaryotes based on 21 conserved universal marker genes (Supplementary Table 14)**. Single-copy marker gene sequences were extracted from the 100 eukaryotes included in the reference species set. Eukaryotes that contained less than 65% of all marker genes (20 species, see SOMX) were excluded from the phylogenetic analysis. Sequences were aligned using MAFFT L-INS-i and trimmed using BMGE (alignment length = 5222 bp). The maximum likelihood phylogeny was inferred IQ-TREE2 v2.1.2 with the LG+C60+R+F model and ultrafast bootstrap approximation (left) and SH-like approximate likelihood (right) tests, both run with 1000 replicates. Scale bar: average number of substitutions per site. The tree is midpoint-rooted.

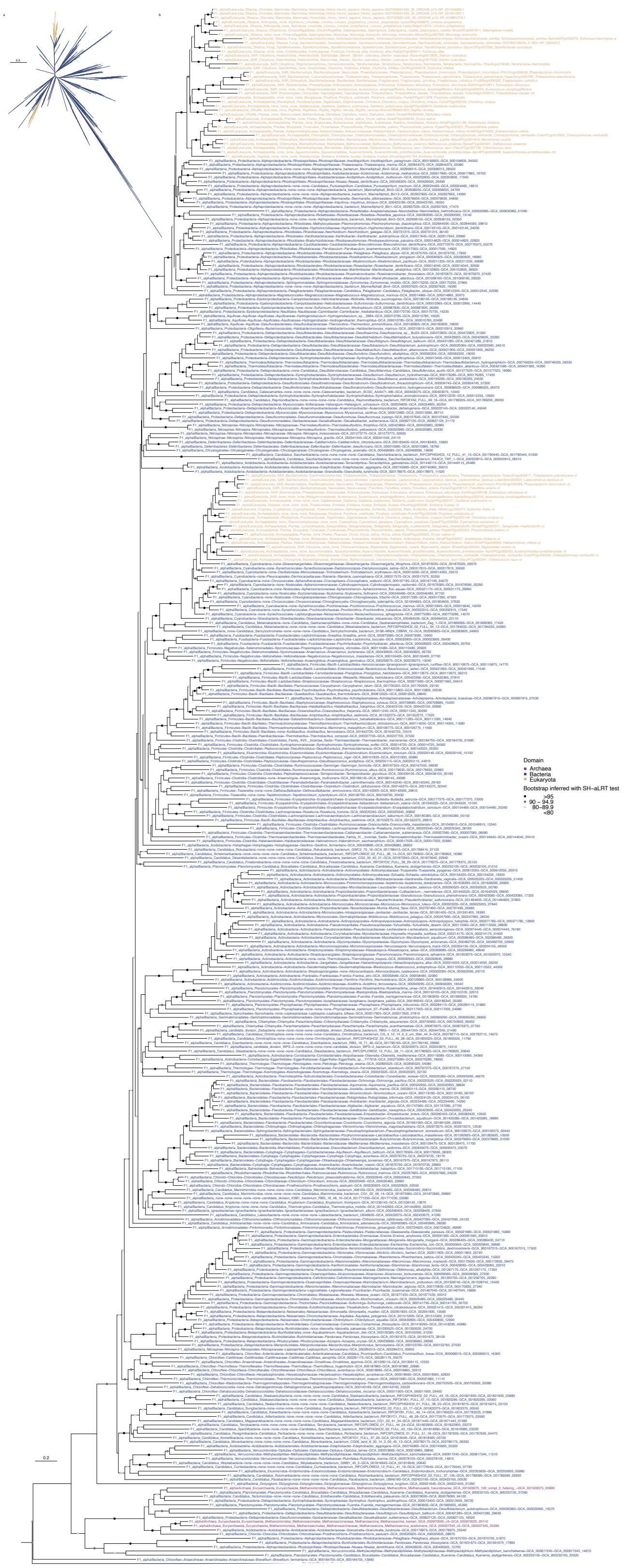
a
b
0.2
Domain
Archaea
Bacteria
Eukaryota
Bootstrap inferred with SH−aLRT test
>95
90 – 94.9
80–89.9
<80
0.2

**Supplementary Figure 4**: **Phylogenetic history of the F1-alpha (*nc*F1) ATP synthase subunit in 800 examined Archaea (red), Bacteria (blue), and eukaryotes (yellow)**. The alignment contains 371 sequences and was trimmed with BMGE (alignment length = 453 bp). The maximum likelihood tree was inferred in IQ-TREE2 v2.1.2 with the LG+C60+R+F model with ultrafast bootstrap approximation (left) and SH-like approximate likelihood (right), each with 1000 replicates. Scale bar: average number of substitutions per site. (a) unrooted, (b) midpoint-rooted.

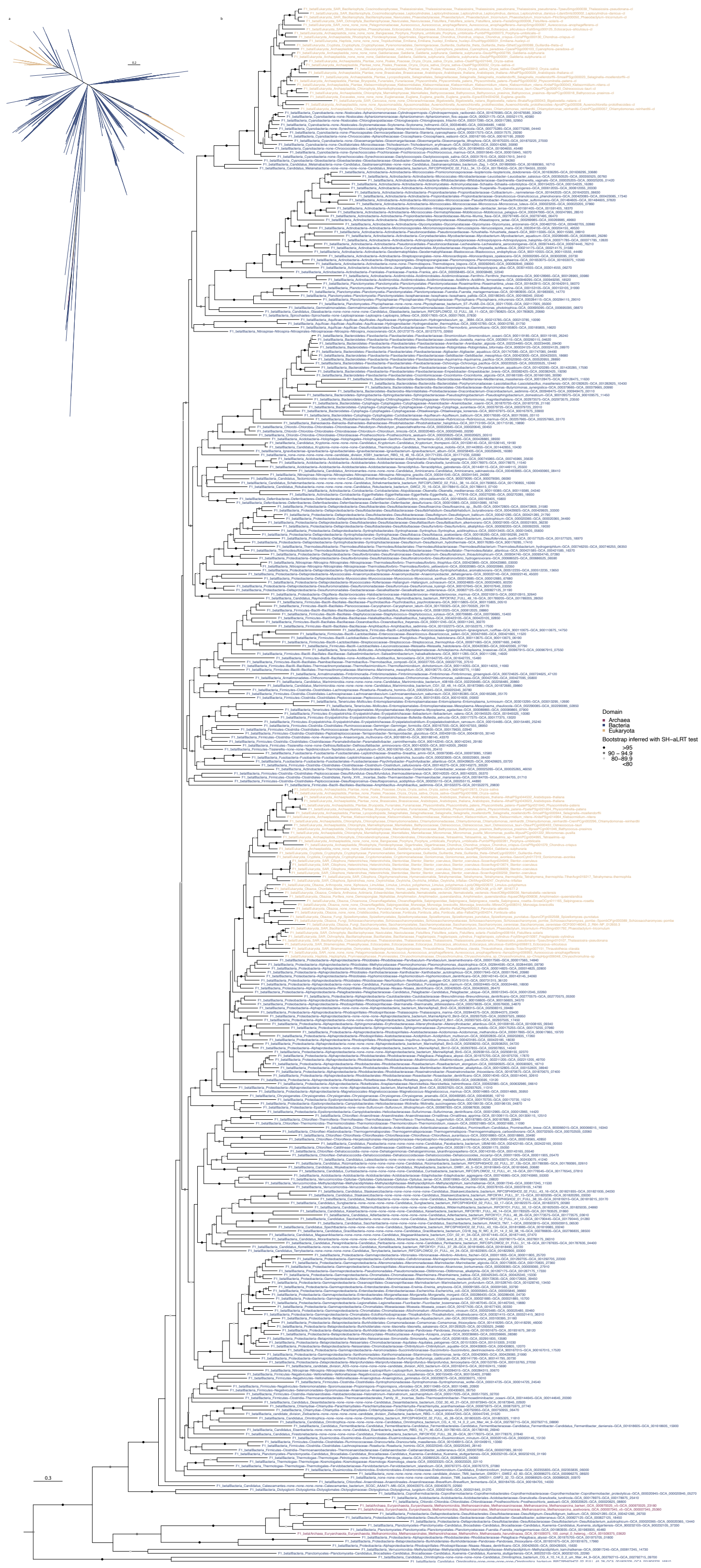
a
b
Domain
Archaea
Bacteria
Eukaryota
Bootstrap inferred with SH–aLRT test
>95
90 – 94.9
80–89.9
<80
0.3

**Supplementary Figure 5**: **Phylogenetic history of the F1-beta (*c*F1) ATP synthase subunit in the 800 examined Archaea (red), Bacteria (blue), and eukaryotes (yellow)**. The alignment contains 391 sequences and was trimmed with BMGE (alignment length = 434 bp). The maximum likelihood tree was inferred in IQ-TREE2 v2.1.2 with the LG+C60 model with ultrafast bootstrap approximation (left) and SH-like approximate likelihood (right), each with 1000 replicates. Scale bar: average number of substitutions per site. (a) unrooted, (b) midpoint-rooted.

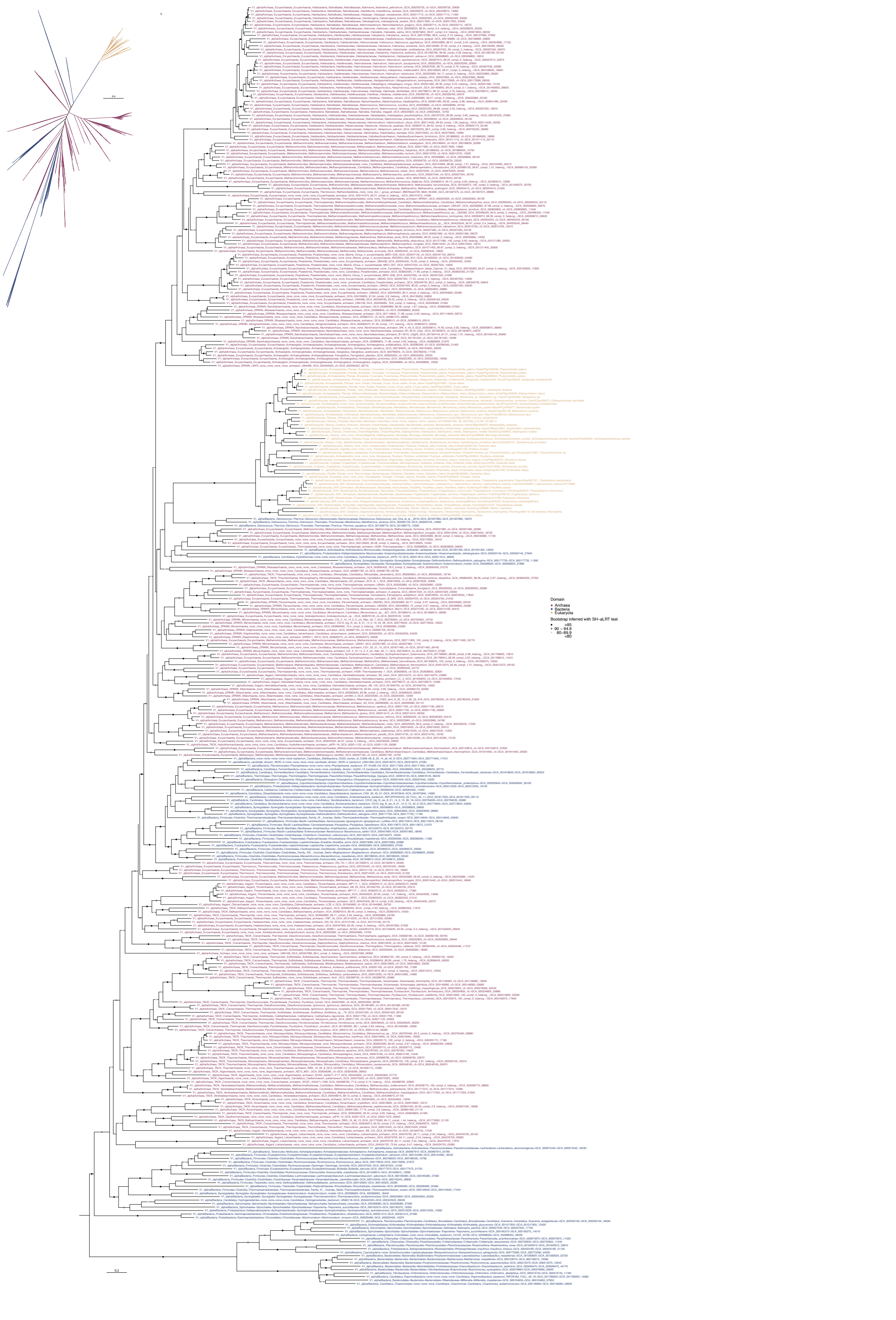
Domain
Archaea
Bacteria
Eukaryota
Bootstrap inferred with SH–aLRT test
>95
90 – 94.9
80–89.9
<80

**Supplementary Figure 6**: **Phylogenetic history of the A1/V1 A (*c*A1V1) ATP synthase subunit in the 800 examined Archaea (red), Bacteria (blue), and eukaryotes (yellow)**. The alignment contains 368 sequences and was trimmed with BMGE (alignment length = 512 bp). The maximum likelihood tree was inferred in IQ-TREE2 v2.1.2 with the LG+C50+R+F model with ultrafast bootstrap approximation (left) and SH-like approximate likelihood (right), each with 1000 replicates. Scale bar: average number of substitutions per site. (a) unrooted, (b) midpoint-rooted.

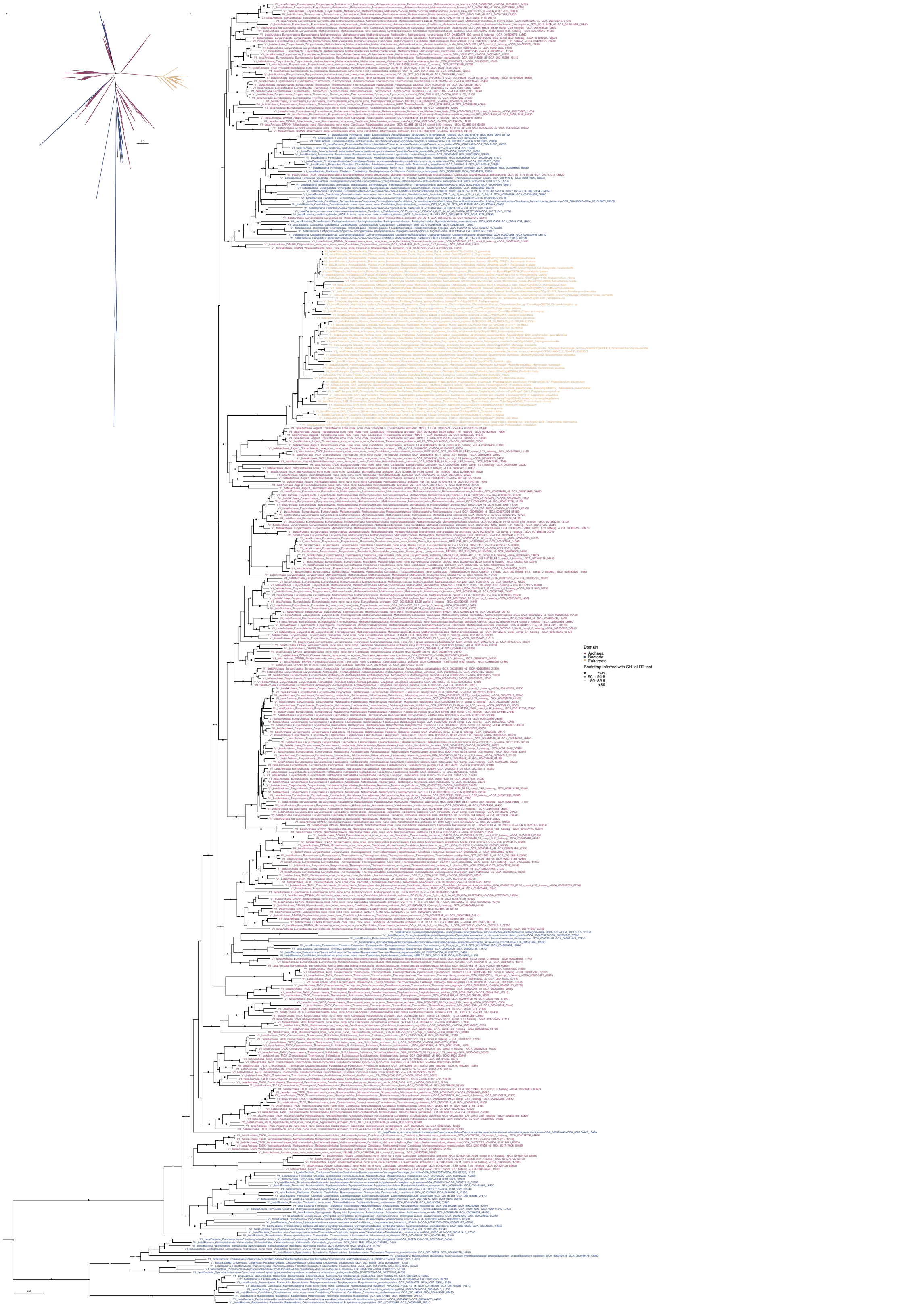
Domain
Archaea
Bacteria
Eukaryota
Bootstrap inferred with SH−aLRT test
>95
90 – 94.9
80–89.9
<80

**Supplementary Figure 7**: **Phylogenetic history of the A1/V1 B (*nc*A1V1) ATP synthase subunit in the 800 examined Archaea (red), Bacteria (blue), and eukaryotes (yellow)**. The alignment contains 390 sequences and was trimmed with BMGE (alignment length = 433 bp). The maximum likelihood tree was inferred with the LG+C60+R+F model with ultrafast bootstrap approximation (left) and SH-like approximate likelihood (right), each with 1000 replicates. Scale bar: average number of substitutions per site. (a) unrooted, (b) midpoint-rooted.

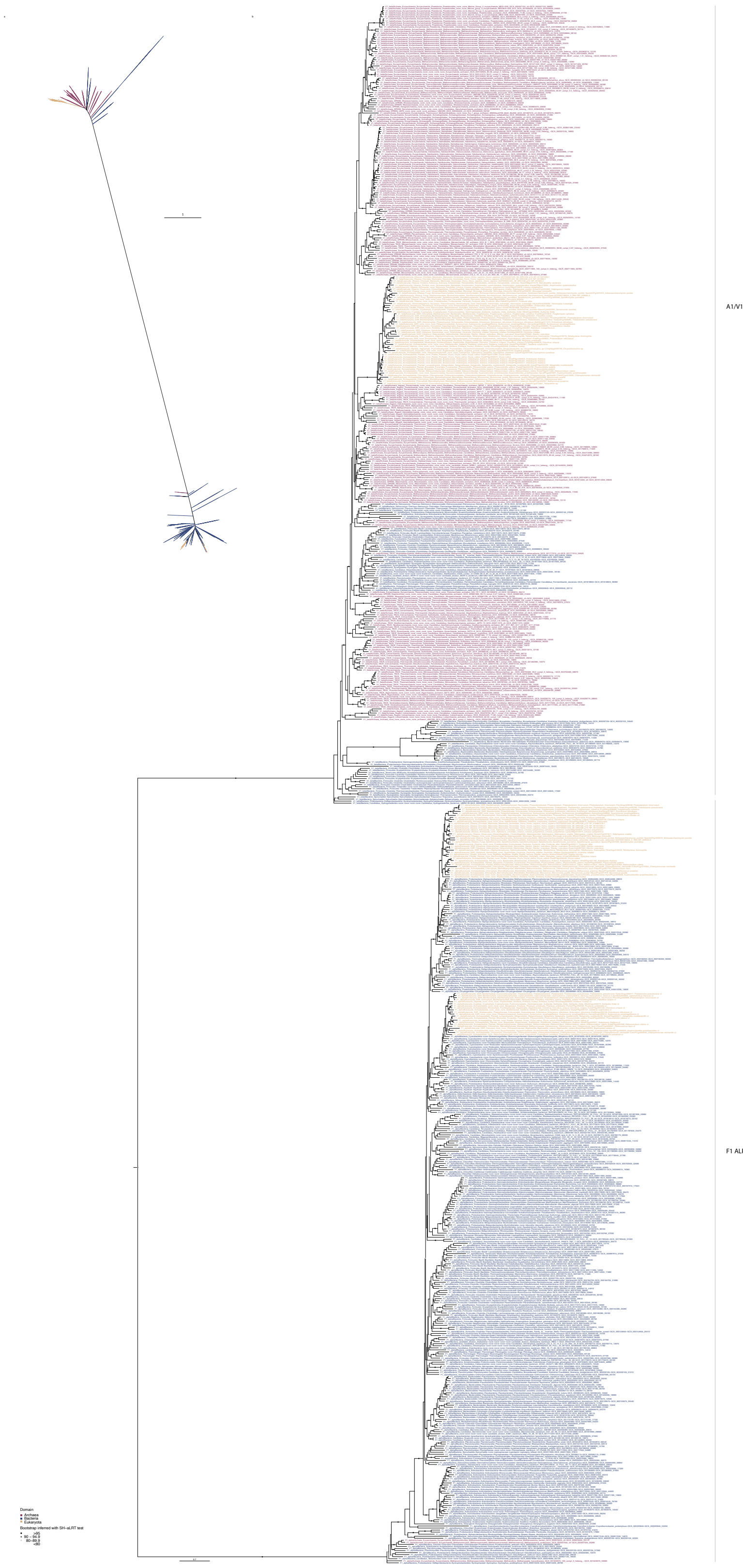

A1/V1 BETA
F1 ALPHA
Domain
Archaea
Bacteria
Eukaryota
Bootstrap inferred with SH–aLRT test
>95
90 – 94.9
80–89.9
<80

**Supplementary Figure 8**: **Phylogenetic history of the non-catalytic (*nc*) subunits of the F- and A/V-type ATP synthase (F1-alpha and A1V1 B) in the 800 examined Archaea (red), Bacteria (blue), and eukaryotes (yellow)**. The alignment contains 761 sequences and was trimmed with BMGE (alignment length = 389 bp). The maximum likelihood tree was inferred in IQ-TREE2 v2.1.2 with the LG+C60+R+F model with ultrafast bootstrap approximation (left) and SH-like approximate likelihood (right), each with 1000 replicates. Scale bar: average number of substitutions per site. (a) unrooted, (b) midpoint-rooted.

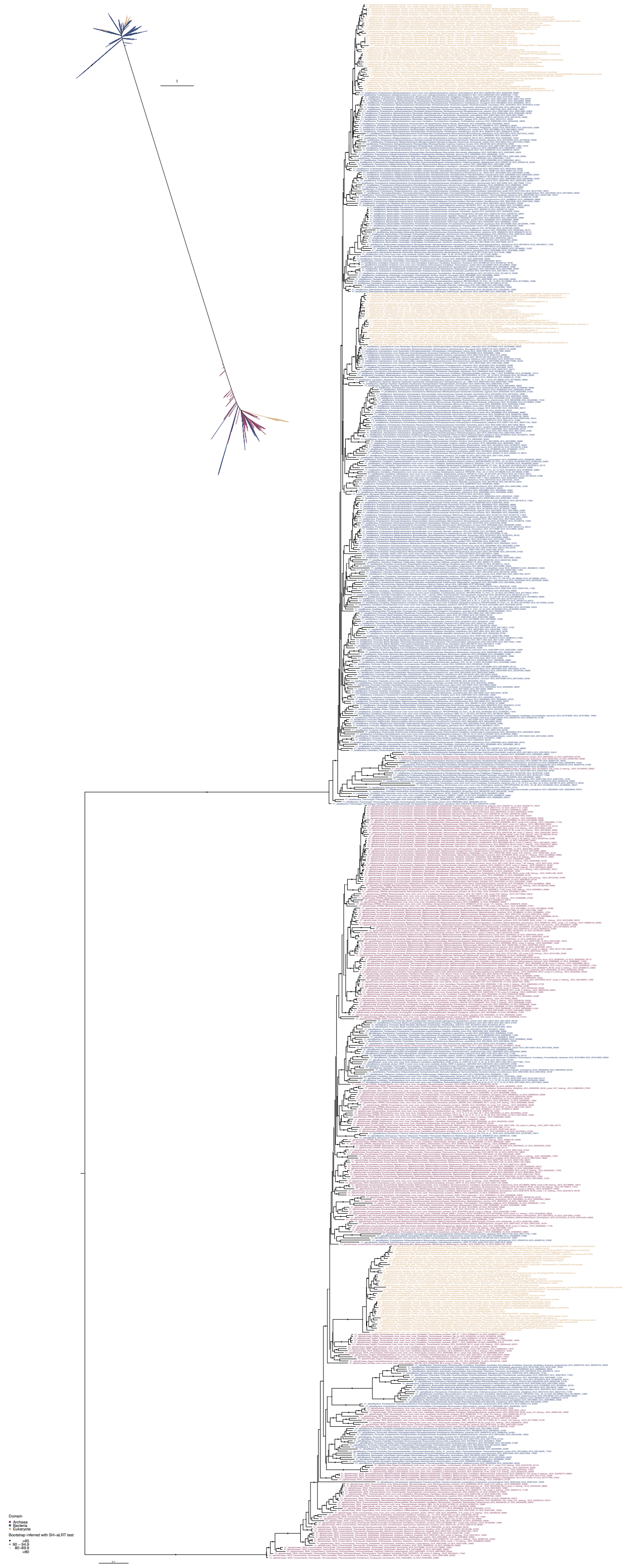

**Supplementary Figure 9**: **Phylogenetic history of the catalytic (*c*) subunits of the F- and A/V-type ATP synthase (F1-beta and A1V1 A) in the 800 examined Archaea (red), Bacteria (blue), and eukaryotes (yellow)**. The alignment contains 759 sequences and was trimmed with BMGE (alignment length = 390 bp). The maximum likelihood tree was inferred in IQ-TREE2 v2.1.2 with the LG+C60+R+F model with ultrafast bootstrap approximation (left) and SH-like approximate likelihood (right), each with 1000 replicates. Scale bar: average number of substitutions per site. (a) unrooted, (b) midpoint-rooted.

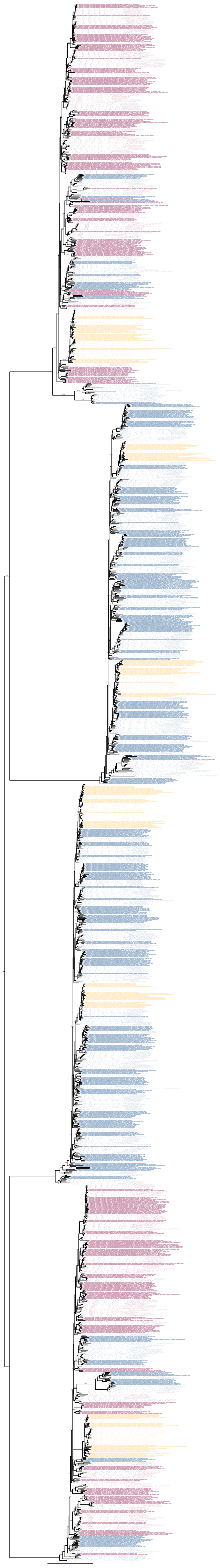

**Supplementary Figure 10**: **Phylogenetic history of the all headpiece subunits of the F- and A/V-type ATP synthase (F1-alpha/*nc*F1, F1-beta/*c*F1, A1/V1 A/*c*A1V1, A1/V1 B/*nc*A1V1) in the 800 examined Archaea (red), Bacteria (blue), and eukaryotes (yellow)**. The alignment contains 1520 sequences and was trimmed with BMGE (alignment length = 350 bp). The maximum likelihood tree was inferred in IQ-TREE2 v2.1.2 with the LG+C50+R+F model with ultrafast bootstrap approximation (left) and SH-like approximate likelihood (right), each with 1000 replicates. Scale bar: average number of substitutions per site. (a) unrooted, (b) midpoint-rooted.

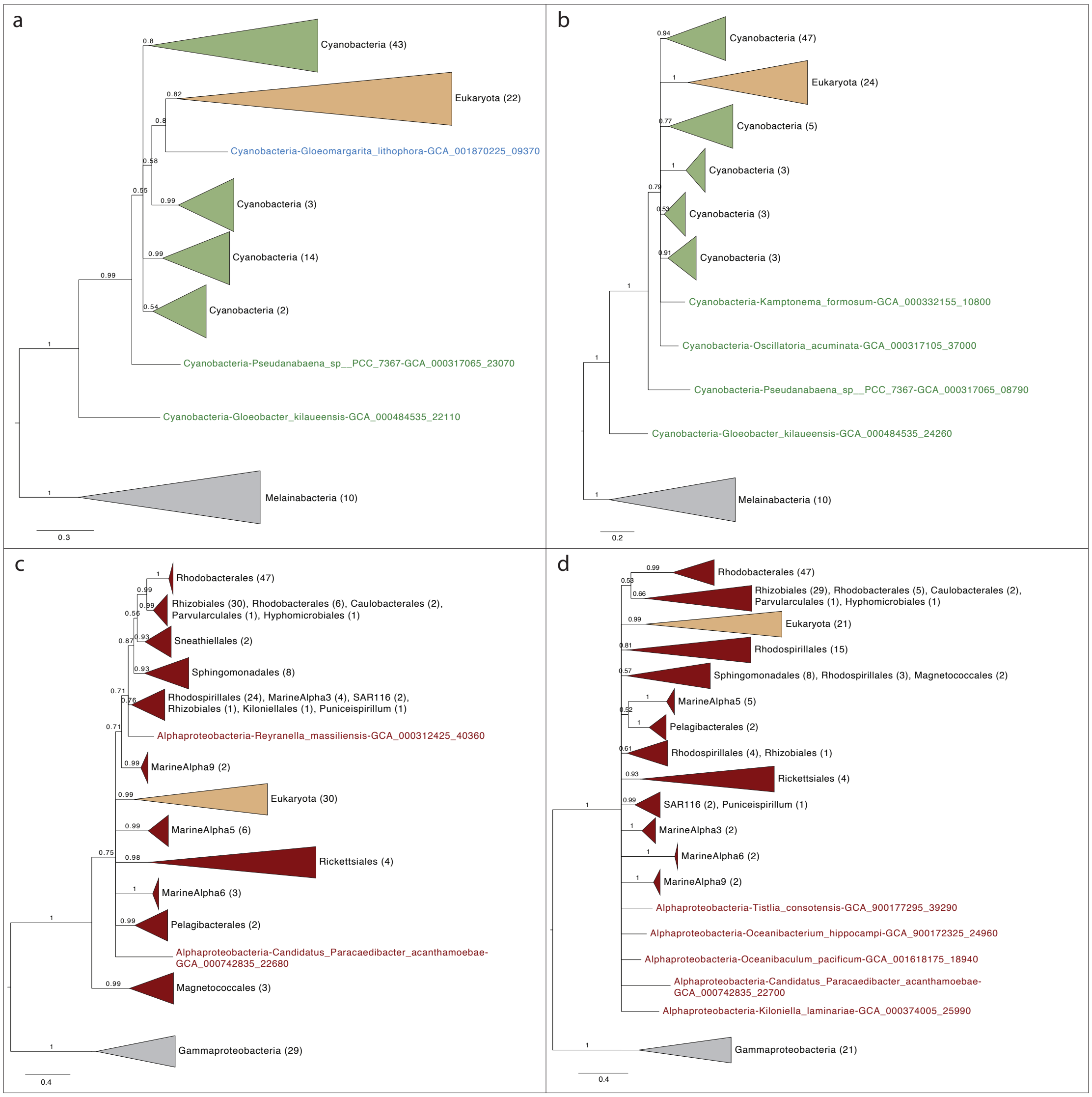

a
Cyanobacteria (43)
Eukaryota (22)
Cyanobacteria-Gloeomargarita_lithophora-GCA_001870225_09370
Cyanobacteria (3)
Cyanobacteria (14)
Cyanobacteria (2)
Cyanobacteria-Pseudanabaena_sp__PCC_7367-GCA_000317065_23070
Cyanobacteria-Gloeobacter_kilaueensis-GCA_000484535_22110
Melainabacteria (10)
0.3
b
Cyanobacteria (47)
Eukaryota (24)
Cyanobacteria (5)
Cyanobacteria (3)
Cyanobacteria (3)
Cyanobacteria (3)
Cyanobacteria-Kamptonema_formosum-GCA_000332155_10800
Cyanobacteria-Oscillatoria_acuminata-GCA_000317105_37000
Cyanobacteria-Pseudanabaena_sp__PCC_7367-GCA_000317065_08790
Cyanobacteria-Gloeobacter_kilaueensis-GCA_000484535_24260
Melainabacteria (10)
0.2
c
Rhodobacterales (47)
Rhizobiales (30), Rhodobacterales (6), Caulobacterales (2),
Parvularculales (1), Hyphomicrobiales (1)
Sneathiellales (2)
Sphingomonadales (8)
Rhodospirillales (24), MarineAlpha3 (4), SAR116 (2),
Rhizobiales (1), Kiloniellales (1), Puniceispirillum (1)
Alphaproteobacteria-Reyranella_massiliensis-GCA_000312425_40360
MarineAlpha9 (2)
Eukaryota (30)
MarineAlpha5 (6)
Rickettsiales (4)
MarineAlpha6 (3)
Pelagibacterales (2)
Alphaproteobacteria-Candidatus_Paracaedibacter_acanthamoebae-
GCA_000742835_22680
Magnetococcales (3)
Gammaproteobacteria (29)
0.4
d
Rhodobacterales (47)
Rhizobiales (29), Rhodobacterales (5), Caulobacterales (2),
Parvularculales (1), Hyphomicrobiales (1)
Eukaryota (21)
Rhodospirillales (15)
Sphingomonadales (8), Rhodospirillales (3), Magnetococcales (2)
MarineAlpha5 (5)
Pelagibacterales (2)
Rhodospirillales (4), Rhizobiales (1)
Rickettsiales (4)
SAR116 (2), Puniceispirillum (1)
MarineAlpha3 (2)
MarineAlpha6 (2)
MarineAlpha9 (2)
Alphaproteobacteria-Tistlia_consotensis-GCA_900177295_39290
Alphaproteobacteria-Oceanibacterium_hippocampi-GCA_900172325_24960
Alphaproteobacteria-Oceanibaculum_pacificum-GCA_001618175_18940
Alphaproteobacteria-Candidatus_Paracaedibacter_acanthamoebae-
GCA_000742835_22700
Alphaproteobacteria-Kiloniella_laminariae-GCA_000374005_25990
Gammaproteobacteria (21)
0.4

**Supplementary Figure 11**: **Bayesian phylogenetic placement of eukaryotic F-type ATP synthase catalytic and non-catalytic subunits assessing plastid and mitochondrial origins**. For computational tractability, we downsampled the taxa subsets containing eukaryotes, alphaproteobacteria, and gammaproteobacteria to a maximum of 250 taxa (*nc*F1A: 211, *c*F1B:185 sequences). Sequences were cleaned, filtered, de-replicated, aligned, and trimmed using the same conditions described in the main text. Bayesian phylogenies were constructed using PhyloBayes-MPI (version 1.5) using the CAT-GTR model with four discrete gamma categories for rates across sites; for each alignment, four independent Markov Chain Monte Carlo (MCMC) chains were run. Each chain was run over 100,000 iterations (or until convergence). Convergence was evaluated using the bpcomp and tracecomp tools within PhyloBayes-MPI, with 1000 generations discarded as burn-in and sub-sampling every 10 trees. The final consensus trees were generated through bpcomp using the same settings. a. Bayesian phylogeny of F1-alpha (*nc*F1) plastid eukaryotic homologs, Cyanobacteria, and Melainabacteria. The alignment has 97 sequences, was trimmed with BMGE (alignment length=505 bp), and a Bayesian phylogeny was inferred with (CAT + GTR) model (see above). b. Bayesian phylogeny of F1-beta (*c*F1) plastid eukaryotic homologs, Cyanobacteria, and Melainabacteria. The alignment has 98 sequences, was trimmed with BMGE (alignment length=481 bp), and a Bayesian phylogeny was inferred with (CAT+GTR) model (see above). a and b are manually rooted with Melainabacteria as an outgroup. c. Bayesian phylogeny of F1-alpha (*nc*F1) mitochondrial eukaryotic homologs, Alphaproteobacteria, and Gammaproteobacteria. The alignment has 211 sequences, was trimmed with BMGE (alignment length=457 bp), and a Bayesian phylogeny inferred with (CAT+GTR) model (see above). d. Bayesian phylogeny of F1-beta (*c*F1) mitochondrial eukaryotic homologs, Alphaproteobacteria, and Gammaproteobacteria. The alignment has 185 sequences, was trimmed with BMGE (alignment length=497 bp), and a Bayesian phylogeny was inferred with (CAT+GTR) model (see above). c and d are manually rooted with Gammaproteobacteria as an outgroup.

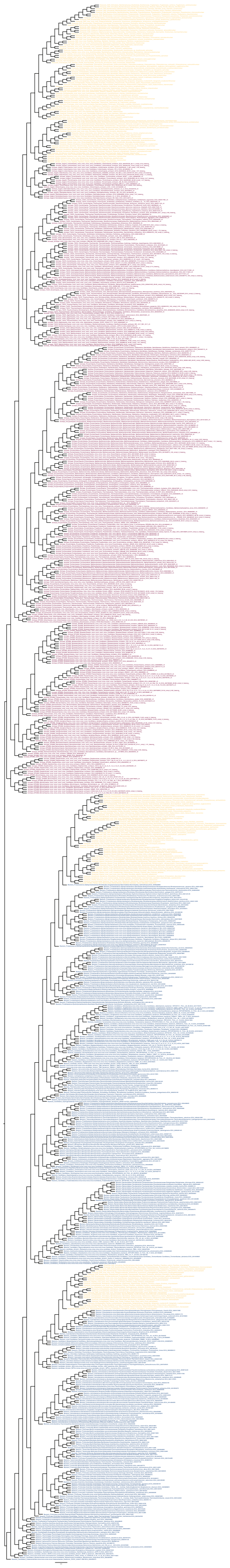

**Supplementary Figure 12**: **Edited1 species phylogeny used for molecular dating (see Methods, Edited1, Supplementary Figure 21)**. Select eukaryotic speciations were constrained in order to apply necessary fossil calibrations to the tree (see Methods). The Edited1 constrained topology was inferred in IQ-TREE2. The phylogeny includes 863 taxa of examined Archaea (red), Bacteria (blue), and nuclear, mitochondrial, and plastid eukaryotic homologs (yellow).

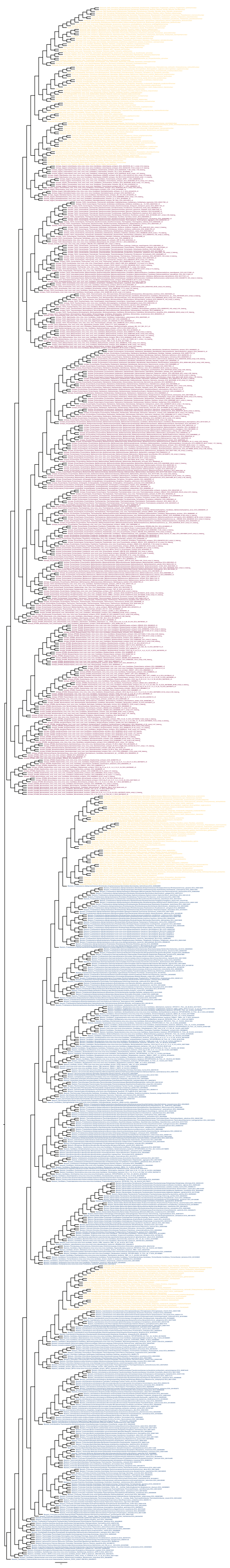

**Supplementary Figure 13**: **Edited2 species phylogeny used for molecular dating (see Methods, Edited2, Supplementary Figure 21)**. In addition to the eukaryotic constraints applied to the Edited1 topology (Supplementary Figure 12) we positioned the nuclear homologs sister to the Hodarchaea (see Methods) and the mitochondrial homologs sister to all alphaproteobacteria excluding the *Magnetococcales* (see Methods). The Edited2 constrained topology was inferred in IQ-TREE2. The phylogeny includes 863 taxa of examined Archaea (red), Bacteria (blue), and nuclear, mitochondrial, and plastid eukaryotic homologs (yellow).

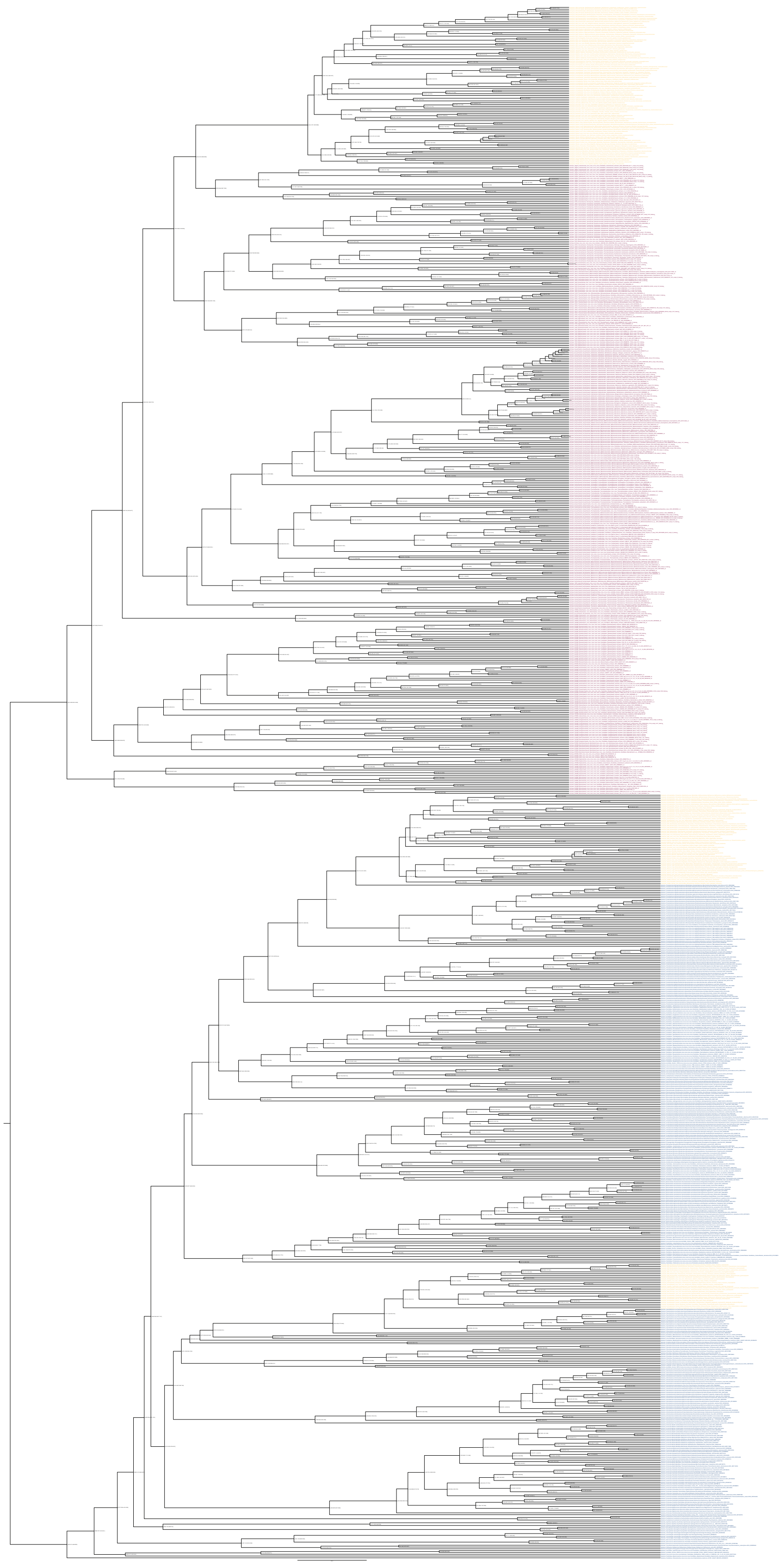

**Supplementary Figure 14: Dated cross-braced ribosomal species tree Edited1 (long) including nuclear, mitochondrial, and plastid eukaryotic homologs (yellow), Archaea (red), and Bacteria (blue)**. Age ranges are indicated for each node, represented as 95% Highest Posterior densities (HPDs).

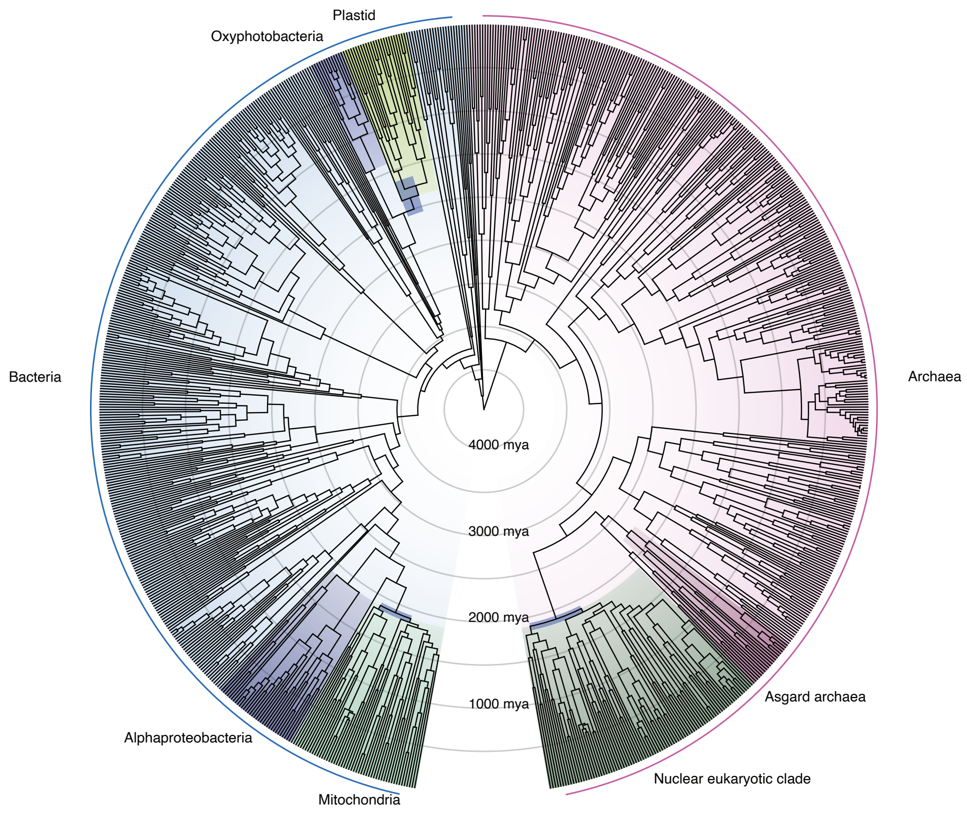

**Supplementary Figure 15**: **Dated cross-braced ribosomal species tree Edited1 (circularized) including nuclear (green), mitochondrial (green), and plastid (yellow) eukaryotic homologs, Archaea (red), and Bacteria (blue)**. Alphaproteobacteria and Oxyphotobacteria are highlighted in dark blue, while Asgard archaea are highlighted in dark red. Geologic time scale is represented as concentric circles every 500 million years.

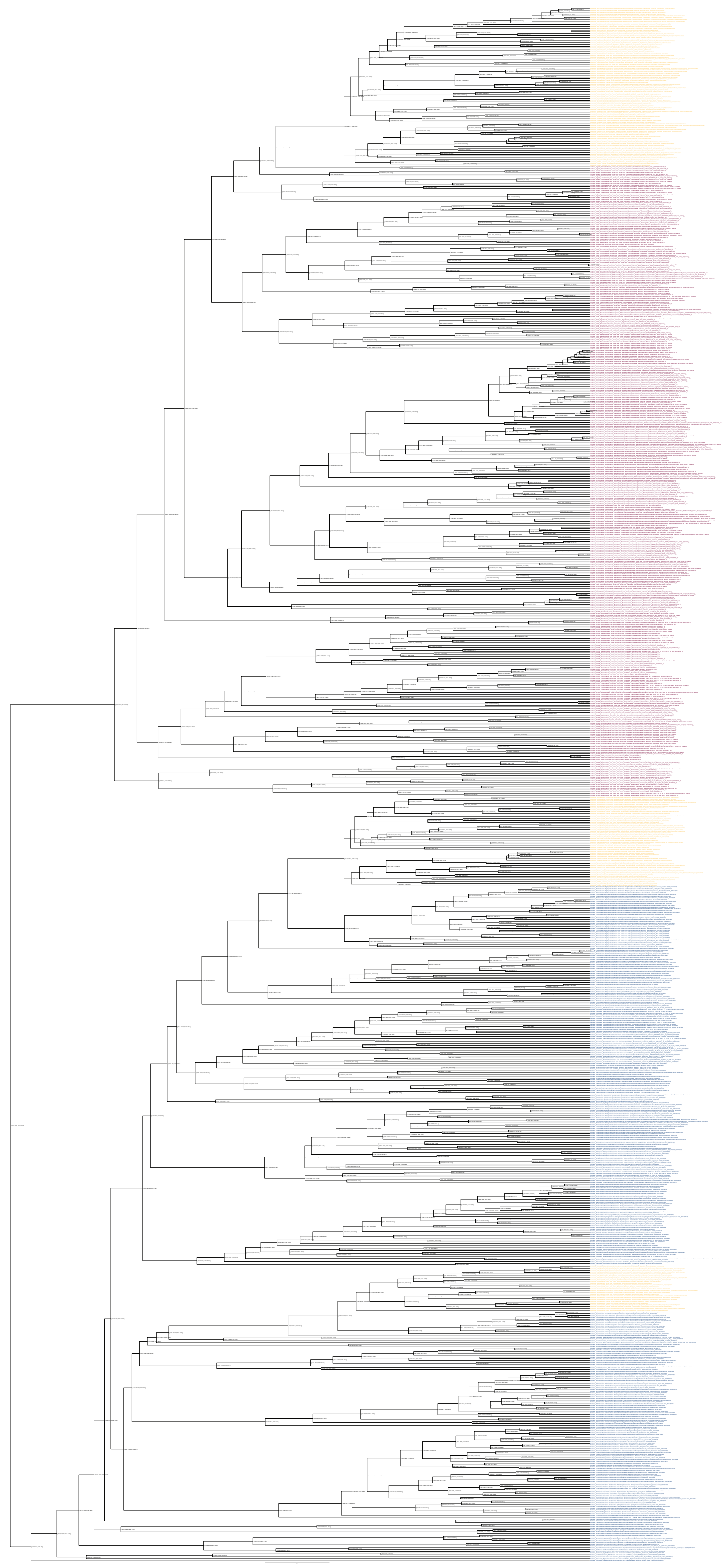

548

**Supplementary Figure 16**: **Dated cross-braced ribosomal species tree Edited2 (long) including nuclear, mitochondrial, and plastid eukaryotic homologs (yellow), Archaea (red), and Bacteria (blue)**. Age ranges are indicated for each node, represented as 95% Highest Posterior densities (HPDs).

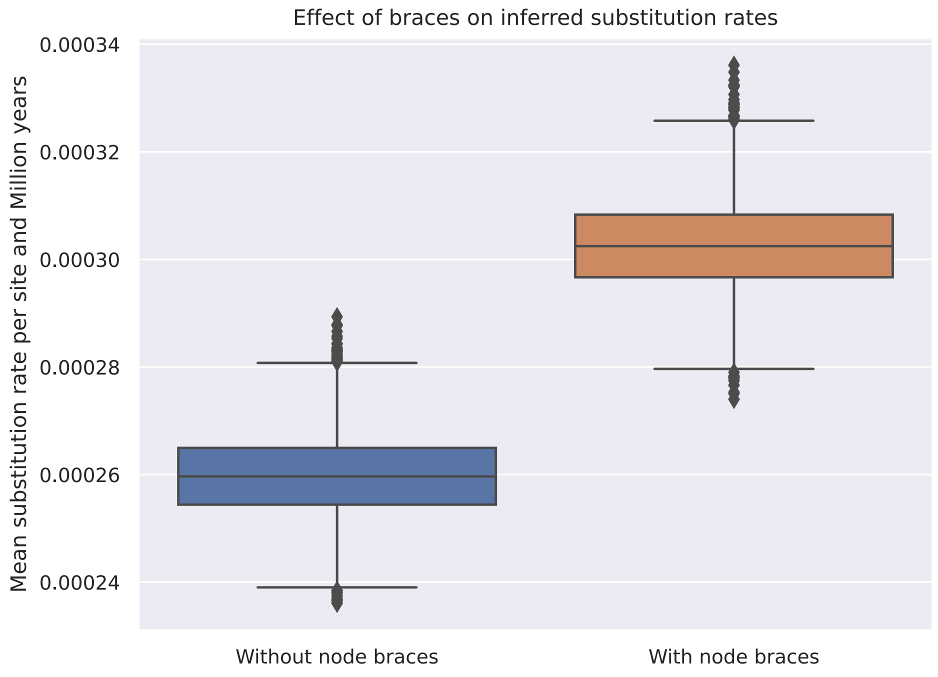

**Supplementary Figure 17**: **The effect of braces on inferred substitution rates**. By propagating eukaryotic fossil information to otherwise poorly-constrained regions of the bacterial phylogeny, bracing equivalent speciation nodes substantially increases overall rates of molecular evolution (measured as the expected number of substitutions per site per Million years).

559

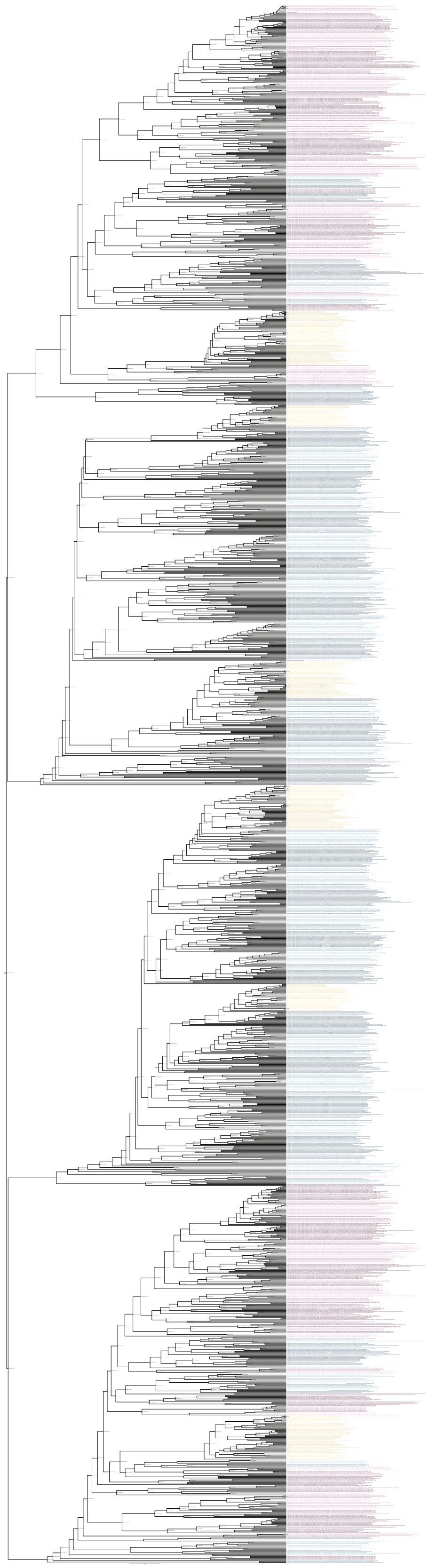

**Supplementary Figure 18**: **Dated cross-braced combined ATP synthase gene tree (long) including Archaea (red), Bacteria (blue), and Eukaryotes (yellow)**. Annotations for F- and A/V-type catalytic and non-catalytic subunits are assigned to each tip name. Age ranges are indicated for each node, represented as 95% Highest Posterior densities (HPDs).

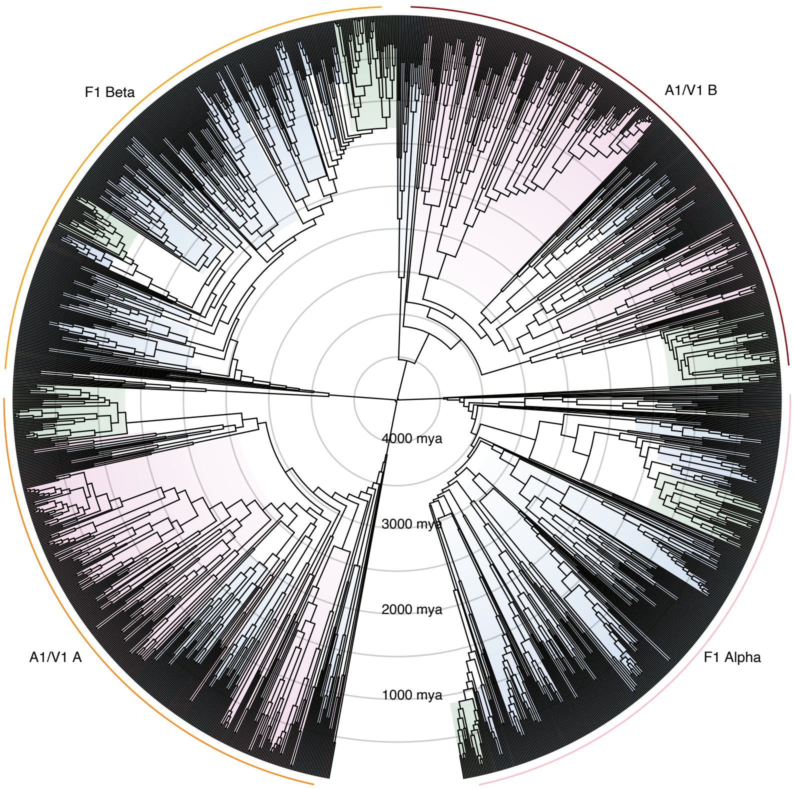

**Supplementary Figure 19**: **Dated cross-braced combined ATP synthase gene tree (circularized) including Archaea (red), Bacteria (blue), and Eukaryotes (green)**. Geologic time scale is represented as concentric circles every 500 million years.

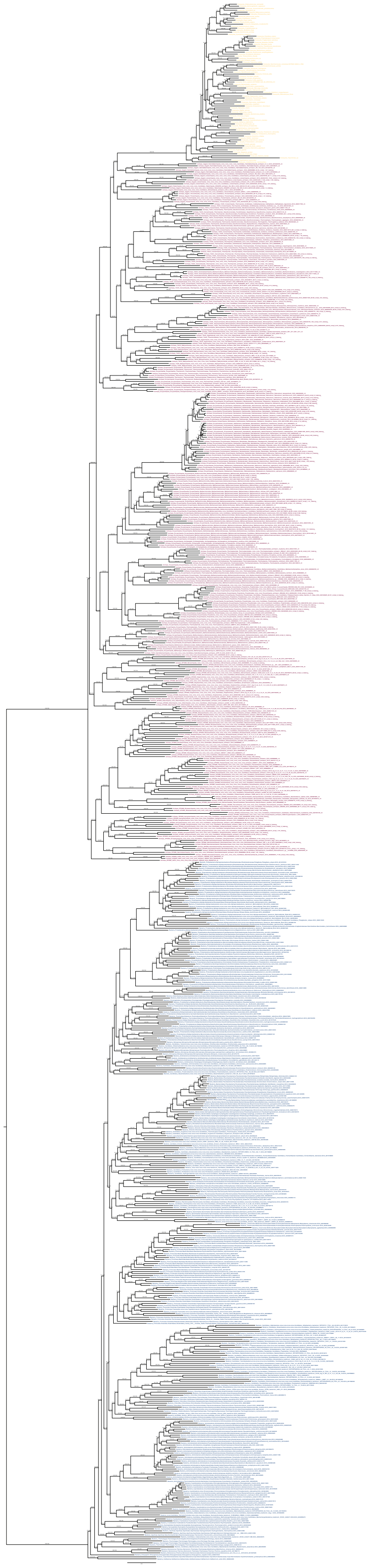

**Supplementary Figure 20**: **Maximum-likelihood concatenated species phylogeny inferred using 21 single-copy marker genes (see Methods)**. The phylogeny contains 350 Archaea (red), 350 Bacteria (blue), and 80 Eukaryotes (yellow) (780 total). Individual single-gene alignments were inferred with MAFFT L-INS-i, trimmed with BMGE v1.12 (settings: -m BLOSUM30 -h 0.55), and concatenated (see Methods). The final trimmed concatenated alignment contained 3367 positions and the phylogeny was inferred in IQ-TREE2 v2.1.2 using the LG+C20+R+F model.

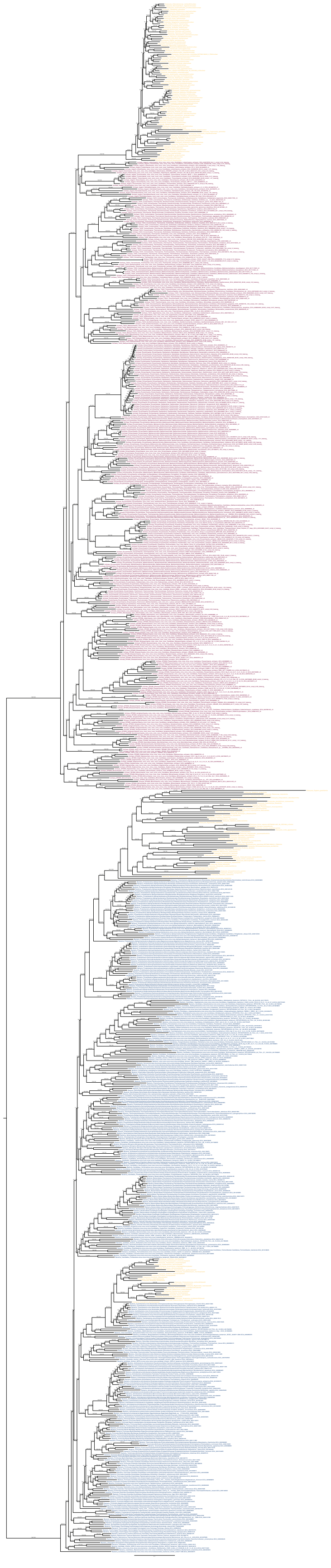

**Supplementary Figure 21**: **Maximum-likelihood concatenated species phylogeny inferred using 12 ribosomal marker genes (see Methods) and including Archaea (red), Bacteria (blue), and nuclear, mitochondrial, and plastid eukaryotic homologs (yellow)**. The phylogeny contains 350 Archaea (red), 350 Bacteria (blue), and 88 nuclear taxa, 50 mitochondrial taxa, and 25 plastid taxa (yellow). The final concatenated alignment contains

#### Supplementary Table Legends

1. **Supplementary Table 1**: NCBI and GTDB taxonomic information for 350 Archaea, 350 Bacteria, and 100 Eukaryota (800 reference taxa) included in this study.
2. **Supplementary Table 2**: Annotation table of proteins in 800 reference taxa. Protein annotations were derived from several different databases (see Methods): ArCOG = Archaeal Clusters of Orthologous Genes. KO = KEGG Orthology. CAZymes = Carbohydrate-Active enZYmes. TCDB = Transporter classification database. HydDB = Hydrogenase database. COG = NCBI Clusters of Orthologous groups.
3. **Supplementary Table 3**: List of COGs representing the F- and A/V-type ATP synthase subunits and three lipid biosynthesis genes (see Methods).
4. **Supplementary Table 4:** Count table of F- and A/V-type ATP synthase subunits represented by COG families (see Methods, Figure 1, Supplementary Figure 2, Supplementary Table 3) in 800 reference species.
5. **Supplementary Table 5**: Table summarizing presence of key metabolic organelles in 100 Eukaryotes sampled in this study. Information includes the presence of a true mitochondrion, a mitochondrion-related organelle (MRO), and/or plastid (primary, secondary, kleptoplast).
6. **Supplementary Table 6**: A summary table of the gene-tree-species-tree reconciliation results. Probability of each gene family being present at the root for different subunits, taxon sampling, substitution models, root positions and ancestral node. For each of these different conditions we infer either the presence or absence of the different subunits for LACA, LBCA and LUCA. AB = root between Archaea and Bacteria. Grac = root within Bacteria, sister to Gracilicutes. Euks = eukaryotes. C60 and C20 = the number of amino-acid replacement profiles in the mixture model for species tree estimation (see methods). Node_number = corresponding node number on the species tree. PP = presence probability. LL = loglikelihood. SUM_LL = sum of all subunit loglikelihoods for that species tree and taxon sampling. OR1 = default origination rate parameters.
7. **Supplementary Table 7**: Ancestral sequence reconstruction states table. Inferred amino acid states at each position of each node in the combined ATP synthase protein phylogeny (Figure 3A, Supplementary Figure 10, see Methods).
8. **Supplementary Table 8**: Summary of key phylogenetic results from ATP synthase protein phylogenies (Supplementary Figures 4-10). All alignments, treefiles, and other IQ-TREE output files are available in the data repository, 10.5281/zenodo.7807739)
9. **Supplementary Table 9**: Fossil calibration node assignments (i.e., calibration, node leaf/taxon assignment, age ranges, and probability masses) and braces (node leaf/taxon assignments) information applied to the Edited1, Edited2, and ATP synthase gene tree dating analyses.
10. **Supplementary Table 10**: Summarized age ranges for key speciation nodes in the species tree and ATP synthase gene tree dating analyses.
11. **Supplementary Table 11**: Taxonomic information and select metadata for 100 eukaryotic genomes/transcriptomes included in the study.
12. **Supplementary Table 12**: Summary checksheet of manual inspection of single gene trees inferred for 27 single-copy marker genes (Moody et al., 2022) [15] used to generate the concatenated species

phylogeny. Manual inspection included checks for domain monophyly, and the presence of paralogous sequences, contaminating sequences, and long-branch attraction (LBA). Marker gene distribution was used to filter genomes with poor marker gene coverage (65% threshold). All alignments, treefiles, and other IQ-TREE output files are available in the data repository, 10.5281/zenodo.7807739.

13. **Supplementary Table 13**: Count table of 21 (of 27 original) single-copy marker genes (Supplementary Table 12) present on genomes of 800 reference taxa. Includes count and percentage distribution for eukaryotic nuclear, mitochondrial, and plastid homologs which were used for marker-gene presence cutoffs (see Methods).
14. **Supplementary Table 14**: Summary of key phylogenetic results of the concatenated species phylogenies (inferred using 21 single-copy marker genes, see Supplementary Table 12) including model selection, software versions, and taxon selections. All alignments, treefiles, and other IQ-TREE output files are available in the data repository, 10.5281/zenodo.7807739
15. **Supplementary Table 15**: Results of Approximately Unbiased (AU) test to assess the statistical significance of the two concatenated species phylogenies inferred in IQ-TREE2 v2.1.2 with the LG+C20+R+F and LG+C60+R+F models (see methods). All alignments, treefiles, and other IQ-TREE output files are available in the data repository, 10.5281/zenodo.7807739
16. **Supplementary Table 16**: Summary of Ribosomal marker genes used to generate concatenated species tree containing mitochondrial and plastid eukaryotic homologs (nuclear homologs are those selected from the original concatenated species tree analysis, see Supplementary Table 12). Data includes lists of mitochondrial and plastid homologs for each representative COG marker gene corresponding to the 12 single-copy ribosomal marker genes. Mitochondrial and plastid genes are listed for each of the 12 single-copy ribosomal marker genes. All alignments, treefiles, and other IQ-TREE output files are available in the data repository, 10.5281/zenodo.7807739
17. **Supplementary Table 17**: Taxonomic mapping files used to generate the presence-absence summary plots (Figure 1, Figure 2, and Supplementary Figure 2). Archaea (n=350), Bacteria (n=350), and Eukaryotes (n=80, see Methods for filtering step) used were assessed at the clade- or species-level defined by the collapsed and long concatenated species tree, respectively (Figure 1, Supplementary Figure 2, Supplementary Figure 20). 100 eukaryotes used to assess organelle presence were clustered based on eukaryotic supergroups defined in Burki et al., 2020[67] (Figure 2).
18. **Supplementary Table 18**: Results of Approximately Unbiased (AU) test to assess the statistical significance of the topologies of the maximum-likelihood (ML) and two constrained ribosomal concatenated phylogenies used for dating (ML, Edited1, and Edited2). All alignments, treefiles, and other IQ-TREE output files are available in the data repository, 10.5281/zenodo.7807739

#### Supplementary Data File Legends

**Supplementary Data File 1:** Bracing file (json) for Edited1 species tree (see Methods; Supplementary Figures 12, 14, 15; Supplementary Table 10).

**Supplementary Data File 2:** Bracing file (json) for Edited2 species tree (see Methods; Figure 5C; Supplementary Figures 13, 16; Supplementary Table 10).

**Supplementary Data File 3:** Bracing file (json) for ATP synthase gene tree (see Methods; Figure 3A; Supplementary Figures 10, 18, 19; Supplementary Table 10).

#### Supplementary References

1. Müller, V. & Grüber, G. ATP synthases: structure, function and evolution of unique energy converters. *Cell. Mol. Life Sci.* **60**, 474–494 (2003).
2. Mulkidjanian, A. Y., Makarova, K. S., Galperin, M. Y. & Koonin, E. V. Inventing the dynamo machine: the evolution of the F-type and V-type ATPases. *Nat. Rev. Microbiol.* **5**, 892–899 (2007).
3. Hilario, E. & Gogarten, J. P. The prokaryote-to-eukaryote transition reflected in the evolution of the V/F/A-ATPase catalytic and proteolipid subunits. *J. Mol. Evol.* **46**, 703–715 (1998).
4. Stewart, A. G., Laming, E. M., Sobti, M. & Stock, D. Rotary ATPases--dynamic molecular machines. *Curr. Opin. Struct. Biol.* **25**, 40–48 (2014).
5. Forgac, M. Vacuolar ATPases: rotary proton pumps in physiology and pathophysiology. *Nat. Rev. Mol. Cell Biol.* **8**, 917–929 (2007).
6. Grüber, G., Manimekalai, M. S. S., Mayer, F. & Müller, V. ATP synthases from archaea: the beauty of a molecular motor. *Biochim. Biophys. Acta* **1837**, 940–952 (2014).
7. Kumar, A., Manimekalai, M. S. S., Balakrishna, A. M., Jeyakanthan, J. & Grüber, G. Nucleotide binding states of subunit A of the A-ATP synthase and the implication of P-loop switch in evolution. *J. Mol. Biol.* **396**, 301–320 (2010).
8. Mulkidjanian, A. Y., Galperin, M. Y., Makarova, K. S., Wolf, Y. I. & Koonin, E. V. Evolutionary primacy of sodium bioenergetics. *Biol. Direct* **3**, 13 (2008).
9. Lane, N. & Martin, W. F. The Origin of Membrane Bioenergetics. *Cell* vol. 151 1406–1416 Preprint at https://doi.org/10.1016/j.cell.2012.11.050 (2012).
10. Weiss, M. C. *et al.* The physiology and habitat of the last universal common ancestor. *Nat Microbiol* **1**, 16116 (2016).
11. Soo, R. M., Hemp, J., Parks, D. H., Fischer, W. W. & Hugenholtz, P. On the origins of oxygenic photosynthesis and aerobic respiration in Cyanobacteria. *Science* **355**, 1436–1440 (2017).

12. Shih, P. M., Hemp, J., Ward, L. M., Matzke, N. J. & Fischer, W. W. Crown group Oxyphotobacteria postdate the rise of oxygen. *Geobiology* **15**, 19–29 (2017).
13. Fournier, G. P. *et al.* The Archean origin of oxygenic photosynthesis and extant cyanobacterial lineages. *Proc. Biol. Sci.* **288**, 20210675 (2021).
14. Wang, S. & Luo, H. Dating Alphaproteobacteria evolution with eukaryotic fossils. *Nat. Commun.* **12**, 3324 (2021).
15. Moody, E. R. R. *et al.* An estimate of the deepest branches of the tree of life from ancient vertically evolving genes. *Elife* **11**, (2022).
16. Martinez-Gutierrez, C. A., Uyeda, J. C. & Aylward, F. O. A Timeline of Bacterial and Archaeal Diversification in the Ocean. Preprint at https://doi.org/10.1101/2022.10.27.514092.
17. Betts, H. C. *et al.* Integrated genomic and fossil evidence illuminates life's early evolution and eukaryote origin. *Nat Ecol Evol* **2**, 1556–1562 (2018).
18. Martinez-Gutierrez, C. A. & Aylward, F. O. Phylogenetic Signal, Congruence, and Uncertainty across Bacteria and Archaea. *Mol. Biol. Evol.* **38**, 5514–5527 (2021).
19. Ward, L. M. & Shih, P. M. The evolution and productivity of carbon fixation pathways in response to changes in oxygen concentration over geological time. *Free Radic. Biol. Med.* **140**, 188–199 (2019).
20. Soo, R. M., Hemp, J. & Hugenholtz, P. Evolution of photosynthesis and aerobic respiration in the cyanobacteria. *Free Radic. Biol. Med.* **140**, 200–205 (2019).
21. Boussau, B., Karlberg, E. O., Frank, a. C., Legault, B.-A. & Andersson, S. G. E. Computational inference of scenarios for alpha-proteobacterial genome evolution. *Proc. Natl. Acad. Sci. U. S. A.* **101**, 9722–9727 (2004).
22. Liu, Y. *et al.* Expanded diversity of Asgard archaea and their relationships with eukaryotes. *Nature* **593**, 553–557 (2021).
23. Cunha, V. D., Da Cunha, V., Gaïa, M. & Forterre, P. The expanding Asgard archaea and their elusive

relationships with Eukarya. *mLife* vol. 1 3–12 Preprint at https://doi.org/10.1002/mlf2.12012 (2022).

24. Martijn, J., Vosseberg, J., Guy, L., Offre, P. & Ettema, T. J. G. Deep mitochondrial origin outside the sampled alphaproteobacteria. *Nature* **557**, 101–105 (2018).
25. Martin, W. F., Garg, S. & Zimorski, V. Endosymbiotic theories for eukaryote origin. *Philos. Trans. R. Soc. Lond. B Biol. Sci.* **370**, 20140330 (2015).
26. Muñoz-Gómez, S. A. *et al.* Site-and-branch-heterogeneous analyses of an expanded dataset favour mitochondria as sister to known Alphaproteobacteria. *Nat Ecol Evol* **6**, 253–262 (2022).
27. Nelson, D. R. 178043: lithic-quartz sandstone, Strelley Pool. in *Compilation of geochronology data, June 2007 update* (Geological Survey of Western Australia, 2005b).
28. Hanan, B. B. & Tilton, G. R. 60025: relict of primitive lunar crust? *Earth and Planetary Science Letters* vol. 84 15–21 Preprint at https://doi.org/10.1016/0012-821x(87)90171-3 (1987).
29. Barboni, M. *et al.* Early formation of the Moon 4.51 billion years ago. *Sci Adv* **3**, e1602365 (2017).
30. Satkoski, A. M., Beukes, N. J., Li, W., Beard, B. L. & Johnson, C. M. A redox-stratified ocean 3.2 billion years ago. *Earth and Planetary Science Letters* vol. 430 43–53 Preprint at https://doi.org/10.1016/j.epsl.2015.08.007 (2015).
31. Bosak, T., Knoll, A. H. & Petroff, A. P. The Meaning of Stromatolites. *Annual Review of Earth and Planetary Sciences* vol. 41 21–44 Preprint at https://doi.org/10.1146/annurev-earth-042711-105327 (2013).
32. Cairns-Smith, A. G. Precambrian solution photochemistry, inverse segregation, and banded iron formations. *Nature* vol. 276 807–808 Preprint at https://doi.org/10.1038/276807a0 (1978).
33. Byerly, G. R., Kröner, A., Lowe, D. R., Todt, W. & Walsh, M. M. Prolonged magmatism and time constraints for sediment deposition in the early Archean Barberton greenstone belt: evidence from the Upper Onverwacht and Fig Tree groups. *Precambrian Research* vol. 78 125–138 Preprint at

https://doi.org/10.1016/0301-9268(95)00073-9 (1996).

34. Kamo, S. L. & Davis, D. W. Reassessment of Archean crustal development in the Barberton Mountain Land, South Africa, based on U-Pb dating. *Tectonics* vol. 13 167–192 Preprint at https://doi.org/10.1029/93tc02254 (1994).

35. Demoulin, C. F. *et al.* Cyanobacteria evolution: Insight from the fossil record. *Free Radic. Biol. Med.* **140**, 206–223 (2019).

36. Hodgskiss, M. S. W. *et al.* New insights on the Orosirian carbon cycle, early Cyanobacteria, and the assembly of Laurentia from the Paleoproterozoic Belcher Group. *Earth and Planetary Science Letters* vol. 520 141–152 Preprint at https://doi.org/10.1016/j.epsl.2019.05.023 (2019).

37. Gardiner, N. J., Wacey, D., Kirkland, C. L., Johnson, T. E. & Jeon, H. Zircon U–Pb, Lu–Hf and O isotopes from the 3414 Ma Strelley Pool Formation, East Pilbara Terrane, and the Palaeoarchaean emergence of a cryptic cratonic core. *Precambrian Research* vol. 321 64–84 Preprint at https://doi.org/10.1016/j.precamres.2018.11.023 (2019).

38. Brocks, J. J. & Schaeffer, P. Okenane, a biomarker for purple sulfur bacteria (Chromatiaceae), and other new carotenoid derivatives from the 1640Ma Barney Creek Formation. *Geochimica et Cosmochimica Acta* vol. 72 1396–1414 Preprint at https://doi.org/10.1016/j.gca.2007.12.006 (2008).

39. Munson, T. J., Denyszyn, S. W., Simmons, J. M. & Kunzmann, M. A 1642 Ma age for the Fraynes Formation, Birrindudu Basin, confirms correlation with the economically significant Barney Creek Formation, McArthur Basin, Northern Territory. *Australian Journal of Earth Sciences* vol. 67 321–330 Preprint at https://doi.org/10.1080/08120099.2020.1669708 (2020).

40. Page, R. W. & Sweet, I. P. Geochronology of basin phases in the western Mt Isa Inlier, and correlation with the McArthur Basin∗. *Australian Journal of Earth Sciences* vol. 45 219–232 Preprint at https://doi.org/10.1080/08120099808728383 (1998).

41. Li, H. *et al.* Recent advances in the study of the Mesoproterozoic geochronology in the North China Craton. *Journal of Asian Earth Sciences* vol. 72 216–227 Preprint at https://doi.org/10.1016/j.jseaes.2013.02.020 (2013).

42. Butterfield, N. J. *Bangiomorpha pubescens*n. gen., n. sp.: implications for the evolution of sex, multicellularity, and the Mesoproterozoic/Neoproterozoic radiation of eukaryotes. *Paleobiology* vol. 26 386–404 Preprint at https://doi.org/10.1666/0094-8373(2000)026<0386:bpngns>2.0.co;2 (2000).

43. Gibson, T. M. *et al.* Precise age of Bangiomorpha pubescens dates the origin of eukaryotic photosynthesis. *Geology* vol. 46 135–138 Preprint at https://doi.org/10.1130/g39829.1 (2018).

44. Agić, H. Origin and Early Evolution of the Eukaryotes: Perspectives from the Fossil Record. *Prebiotic Chemistry and the Origin of Life* 255–289 Preprint at https://doi.org/10.1007/978-3-030-81039-9_11 (2021).

45. Fralick, P., Davis, D. W. & Kissin, S. A. The age of the Gunflint Formation, Ontario, Canada: single zircon UPb age determinations from reworked volcanic ash. *Canadian Journal of Earth Sciences* vol. 39 1085–1091 Preprint at https://doi.org/10.1139/e02-028 (2002).

46. Loron, C. C., Rainbird, R. H., Turner, E. C., Wilder Greenman, J. & Javaux, E. J. Organic-walled microfossils from the late Mesoproterozoic to early Neoproterozoic lower Shaler Supergroup (Arctic Canada): Diversity and biostratigraphic significance. *Precambrian Research* vol. 321 349–374 Preprint at https://doi.org/10.1016/j.precamres.2018.12.024 (2019).

47. Loron, C. C. *et al.* Early fungi from the Proterozoic era in Arctic Canada. *Nature* **570**, 232–235 (2019).

48. van Acken, D., Thomson, D., Rainbird, R. H. & Creaser, R. A. Constraining the depositional history of the Neoproterozoic Shaler Supergroup, Amundsen Basin, NW Canada: Rhenium-osmium dating of black shales from the Wynniatt and Boot Inlet Formations. *Precambrian Research* vol. 236 124–131

Preprint at https://doi.org/10.1016/j.precamres.2013.07.012 (2013).

49. Redecker, D., Kodner, R. & Graham, L. E. Glomalean Fungi from the Ordovician. *Science* vol. 289 1920–1921 Preprint at https://doi.org/10.1126/science.289.5486.1920 (2000).

50. Redecker, D., Kodner, R. & Graham, L. E. Palaeoglonius grayi from the Ordovician. *Mycotaxon - Ithaca Ny-* (2002).

51. Goldman, D. *et al.* The Ordovician Period. *Geologic Time Scale 2020* 631–694 Preprint at https://doi.org/10.1016/b978-0-12-824360-2.00020-6 (2020).

52. Liu, A. G., McIlroy, D., Matthews, J. J. & Brasier, M. D. A new assemblage of juvenile Ediacaran fronds from the Drook Formation, Newfoundland. *Journal of the Geological Society* vol. 169 395–403 Preprint at https://doi.org/10.1144/0016-76492011-094 (2012).

53. Dunn, F. S. *et al.* The developmental biology of *Charnia* and the eumetazoan affinity of the Ediacaran rangeomorphs. *Science Advances* vol. 7 Preprint at https://doi.org/10.1126/sciadv.abe0291 (2021).

54. Matthews, J. J. *et al.* A CHRONOSTRATIGRAPHIC FRAMEWORK FOR THE RISE OF THE EDIACARAN MACROBIOTA: NEW CONSTRAINTS FROM MISTAKEN POINT ECOLOGICAL RESERVE, NEWFOUNDLAND. *Geological Society of America Abstracts with Programs* Preprint at https://doi.org/10.1130/abs/2020am-352772 (2020).

55. Yang, C. *et al.* The tempo of Ediacaran evolution. *Sci Adv* **7**, eabi9643 (2021).

56. Dunn, F. S. *et al.* A crown-group cnidarian from the Ediacaran of Charnwood Forest, UK. *Nat Ecol Evol* **6**, 1095–1104 (2022).

57. Wilby, P. R., Carney, J. N. & Howe, M. P. A. A rich Ediacaran assemblage from eastern Avalonia: Evidence of early widespread diversity in the deep ocean. *Geology* vol. 39 655–658 Preprint at https://doi.org/10.1130/g31890.1 (2011).

58. Yin, Z. *et al.* Diverse and complex developmental mechanisms of early Ediacaran embryo-like fossils

from the Weng'an Biota, southwest China. *Philos. Trans. R. Soc. Lond. B Biol. Sci.* **377**, 20210032 (2022).

59. Steiner, M., Li, G., Qian, Y., Zhu, M. & Erdtmann, B.-D. Neoproterozoic to Early Cambrian small shelly fossil assemblages and a revised biostratigraphic correlation of the Yangtze Platform (China). *Palaeogeography, Palaeoclimatology, Palaeoecology* vol. 254 67–99 Preprint at https://doi.org/10.1016/j.palaeo.2007.03.046 (2007).
60. Runnegar, B. Muscle scars, shell form and torsion in Cambrian and Ordovician univalved molluscs. *Lethaia* vol. 14 311–322 Preprint at https://doi.org/10.1111/j.1502-3931.1981.tb01104.x (1981).
61. Maloof, A. C. *et al.* Constraints on early Cambrian carbon cycling from the duration of the Nemakit-Daldynian–Tommotian boundary δ13C shift, Morocco. *Geology* vol. 38 623–626 Preprint at https://doi.org/10.1130/g30726.1 (2010).
62. Hughes, N. F. & McDougall, A. B. Barremian-Aptian angiospermid pollen records from southern England. *Review of Palaeobotany and Palynology* vol. 65 145–151 Preprint at https://doi.org/10.1016/0034-6667(90)90065-q (1990).
63. Clarke, J. T., Warnock, R. C. M. & Donoghue, P. C. J. Establishing a time-scale for plant evolution. *New Phytol.* **192**, 266–301 (2011).
64. Judd, W. S. & Olmstead, R. G. A survey of tricolpate (eudicot) phylogenetic relationships. *Am. J. Bot.* **91**, 1627–1644 (2004).
65. Ogg, J. G. Geomagnetic Polarity Time Scale. *Geologic Time Scale 2020* 159–192 Preprint at https://doi.org/10.1016/b978-0-12-824360-2.00005-x (2020).
66. Vigran, J. O., Mangerud, G., Mørk, A., Bugge, T. & Weitschat, W. Biostratigraphy and sequence stratigraphy of the lower and middle Triassic deposits from the Svalis Dome, Central Barents Sea, Norway. *Palynology* vol. 22 89–141 Preprint at https://doi.org/10.1080/01916122.1998.9989505 (1998).

67. Burki, F., Roger, A. J., Brown, M. W. & Simpson, A. G. B. The New Tree of Eukaryotes. *Trends Ecol. Evol.* **35**, 43–55 (2020).

68. Nirody, J. A., Budin, I. & Rangamani, P. ATP synthase: Evolution, energetics, and membrane interactions. *J. Gen. Physiol.* **152**, (2020).

69. Tatusov, R. L., Koonin, E. V. & Lipman, D. J. A genomic perspective on protein families. *Science* **278**, 631–637 (1997).

70. Galperin, M. Y., Kristensen, D. M., Makarova, K. S., Wolf, Y. I. & Koonin, E. V. Microbial genome analysis: the COG approach. *Brief. Bioinform.* **20**, 1063–1070 (2019).

71. Galperin, M. Y. *et al.* COG database update: focus on microbial diversity, model organisms, and widespread pathogens. *Nucleic Acids Res.* **49**, D274–D281 (2021).
